# Clonal autoantibodies identify microbial antigen as trigger of autoreactive B cells in systemic sclerosis

**DOI:** 10.64898/2026.03.17.712484

**Authors:** Sam Neppelenbroek, Sophie I.E. Liem, Theo van Laar, Eva M. Hoekstra, Corrie M. Wortel, E.W. Nivine Levarht, Cynthia M. Fehres, Nynke H. Dekker, Jeska K. de Vries-Bouwstra, René E.M. Toes, Hans U. Scherer

## Abstract

**Objectives:** Transformative observations demonstrate unprecedented success of B cell-depleting interventions in many human autoimmune diseases, calling for a deeper understanding of the triggers leading to B cell-mediated autoimmunity and its perpetuation in human disease. Here, we investigated whether the autoreactive B cell response targeting human topoisomerase 1 (TOP1), a hallmark of systemic sclerosis, could cross-react with TOP1 of microbial origin.

**Methods:** Homologies between human and microbial TOP1 were analyzed using Foldseek. TOP1-reactive monoclonal antibodies from patient-derived, human TOP1-reactive B cell receptors were generated and assessed for reactivity against human TOP1 and TOP1 from a prototypic yeast, *Saccharomyces cerevisiae* (*S. cerevisiae*). Reactivity of polyclonal serum IgG from anti-TOP1 autoantibody (ATA)^+^, anti-centromere autoantibody (ACA)^+^ SSc patients and healthy donors (HDs) was tested. Finally, B cell lines were generated expressing human ATA to study B cell activation upon antigenic stimulation.

**Results:** Structural homologues of human TOP1 were found in many microbes, particularly in fungi. Taking TOP1 from *S. cerevisiae* as a prototype, microbial TOP1 was recognized by polyclonal patient IgG and by several monoclonal ATAs. Importantly, *S. cerevisiae* TOP1 also activated B cells expressing a patient-derived, human TOP1-reactive B cell receptor. Patients affected by interstitial lung disease most frequently showed recognition of microbial TOP1.

**Conclusions:** These findings identify fungi as potential drivers of immune dysregulation in human autoimmunity, specifically in SSc, highlighting microbial antigen cross-reactive cells as important therapeutic targets. Moreover, these data provide first functional evidence for a breach of B cell tolerance against human TOP1 triggered by cross-reactivity to fungal TOP1.

## Introduction

Systemic sclerosis (SSc) is a chronic autoimmune disease (AID) in which vasculopathy and fibrosis affect the function of internal organs and the skin. The clinical disease course varies strongly between patients, particularly in terms of organ involvement and rate of disease progression. This heterogeneity, together with a lack of effective and curative treatments, makes the disease unpredictable and complicates its clinical management. Therefore, it is crucial to understand the mechanisms driving the disease to reduce SSc-related morbidity and mortality.

Anti-nuclear autoantibodies hallmark SSc and define its clinical phenotype and course of disease (*1*). As is the case for other AIDs, striking clinical improvements have been observed in severely ill patients upon B cell depletion, for example with rituximab or anti-CD19 chimeric antigen-receptor (CAR)-T cells (*2–4*). These observations indicate a pivotal role of (autoreactive) B cells in disease initiation and progression. How such B cells are generated, activated and maintained, however, remains unclear. The human nuclear protein topoisomerase 1 (TOP1) is the most prevalent target of autoreactive B cells in patients with diffuse cutaneous SSc (*5, 6*). Anti-TOP1 autoantibodies (ATAs) associate with fibrosis of skin and lungs as well as mortality (*6–8*). Recently, we also found an association between the activity of the B cell response generating ATA and disease progression as well as pulmonary fibrosis (*9, 10*). Hence, it is conceivable that active, TOP1-reactive B cells contribute directly or indirectly to the inflammatory and fibrotic processes in affected organs. Despite these indications, the triggers driving the initiation and activation of TOP1-reactive B cells are undefined.

Microbes have been implicated in the break of B cell tolerance towards self-antigens in several autoimmune diseases (*11, 12*). Antigens derived from microbes can trigger the B cell receptor (BCR) of autoreactive B cells by structurally resembling the autoantigen, a concept known as molecular mimicry (*12*). For example, commensal bacteria expressing orthologs of the human Ro60 protein were identified as targets of autoreactive B cells in systemic lupus erythematosus (*13*). By analogy, we considered that antibodies and BCRs reactive to human TOP1 could also recognize TOP1 from microbial species. SSc patients could be exposed to such microbial TOP1 antigens in affected organs, especially the lungs, the skin or the gut (*14*). Also, the majority of ATA-IgG^+^ patients harbor ATA-IgA, the predominant isotype in mucosal immune responses (*9, 15*). Nevertheless, there are only limited studies on the presence and abundance of microbes in affected SSc tissues (*16–18*). Moreover, it is unknown whether ATAs can recognize microbial antigens. Elucidating this possibility would be relevant as it may indicate that microbial TOP1 contributes to the breakdown of immune tolerance to human TOP1 in systemic sclerosis, as well as the activation of TOP1-reactive B cells in progressive disease. Based on the above, we searched for structural homologies between human and microbial TOP1. We studied the recognition of microbial TOP1 by patient-derived ATAs and the ability of microbial TOP1 to activate ATA-expressing B cells. We found that TOP1 from *Saccharomyces cerevisiae* (*S. cerevisiae*), a microbial TOP1 sharing structural similarities with human TOP1, is recognized by selected human ATAs and capable of activating ATA-expressing human B cells. The recognition of this microbial TOP1 was enhanced in severe ATA^+^ SSc patients and in patients with clinically relevant interstitial lung disease (ILD). Together, these results provide evidence that microbial TOP1 could be involved in the initiation and activation of TOP1-reactive B cells in patients and suggest that this process could drive severe SSc.

## Materials and Methods

The experimental procedures for the computational analysis of structural homology between human and microbial TOP1 using FoldSeek, recombinant production and purification of human and *S. cerevisiae* TOP1, generation of patient-derived ATA mAbs, gel electrophoresis, anti-human TOP1-IgG and anti-S. cerevisiae TOP1-IgG ELISA, generation of Ramos B cell lines expressing TOP1-reactive IgG BCRs and stimulation of ATA-expressing Ramos B cell lines with human and S. cerevisiae TOP1 are provided with the details on the study approval and statistical analysis in the supplementary method section.

### Study design

Biological samples from SSc patients were obtained at the Department of Rheumatology at Leiden University Medical Center (LUMC, The Netherlands). All SSc patients fulfilled the 2013 American College of Rheumatology (ACR)/European League Against Rheumatism (EULAR) classification criteria for SSc (*19*). Patients on B cell-depleting therapies and patients with a history of hematopoietic stem cell transplantation were excluded.

Cohort 1 - Plasma samples from ATA^+^ SSc patients (n=22), ACA^+^ SSc patients (n=20) and HDs (n=16) included in a previous study (Table 1) (*10*). Patients were recruited irrespective of disease state or organ involvement.

**Table 1.** Characteristics of SSc patients and healthy donors included in the cohort 1.

|  | ATA<sup>+</sup> SSc patients<br>(n=22) | ACA<sup>+</sup> SSc patients<br>(n=20) | Healthy donors<br>(n=16) |
| --- | --- | --- | --- |
| Age, years, mean (SD) | 52 (16) | 62 (11) | 37 (12) |
| Female, n (%) | 12 (55%) | 18 (90%) | 12 (75%) |
| Duration since first NRP symptom, months, median (IQR) | 63 (39-102) | 120 (95-221) |  |
| Diffuse cutaneous SSc, n (%) | 7 (32%) | 0 (0%) |  |
| mRSS, median (IQR) | 5 (2-7) | 3 (0-8) |  |
| ILD on HRCT, n (%) | 14 (64%) | 0 (0%) |  |
| Clinically relevant ILD, n (%) | 2 (9%) | 0 (0%) |  |
| Pulmonary hypertension, n (%) | 0 (0%) | 2 (10%) |  |
| Digital ulcers, n (%) | 2 (9%) | 0 (0%) |  |
| Current immunosuppressive use, n (%) | 12 (55%) | 6 (30%) |  |

Patients and controls were included in the context of a previous study in which patient selection and characteristics are described in detail (Wortel et al. RMD Open, 2023(10)). Clinical characteristics represent data collected at moment of sampling. SD = standard deviation, IQR = interquartile range, NRP = non-Raynaud's phenomenon, mRSS = modified Rodnan skin score, ILD = interstitial lung disease, HRCT = high-resolution computer tomography scan, Clinically relevant ILD = presence of ILD on HRCT combined with a forced vital capacity below 80% of the predicted value.

Cohort 2 - To replicate findings in cohort 1 and to explore possible associations with disease severity, serum samples from ATA^+^ SSc patients (n=39) were selected from the CCISS cohort (*20*). Selection was performed by two independent physicians (JKdVB, SIEL), experienced in the field of SSc, with the aim to identify patients at both ends of the SSc disease severity spectrum: severe ATA^+^ SSc (n=18) versus mild ATA^+^ SSc (n=21) (Table 2). This approach was chosen because currently available disease activity indices have significant limitations for hypothesis-generating studies. For example, they often fail to distinguish between disease activity and irreversible damage, and inadequately capture involvement of specific organ systems, such as the gastrointestinal tract (*21*). Patients with severe SSc were selected by chart review and clinical expertise taking into account all available data over time including involvement of skin, lung, heart, kidney, gastrointestinal tract and vascular system. Next, patients with mild disease matched with severe patients for age and disease duration. Both physicians had to agree on classification of a patient as being mild or severe; if not, the patient was excluded from cohort 2. Samples collected at time of inclusion in the CCISS cohort were used. Serum samples from age- and sex-matched ACA^+^ SSc patients (n=10) and HDs (n=10) obtained from the CCISS cohort and Leiden University Medical Center Voluntary Donor Service, respectively, were used as controls (Table 2).

**Table 2.** Characteristics of SSc patients and healthy donors included in cohort 2.

|  | ATA<sup>+</sup> SSc patients<br>(n=39) |  |  |  | ACA<sup>+</sup> SSc patients<br>(n=10) | Healthy donors<br>(n=10) |
| --- | --- | --- | --- | --- | --- | --- |
|  | Baseline |  | Follow-up |  |  |  |
|  | Mild<br>(n=21) | Severe<br>(n=18) | Mild<br>(n=21) | Severe<br>(n=18) |  |  |
| Age, years, mean (SD) | 48 (16) | 50 (14) | - | - | 49 | 51 (3) |
| Female, n (%) | 16 (76%) | 10 (56%) | - | - | 7 (70%) | 7 (70%) |
| Duration since first NRP symptom, months,<br>median (IQR) | 13 (6-64) | 12 (4-38) | - | - | 109 (76-193) |  |
| Baseline to follow-up, months, median (IQR) | - | - | 40 (20-79) | 30 (13-55) | - |  |
| Diffuse cutaneous SSc, n (%) | 1 (5%) | 9 (50%) | 1 (5%) | 12 (67%) | 1 (10%) |  |
| mRSS, median (IQR) | 4 (2-7) | 4 (3-23) | 4 (2-6) | 9 (4-20) | 4 (2-5) |  |
| ILD on HRCT, n (%) | 10 (48%) | 8 (44%) | 12 (57%) | 14 (78%) | 0 (0%) |  |
| Clinically relevant ILD, n (%) | 1 (5%) | 3 (17%) | 1 (5%) | 5 (28%) | 0 (0%) |  |
| Pulmonary hypertension, n (%) | 0 (0%) | 0 (0%) | 0 (0%) | 0 (0%) | 0 (0%) |  |
| Digital ulcers, n (%) | 3 (14%) | 1 (6%) | 1 (5%) | 5 (28%) | 2 (20%) |  |
| ASCT between baseline & follow-up, n (%) | - | - | 0 | 3 (17%) | - |  |
| Current immunosuppressive use, n (%) | 6 (29%) | 6 (33%) | 8 (38%) | 11 (61%) | 2 (20%) |  |
| Azathioprine, n | 1 | 0 | 1 | 0 | 0 |  |
| Hydroxychloroquine, n | 1 | 0 | 1 | 0 | 1 |  |
| Corticosteroids > 5mg/day, n | 0 | 4 | 0 | 2 | 0 |  |
| Methotrexate, n | 4 | 1 | 6 | 3 | 2 |  |
| Cyclophosphamide, n | 0 | 0 | 0 | 3 | 0 |  |
| Mycophenolic acid, n | 0 | 1 | 0 | 3 | 0 |  |
| Other (infliximab, tocilizumab), n | 0 | 1 | 0 | 2 | 0 |  |

Baseline characteristics represent clinical data collected at cohort entry. SD = standard deviation, IQR = interquartile range, NRP = non-Raynaud's phenomenon, mRSS = modified Rodnan skin score, ILD = interstitial lung disease, HRCT = high-resolution computer tomography scan, clinically relevant ILD = presence of ILD on HRCT combined with a forced vital capacity below 80% of the predicted value, ASCT = autologous stem cell transplantation.

Cohort 3 - To replicate the clinical associations found in cohort 2, serum samples from all ATA^+^ SSc patients included in the CCISS cohort (n=163) until 31^st^ of December, 2023, which fulfilled the ACR 2013 criteria for SSc were used (*19, 20*). Samples collected at time of inclusion in the CCISS cohort were used. Serum samples from age- and sex-matched ACA^+^ SSc patients (n=50) and HDs (n=30) obtained from the CCISS cohort and Leiden University Medical Center Voluntary Donor Service (LuVDS), respectively, were used as controls (Table 3).

**Table 3.**
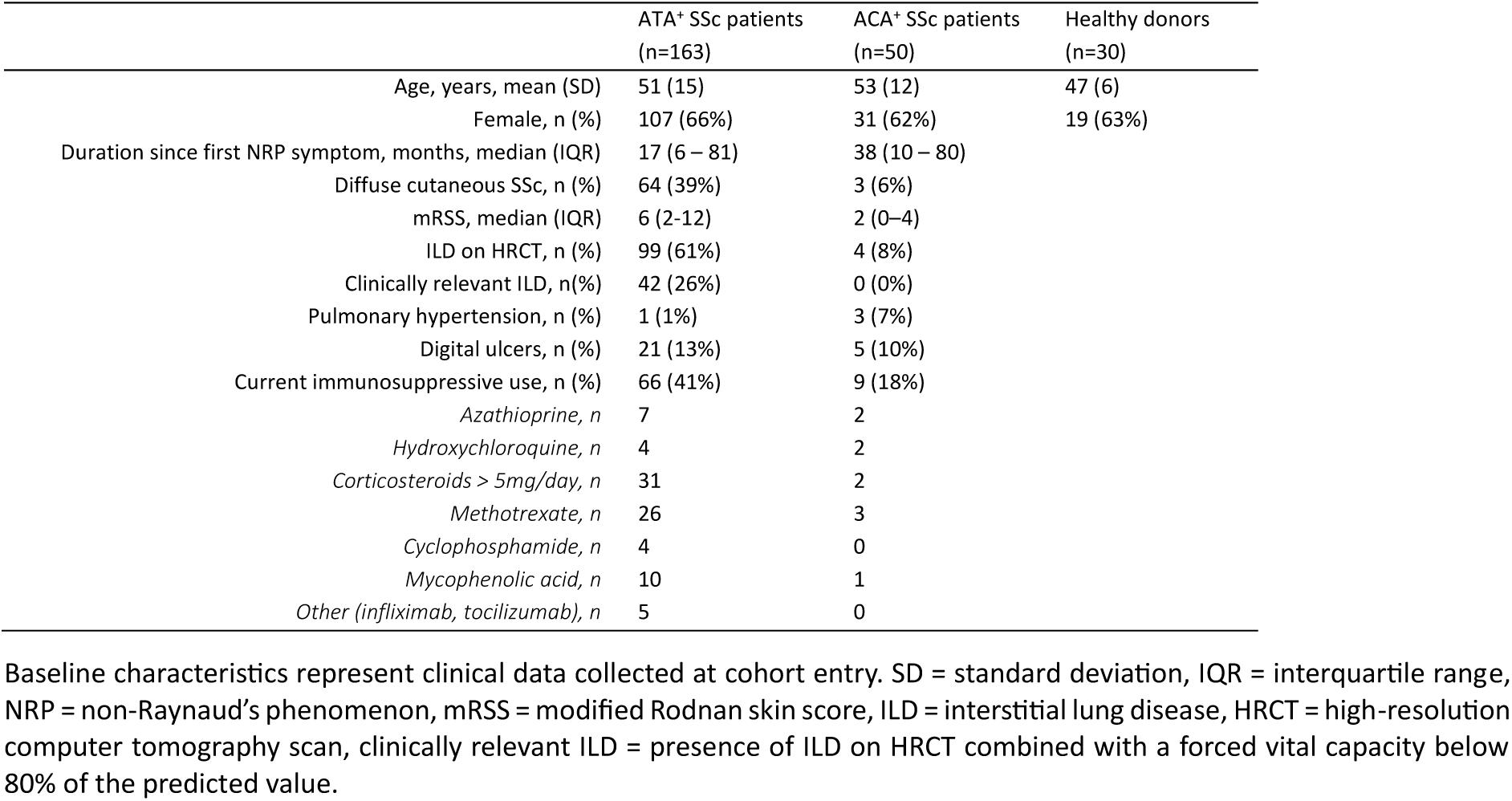
Baseline characteristics of SSc patients and healthy donors included in cohort 3.

Clinical data acquisition: All included patients participate in the ongoing prospective CCISS cohort in which clinical variables of SSc patients are monitored in a structural follow-up program which included an annual comprehensive assessment of organ involvement (*20*). ILD was diagnosed based on the presence of ground glass opacities or interstitial fibrosis on high-resolution computed tomography (HRCT) scan of the thorax. Clinically relevant ILD was defined as presence of ILD on HRCT combined with a FVC below 80% of predicted. Skin involvement was evaluated by assessing the modified Rodnan Skin Score (MRSS) (*22*). Pattern of skin involvement was used to categorize patients into diffuse cutaneous, limited cutaneous or non-cutaneous disease subsets.

## Results

### TOP1 is structurally conserved in microbes, particularly in fungi

To identify microbial proteins that could be recognized by human ATAs, we performed a homology search for proteins sharing structural similarities with human TOP1. We used Foldseek with a crystal structure of human TOP1 (1EJ9, Protein Data Bank) as input (*23, 24*). Foldseek searches databases for proteins or protein fragments with resolved or predicted tertiary protein structures that show structural similarities with a protein of interest (*24*). Like other crystal structures of human TOP1, 1EJ9 does not contain the poorly conserved N-terminal and linker domains (*23*). A taxonomic filter was applied to only identify proteins of fungal, bacterial or viral origin. The search identified 145 fungal, 273 bacterial and 3 viral hits (Fig. 1A). Most hits were TOP1 proteins in the respective microbial compartments (Table S1-S3). Fungal hits showed the highest template modeling scores and lowest expect values (E-values) (Fig. 1A). Interestingly, strong structural similarities were observed between human TOP1 and TOP1 from, for example, *S. cerevisiae, Pseudomonas aeruginosa* and *Variola* virus, despite relatively low sequence identities (41.2%, 17.3% and 17.2%, respectively) (Fig. 1B-D). Together, these findings provided rational to explore cross-recognition of microbial TOP1 molecules by human ATAs.

**Figure 1.**
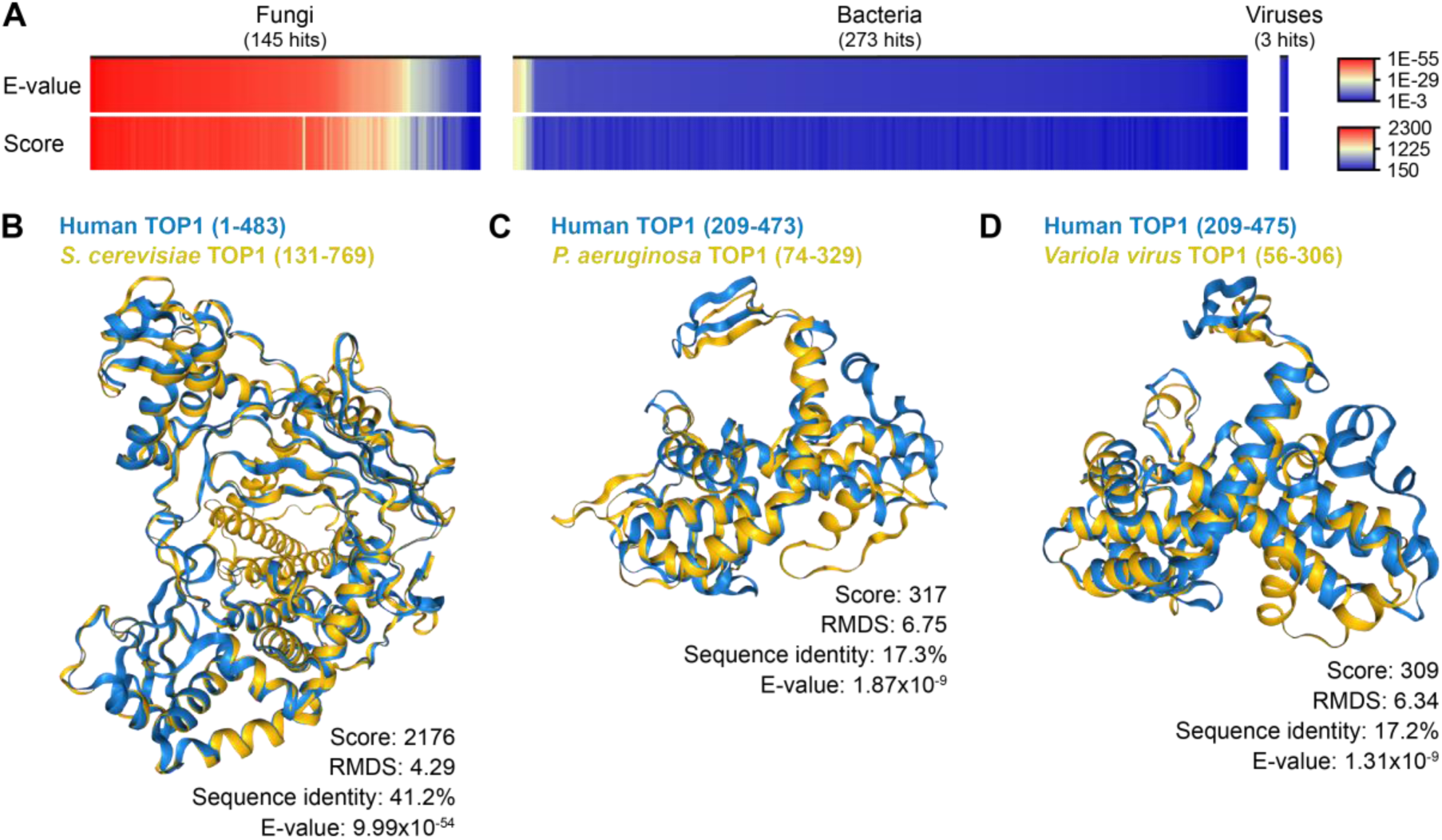
Homology search for the identification of microbial proteins sharing structural similarities with human TOP1. (A) Overview of microbial proteins having structural homology with human TOP1 as predicted by Foldseek. Homology search was based on a crystal structure of human TOP1 (1EJ9, Protein Data Bank). A taxonomic filter was used to specifically identify fungal, bacterial and viral proteins. Hits were ranked based on E-values. (B-D) Structural homology as predicted by Foldseek between human TOP1 (in blue) and TOP1 from *S. cerevisiae* (AlphaFold/Proteome (v4) database) (B)*, P. aeruginosa* (AlphaFold/Proteome (v4) database) (C) and *Variola* virus (PDB100 database (20240101)) (D) (all in yellow). Amino acid positions of predicted structurally overlapping parts are indicated between brackets. Score = structural bit score. RMDS = root mean square deviation. Expect values = E-values.

### Circulating antibodies from ATA^+^ SSc patients recognize TOP1 from S. cerevisiae

We selected TOP1 from *S. cerevisiae* as exemplary model antigen to test the hypothesis. TOP1 from this fungus was amongst the predicted hits with the strongest structural similarity with human TOP1 and contains some homologous stretches of amino acids (Table S1, Fig. S1). In addition, *S. cerevisiae* is abundantly present at mucosal sites, including the lungs (*25–27*). Immunological exposure to *S. cerevisiae* can induce anti-*S. cerevisiae* antibodies (ASCAs) which are associated with Crohn’s disease but have also been detected in healthy donors (HDs) and SSc patients (*28–30*). Therefore, we recombinantly produced and subsequently purified human and *S. cerevisiae* TOP1 (Fig. S2). As expected, human TOP1 was recognized specifically by ATA^+^ SSc patients but not by anti-centromere autoantibody (ACA)^+^ SSc patients and HDs (cohort 1, Fig. 2A, Fig. S3A, Table 1). Interestingly, a subset of ATA^+^ SSc patients (45%) showed also reactivity to *S. cerevisiae* TOP1 (Fig. 2B, Fig. S3B). This reactivity was not observed in plasma from ACA^+^ SSc patients and HDs (Fig. 2B, Fig. S3B). Recognition of human and *S. cerevisiae* TOP1 weakly correlated (r_s_=0.35, p=0.12), with strong variation in the recognition of *S. cerevisiae* TOP1 among ATA^+^ SSc patients with comparable anti-human TOP1-IgG levels (Fig. 2C, Fig. S3C).

**Figure 2.**
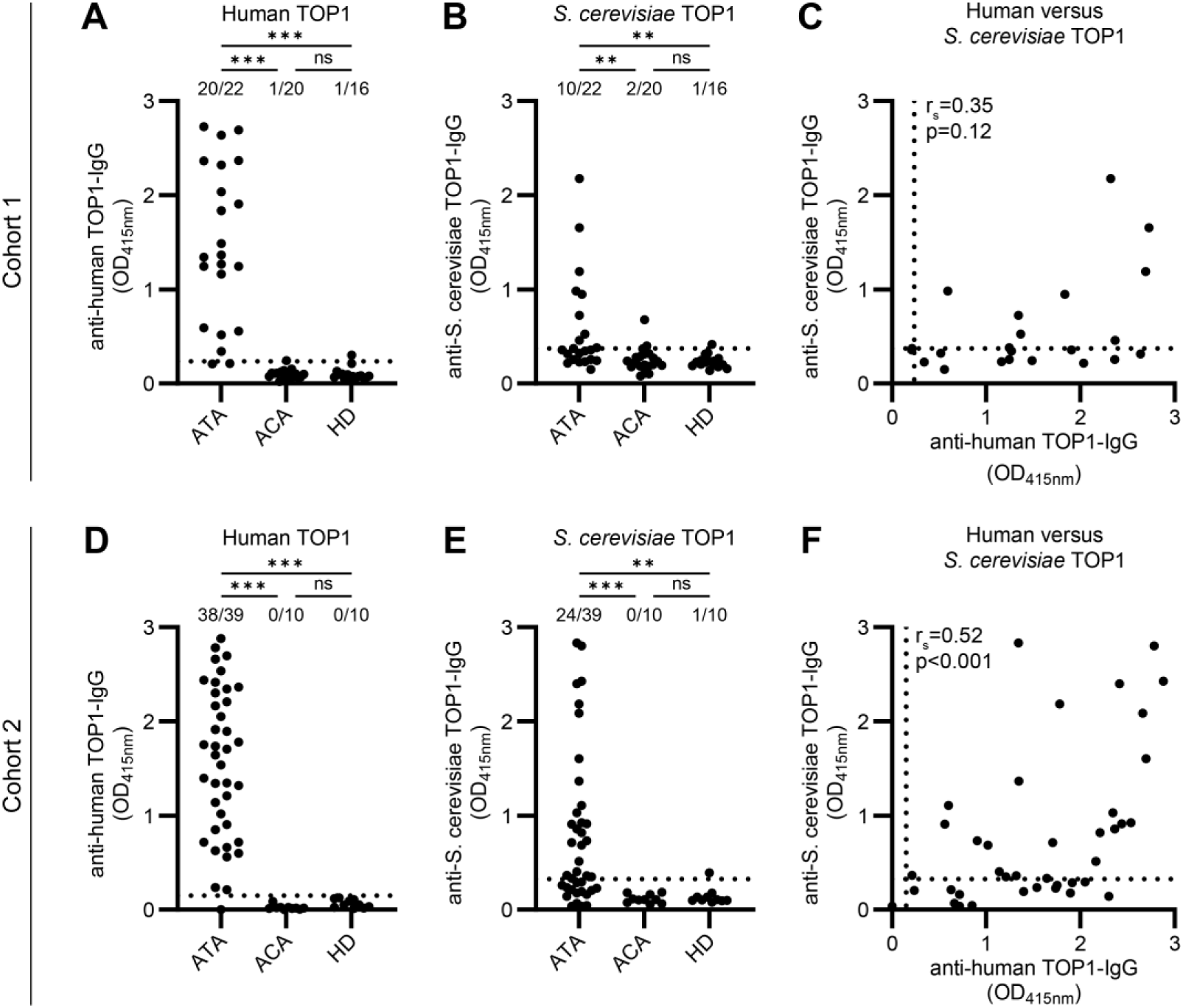
Recognition of *S. cerevisiae* TOP1 by circulating IgG of ATA^+^ SSc patients. (A-B) Reactivity of IgG in plasma of ATA^+^ SSc patients (n=22), ACA^+^ SSc patients (n=20) and healthy donors (HD)(n=16) included in cohort 1 towards human (A) and *S. cerevisiae* (B) TOP1. (C) Correlation between the recognition of human and *S. cerevisiae* TOP1 by ATA^+^ SSc patients in cohort 1. (D-E) Binding of human (D) and *S. cerevisiae* (E) TOP1 by IgG in serum of ATA^+^ SSc patients (n=39), ACA^+^ SSc patients (n=10) and healthy donors (HD)(n=10) included in cohort 2. (F) Relation between the reactivity towards human and *S. cerevisiae* TOP1 by ATA^+^ SSc patients in cohort 2. (A-F) Optical density was measured at 415 nanometers (OD_415nm_) and blank was subtracted. Saturated samples (OD>2.5) were diluted further to determine antibody levels in arbitrary units per milliliter (aU/ml), see Fig. S3. Dotted lines represent cut-off: mean + 2x standard deviation of HDs. (A-B, D-E) Kruskal-Wallis test combined with Dunn’s multiple comparison test was used to test for statistical significant differences. ns = not significant, * = p<0.05, ** = p<0.005, *** = p<0.001. (C, F) Correlations were described using the Spearman’s rank correlation coefficient (r_s_).

To independently replicate these findings, we measured *S. cerevisiae* TOP1 reactivity in serum of ATA^+^ SSc patients in a second cohort (cohort 2, table 2). As in cohort 1, human TOP1 was recognized by IgG of ATA^+^ SSc patients, whereas age- and sex-matched ACA^+^ SSc patients and HDs did not recognize this antigen (Fig. 2D, Fig. S3D). *S. cerevisiae* TOP1 was also recognized by a subset of ATA^+^ SSc patients in this second cohort (62%), but not by ACA^+^ SSc patients and HDs (Fig. 2E, Fig. S3E). A moderate correlation (r_s_=0.52, p<0.001) was found between the recognition of human and *S. cerevisiae* TOP1 (Fig. 2F, Fig. S3F). Similar to the observations in cohort 1, strong heterogeneity was observed in the recognition of *S. cerevisiae* TOP1 between patients with comparable reactivity towards human TOP1. Together, these data indicate that a subgroup of ATA^+^ SSc patients, but not healthy or disease control individuals harbor reactivity towards TOP1 of microbial origin, supporting the concept of potential cross-reactivity.

### Cross-reactivity of patient-derived ATA monoclonal antibodies towards S. cerevisiae TOP1

To more directly investigate cross-reactivity of human ATA to *S. cerevisiae* TOP1, we next used seven patient-derived ATA monoclonal antibodies (ATA mAbs) which were generated based on the BCR sequence of TOP1-reactive B cells from one ATA^+^ SSc patient (Table S4) (*31*). All mAbs showed the expected composition and molecular weight upon purification (Fig. S4A). In line, all ATA mAbs, but not a tetanus toxoid (TT)-reactive control mAb, reacted to human TOP1 (Fig. 3A). Also, none of the mAbs bound the control protein centromere protein B, except for 8D7 which generated a weak signal at high concentrations (Fig. S4B). Reactivity to TOP1 from *S. cerevisiae* was observed for four of the seven ATA mAbs (2F8, 6E3, 8D7, 9D11) in a concentration-dependent manner (Fig. 3B). Binding of ATA-IgG 9D11 to *S. cerevisiae* TOP1 was strong and detectable at lower concentrations (<0.01µg/ml), whereas ATA-IgG 2F8, 6E3 and 8D7 showed detectable binding only at higher concentrations (>1µg/ml). In contrast, the TT-reactive control mAb D2 and three other ATA mAbs (7G6, 2C11 and 7D11) did not recognize *S. cerevisiae* TOP1, even at higher concentrations. Together, these data provide direct evidence that a subset of ATA mAbs reacts to both human and *S. cerevisiae* TOP1.

**Figure 3.**
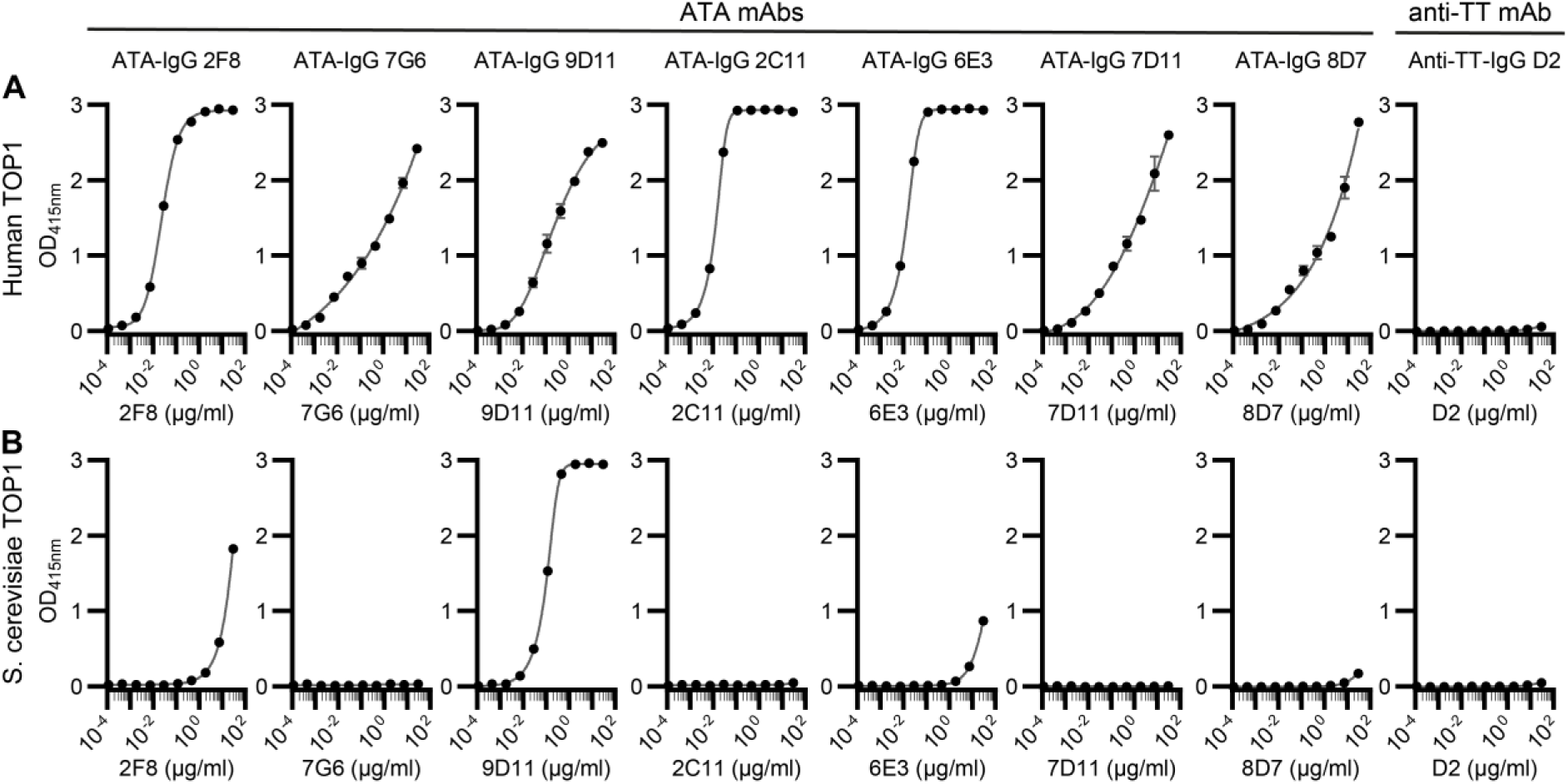
Reactivity of patient-derived ATA mAbs towards human and *S. cerevisiae* TOP1. (A-B) Reactivity of patient-derived ATA mAbs towards human (A) and *S. cerevisiae* TOP1 (B). The mAbs were applied in a 4-fold serial dilution starting at 30µg/ml. A tetanus toxoid (TT)-reactive mAb (anti-TT-IgG D2) was used as negative control. Optical density was measured at 415 nanometers (OD_415nm_) and blank was subtracted. Error bars represent the standard deviation of technical duplicates. Data are representative of three independent experiments. Individual datapoints were connected using a five-parameter logistic regression.

### S. cerevisiae TOP1 can activate an ATA-expressing B cell line

We next investigated the potential functional relevance of the observations described above by assessing the ability of *S. cerevisiae* TOP1 to stimulate B cells directed against human TOP1. To this end, we generated anti-human TOP1 BCR-expressing Ramos B cell lines and studied the ability of *S. cerevisiae* TOP1 to activate these cells in an antigen-specific manner (Fig. 4A). Ramos B cells expressing anti-TOP1-IgG as functional BCR were generated by retroviral transduction of constructs encoding the BCR sequence of B cell clones 2F8, 7G6 and 9D11 (Fig. S5A-B). All TOP1-reactive Ramos B cell lines were activated upon stimulation with anti-IgG (positive control) and human TOP1 (Fig. 4B, Fig. S5C), as evidenced by the phosphorylation of Syk. Importantly, the Ramos B cell line expressing anti-human TOP1 BCR 9D11 was also activated by *S. cerevisiae* TOP1, in contrast to the cell lines expressing 7G6 and 2F8 (Fig. 4A-B, Fig. S5C), corroborating the findings obtained by ELISA. A control Ramos B cell line expressing a citrullinated protein-reactive BCR (ACPA-IgG 3F3) was activated by anti-human IgG but not by human and *S. cerevisiae* TOP1. These data show that TOP1 of microbial origin cannot only be recognized by human ATA, but that it can also activate autoreactive B cells expressing cross-reactive BCRs derived from patients.

**Figure 4.**
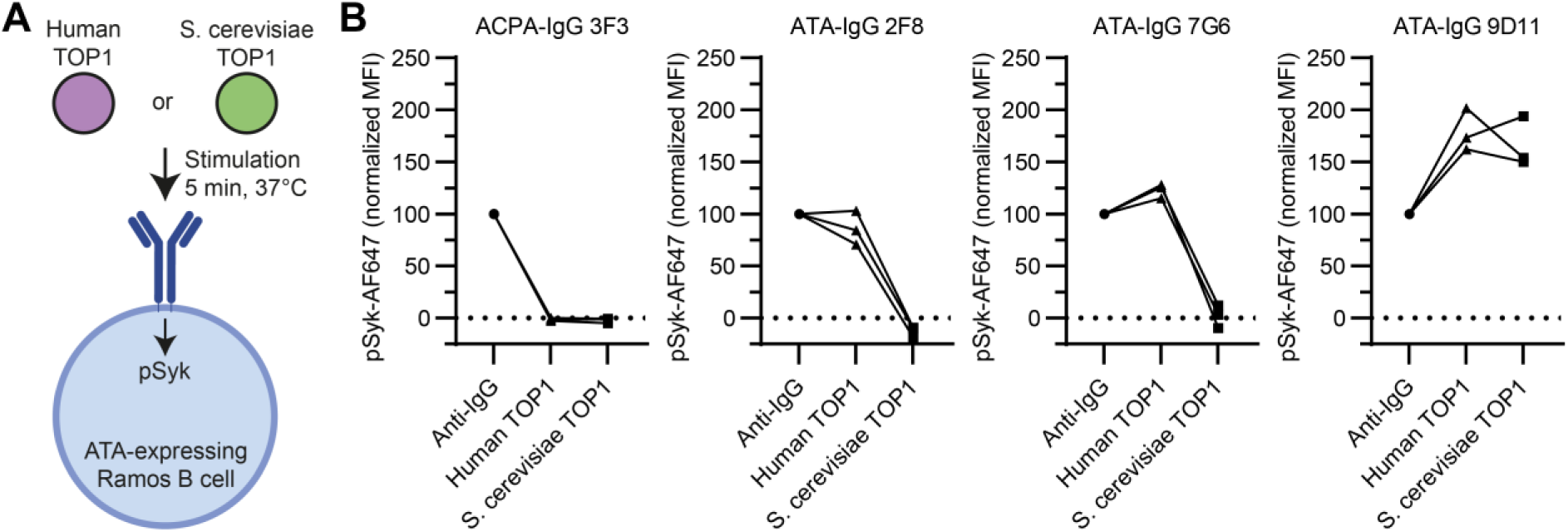
Stimulation of ATA-expressing Ramos B cell lines with human and S. cerevisiae TOP1. (A) Experimental procedure to measure the activation of Ramos B cell lines upon stimulation. ATA-expressing Ramos B cell lines and an ACPA-expressing Ramos B cell line (negative control) were stimulated with 5µg/ml anti-IgG (positive control), human TOP1 or S. cerevisiae TOP1 for 5 minutes at 37°C. Subsequently, the expression of pSyk by stimulated RAMOS B cells was measured by flow cytometry. (B) Expression of pSyk by Ramos B cell lines upon stimulation with anti-IgG, human TOP1 or S. cerevisiae TOP1. MFIs were normalized to the MFI generated in the corresponding anti-IgG condition upon subtracting the MFI obtained in the unstimulated condition. Data is derived from three independent experiments, independent experiment are connected by lines.

### Recognition of S. cerevisiae TOP1 is enhanced in patients with severe disease

We previously demonstrated a possible link between the activity of the TOP1-reactive B cell response and severe disease (*9, 10*). Given the ability of *S. cerevisiae* TOP1 to activate human ATA-expressing B cells, we next questioned whether the recognition of *S. cerevisiae* TOP1 by a subset of ATA^+^ SSc patients would associate with severity of the clinical phenotype. Therefore, we measured anti-human TOP1-IgG and anti-*S. cerevisiae* TOP1-IgG levels in a cohort (cohort 2) consisting of ATA^+^ SSc patients at both ends of the SSc disease severity spectrum: 18 patients with severe SSc and 21 with mild disease (Table 2). In this cohort, levels of anti-human TOP1-IgG were higher in patients with severe disease compared to those with mild SSc (Fig. 5A). Moreover, sera of patients with severe SSc more often harbored antibodies recognizing *S. cerevisiae* TOP1 in comparison to patients with mild SSc (83% versus 43%) (Fig. 5B). Interestingly, the recognition of *S. cerevisiae* TOP1 was particularly observed in patients with clinically relevant ILD (Fig. 5C-D). Male patients and patients with diffuse cutaneous SSc also tended to have higher levels of anti-*S. cerevisiae* TOP1-IgG (Fig. S6A-J). In conclusion, based on the evaluation of patients at the extreme ends of the severity spectrum, we observed that recognition of microbial TOP1 was more prevalent in severe SSc, in particular in those with clinically relevant ILD.

**Figure 5.**
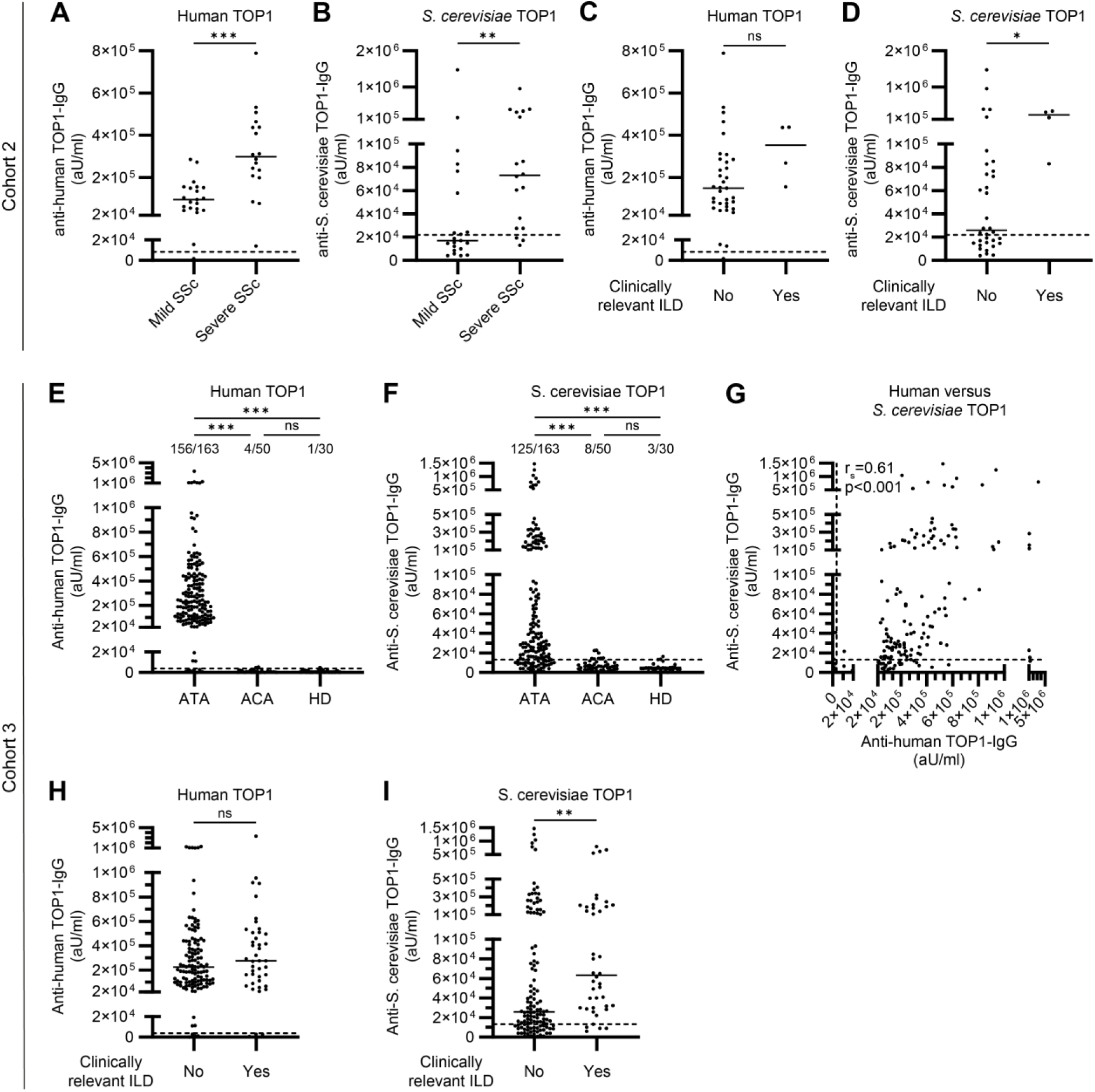
Association between the recognition of *S. cerevisiae* TOP1 and clinical disease in ATA^+^ SSc. (A-B) Levels of anti-human TOP1-IgG (A) and anti-S. cerevisiae TOP1-IgG (B) in serum of ATA^+^ SSc patients included in cohort 2 with mild (n=21) versus severe disease (n=18). (C-D) Comparison of anti-human TOP1-IgG (C) and anti-S. cerevisiae TOP1-IgG (B) levels in serum between ATA^+^ SSc patients included in cohort 2 with and without clinically relevant ILD. (E-F) Reactivity of IgG in plasma of ATA^+^ SSc patients (n=163), ACA+ SSc patients (n=50) and healthy donors (HD)(n=30) included in cohort 3 towards human (E) and S. cerevisiae (F) TOP1. Kruskal-Wallis test combined with Dunn’s multiple comparison test was used to test for statistical significant differences. (G) Correlation between the recognition of human and *S. cerevisiae* TOP1 by ATA^+^ SSc patients in cohort 3. Correlations were described using the Spearman’s rank correlation coefficient (r_s_). (H-I) Comparison of anti-human TOP1-IgG (C) and anti-S. cerevisiae TOP1-IgG (B) levels in serum between ATA^+^ SSc patients included in cohort 3 with and without clinically relevant ILD. (A-I) Levels are expressed in arbitrary units per milliliter (aU/ml). Striped lines represent cut-off: mean + 2x standard devation of HD samples. (A-D, H-I) Mann-Whitney U test was used to test for significant differences. (A-F, H-I) ns = not significant, * = p<0.05, ** = p<0.005, *** = p<0.001.

To replicate the association and evaluate the possible relevance of *S. cerevisiae* TOP1 recognition for clinically relevant ILD in ATA^+^ SSc in a cohort without a selection bias for disease severity, we measured anti-human- and *S. cerevisiae-*TOP1-IgG levels in serum samples from all available ATA^+^ SSc patients (cohort 3, n=163) (*20*). In this cohort, human TOP1 was recognized by the majority of ATA^+^ SSc patients and not by age- and sex-matched ACA^+^ SSc patients and HDs (Fig. 5E, Fig. S7A). A subset of ATA^+^ SSc patients (77%) recognized *S. cerevisiae* TOP1, while most ACA^+^ SSc patients and HDs were negative (Fig. 5F, Fig. S7B). Again, anti-human TOP1- and *S. cerevisiae* TOP1-IgG levels correlated (r_s_=0.61, p<0.001) (Fig. 5G, Fig. S7C). Interestingly, patients with clinically relevant ILD were more commonly anti-*S. cerevisiae* TOP1-IgG^+^ (90% seropositivity in patients with clinically relevant ILD versus 72% in patients without clinically relevant ILD) (Fig. 5H-I). This corresponded to an adjusted odds ratio of 3.4 (95% confidence interval: 1.1 - 10.3, p=0.034), when corrected for sex, disease subset and elevated C-reactive protein levels. Moreover, anti-*S. cerevisiae* TOP1-IgG levels, but not anti-human TOP1-IgG levels, were significantly increased in patients with clinically relevant ILD (Fig. 5H-I). This finding remained statistically significant when patients in cohort 2, who were also included in cohort 3, were excluded (Fig. S7D-E). Also in cohort 3, patients with diffuse cutaneous SSc and male patients tended to have higher levels of *S. cerevisiae* TOP1-IgG (Fig. S7F-O). In conclusion, antibodies recognizing *S. cerevisiae* TOP1 associate with clinically relevant ILD in ATA^+^ SSc.

## Discussion

The clinical management of autoimmune diseases such as SSc is highly complex due to heterogeneity within patient populations and lack of curative treatments. To understand such heterogeneity in the development of disease complications, it is important to understand the mechanisms driving and maintaining autoimmunity. SSc is hallmarked by a break in B cell tolerance towards nuclear proteins. At present, it is unclear how these B cell responses are initiated and activated. Here, we provide evidence that a subset of patient-derived ATAs can cross-react to microbial TOP1 derived from an environmental yeast. More specifically, we found that sera from a subset of ATA^+^ SSc patients and multiple ATA mAbs recognized TOP1 from *S. cerevisiae*, a microbial TOP1 which shares homology with human TOP1. In vitro, *S. cerevisiae* TOP1 induced in the activation of TOP1-reactive B cells. Furthermore, the recognition of *S. cerevisiae* TOP1 was enhanced in patients with severe ATA^+^ SSc and in patients with clinically relevant ILD.

These findings are intriguing as they point to the possibility that microbial antigens initiate and activate TOP1-reactive B cells, a notion supported by a recent large epidemiological study implicating exposure to certain yeast strains to high rates of SSc diagnoses (*32*). Microbes have frequently been implicated in the break of B cell tolerance related to autoimmune disease (*12, 33*). One important mechanism relies on cross-reactivity of autoreactive BCRs to microbial antigens, a feature which has been described for various types of autoantibodies (*11, 34*). An important consideration in this context is that microbial antigen recognition by autoreactive B cells not only activates the BCR (known as signal 1) and, potentially, also innate receptors (signal 2) (*13, 35, 36*). In fact, such B cells may also recruit help from microbial-directed T cells (signal 3) to promote their activation, thereby bypassing the need to break T cell tolerance to human TOP1. In this study, we found that microbial TOP1 can activate B cells expressing an anti-human TOP1 BCR based on cross-reactivity. In this context, it is important to note that we used TOP1 from *S. cerevisiae* as a prototypic yeast-derived TOP1 molecule, but that multiple microbes share TOP1 molecules with structural similarities. This means that not *S. cerevisiae*-derived TOP1 specifically may have a causative role in SSc pathogenesis, but that also TOP1 molecules from other microbial sources could be involved. Nonetheless, irrespective of the origin of microbial TOP1, an intriguing yet unexplored possibility is that the initial B cell response primarily targets the pathogen, and that cross-reactivity to human TOP1 is enhanced or even generated de novo by somatic hypermutation. B cells recruit help from T cells by presenting microbial peptides on MHC class II (*37, 38*). Help from T cells is essential for the maturation of B cell responses by inducing isotype switching, somatic hypermutation and epitope spreading (*39, 40*). Thus, based on our observations, we consider this mechanistic framework relevant for the activation of autoreactive B cells in SSc, a process which likely facilitates the initiation and subsequent maturation of the ATA B cell response in the context of SSc pathogenesis.

Besides this intriguing role that microbial TOP1 might play in the initial break of B cell tolerance towards human TOP1, BCR cross-reactivity could also determine the subsequent dynamics of the ATA B cell response during chronic disease. Exposure to one or multiple microbes, whether in the context of acute infection or chronic exposure (e.g. commensals), could act as continuous source of microbial triggering of TOP1-reactive B cells. As it is possible that ATAs can recognize TOP1 from other microbes as well, variations might exist in avidity and in between patients. Interestingly, TOP1 of fungi showed the strongest structural similarities with human TOP1. Fungi represent a substantial fraction of the human microbiome which, in the context of autoimmunity, has so far received little attention (*41, 42*). Fungi can induce systemic immune reactions and were recently linked to the pathogenesis of various other immune-related diseases, like Crohn’s disease and severe COVID-19 (*28, 41–43*). In the context of SSc, a higher abundance of *Rhodotorula glutinis* RNA sequence reads was detected in the skin of SSc patients, in comparison to HDs (*18*). In addition, reactivity towards the fungal counterparts of human fibrillarin and α-enolase derived from *S. cerevisiae* was detected in sera of SSc patients (*44, 45*). Hence, our findings warrant further exploration of the mycobiome of SSc patients and extended analyses of BCR cross-reactivity to delineate whether fungi indeed serve as a continuous source of antigens triggering autoreactive B cells in this disease.

The association with severe disease makes anti-*S. cerevisiae* TOP1-IgG a promising clinical biomarker. This is particularly important as within patients with reactivity to human TOP1, or ATA positivity, still large heterogeneity exists in clinical disease course for which biomarkers are missing that can stratify between subsets of patients (*21*). The increased recognition of *S. cerevisiae* TOP1, but not human TOP1, by patients with clinically relevant ILD indicates that measuring *S. cerevisiae* TOP1 reactivity might be complementary to measuring human TOP1 reactivity. This additive value is based on the remarkable variability between patients in the recognition of *S. cerevisiae* TOP1, even in patients with comparable anti-human TOP1-IgG levels. Such a biomarker might, ideally in combination with other standard diagnostics, aid in the stratification of ATA+ patients to tailor monitoring, management and treatment of the disease. Additional replication of these findings and longitudinal analyses are warranted to explore the potential of anti-*S. cerevisiae* TOP1-IgG as clinical biomarker. In future steps, it will also be interesting to determine the recognition of microbial TOP1 by individuals with very early SSc, to assess germline-reverted ATA mAbs, and the recognition of other, not yet studied microbial proteins.

In conclusion, the recognition of microbial TOP1 by human ATA and the activation of TOP1-reactive B cells by microbial TOP1 support a role for microbes in the activation of the ATA B cell response in SSc. The increased recognition in patients with severe SSc and in patients with clinically relevant ILD indicates that microbial TOP1 could be a clinically relevant trigger of ATA B cells which relates to disease heterogeneity. Together, these results enhance our understanding of mechanisms underlying the breach of tolerance towards human autoantigens. Also, they hold promise to stratify clinically heterogeneous patient populations based on recognition of a microbial antigen to which the human autoreactive B cell response cross-reacts, and highlight the human mycobiome as an interesting area for further research.

## Acknowledgements

R.Q. Kim and A. Moutsiopoulou (Protein Facility of the Leiden University Medical Center) are thanked for the production of human TOP1. We thank J.F.X. Diffley (The Francis Crick Institute) for providing the yAE42 strain used for production and purification of S. cerevisiae TOP1. Flow cytometry was performed at the Flow Cytometry Core Facility of the Leiden University Medical Center.

## Author contributions

Conceptualization: SN, SIEL, JKdVB, REMT, HUS

Methodology: SN, SIEL, TvL NEWL, NHD, JKdVB, REMT, HUS

Investigation: SN

Analysis of the data – SN, SIEL, EMH

Interpretation of the data – SN, SIEL, TvL, EMH, CMF, NHD, JKdVB, REMT, HUS

Supervision – JKdVB, REMT, HUS

Writing – original draft: SN, HUS

Writing – review & editing: SN, SIEL, TvL, EMH, CMW, NEWL, CMF, NHD, JKdVB, REMT, HUS

