## Supplementary Figures for "Clonal autoantibodies identify microbial antigen as trigger of autoreactive B cells in systemic sclerosis"

### Supplementary Figure 1.

```
Human TOP1            1     WKWWEEERYPEGIKWKFLEHKGPVFAPPYEPLPENVKFYYDGKVMKLSPKAEEVATFFAK    62
                            +KWWE+E   + IKW  L+H G +F PPY+PLP ++K+YYDGK + L P+AEEVA FFA
S. cerevisiae TOP1    131   YKWWEKENEDDTIKWVTLKHNGVIFPPPYQPLPSHIKLYYDGKPVDLPPQAEEVAGFFAA    190

Human TOP1            61    MLDHEYTTKEIFRKNFFKDWRKEMTNEEKNI------ITNLSKCDFTQMSQYFKAQTEAR    114
                            +L+ ++    +F+KNFF D+ +    +E         I  +S+CDFT+M  YF+ Q E +
S. cerevisiae TOP1    191   LLESDHAKNPVFQKNFFNDFLQV—LKESGGPLNGIEIKEFSRCDFTKMFDYFQLQKEQK     248

Human TOP1            115   KQMSKEEKLKIKEENEKLLKEYGFCIMDNHKERIANFKIEPPGLFRGRGNHPKMGMLKRR    174
                            KQ++ +EK +I+ E EK+ ++Y FC +D  +E+++NFK+EPP LFRGRG HPK G LKRR
S. cerevisiae TOP1    249   KQLTSQEKKQIRLEREKFEEDYKFCELDGRREQVGNFKVEPPDLFRGRGAHPKTGKLKRR    308

Human TOP1            175   IMPEDIIINCSKDAKVPSPPPGHKWKEVRHDNKVTWLVSWTENIQGSIKYIMLNPSSRIK    234
                            + PEDI++N SKDA VP  P GHKW E+RHDN V WL  W ENI  S KY+ L  +S +K
S. cerevisiae TOP1    309   VNPEDIVLNLSKDAPVPPAPEGHKWGEIRHDNTVQWLAMWRENIFNSFKYVRLAANSSLK    368

Human TOP1            235   GEKDWQKYETARRLKKCVDKIRNQYREDWKSKEMKVRQRAVALYFIDKLALRAGNEKEEG    294
                            G+ D++K+E AR+LK  +D IR  Y  + KSK M  RQ+AVA+Y+ID +ALRAG EK E
S. cerevisiae TOP1    369   GQSDYKKFEKARQLKSYIDAIRRDYTRNLKSKVMLERQKAVAIYLIDVFALRAGGEKSED    427

Human TOP1            295   ETADTVGCCSLRVEHINLHPELDGQEYVVEFDFLGKDSIRYYNKVPVEKRVFKNLQLF-M    353
                              ADTVGCCSLR EH+ L P        V FDFLGKDSIR+Y++V V+K+VFKNL +F
S. cerevisiae TOP1    428   -EADTVGCCSLRYEHVTLKPPN-----TVIFDFLGKDSIRFYQEVEVDKQVFKNLTIFKR    482

Human TOP1            354   ENKQPEDDLFDRLNTGILNKHLQDLMEGLTAKVFRTYNASITLQQQLKELTAPDENIPAK    413
                              KQP   LFDRL+  ILNK+LQ+ M GLTAKVFRTYNAS T+Q QL +L     ++  K
S. cerevisiae TOP1    483   PPKQPGHQLFDRLDPSILNKYLQNYMPGLTAKVFRTYNASKTMQDQL-DLIPNKGSVAEK    541

Human TOP1            414   ILSYNRANRAVAILCNH----------------------------------QIA------    432
                            IL YN ANR VAILCNH                                  Q+
S. cerevisiae TOP1    542   ILKYNAANRTVAILCNHQRTVTKGHAQTVEKANNRIQELEWQKIRCKRAILQLDKDLLKK    601

Human TOP1            433   ------------------------------------------------------------    433

S. cerevisiae TOP1    602   EPKYFEEIDDLTKEDEATIHKRIIDREIEKYQRKFVRENDKRKFEKEELLPESQLKEWLE    661

Human TOP1            433   ----------------------------------------------------------LG    435
                                                                                      LG
S. cerevisiae TOP1    662   KVDEKKQEFEKELKTGEVELKSSWNSVEKIKAQVEKLEQRIQTSSIQLKDKEENSQVSLG    721

Human TOP1            436   TSKLNFLDPRITVAWCKKWGVPIEKIYNKTQREKFAWAIDMADEDYEF                483
                            TSK+N++DPR++V +CKK+ VPIEKI+ KT REKF WAI+  DE++ F
S. cerevisiae TOP1    722   TSKINYIDPRLSVVFCKKYDVPIEKIFTKTLREKFKWAIESVDENWRF                769
```

**Supplementary Figure 1. Sequence homology between human and *S. cerevisiae* TOP1.**
Alignment of the amino acid sequences of human and *S. cerevisiae* TOP1 as determined by a Foldseek search using a crystal structure of human TOP1 (1EJ9, Protein Data Bank). The sequence of *S. cerevisiae* is derived from the AlphaFold/Proteome (v4) database.

**Supplementary Figure 2.**

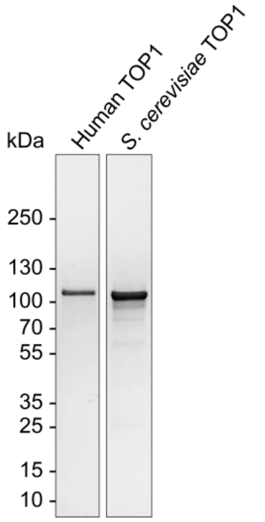


**Supplementary Figure 2. Gel electrophoresis of human and *S. cerevisiae* TOP1.**
Gel electrophoresis of human and *S. cerevisiae* TOP1 upon purification under non-reducing conditions on a polyacrylamide gel with a 4-15% gradient. Protein was visualized with a Coomassie protein stain and molecular weight in kilodaltons (kDa) was estimated using a protein ladder.

### Supplementary Figure 3.

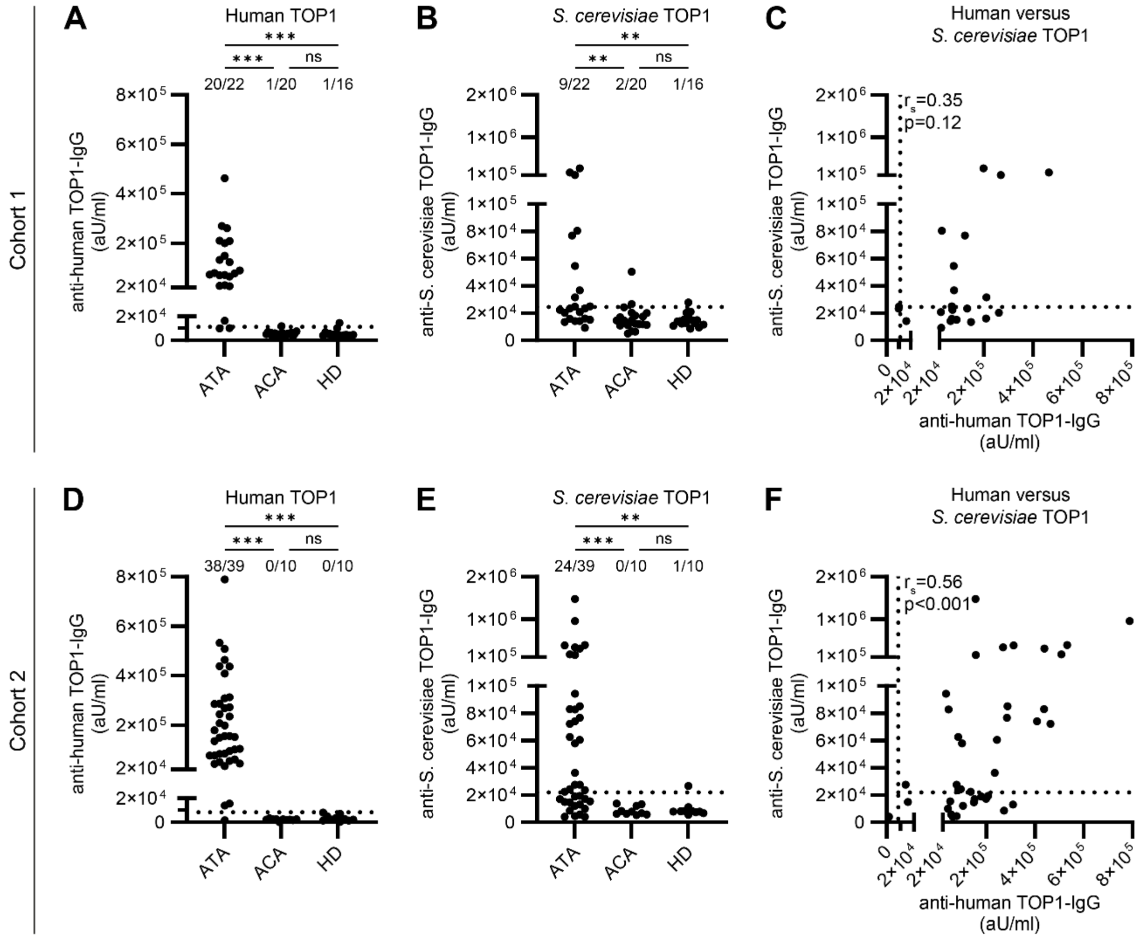


**Supplementary Figure 3. Recognition of *S. cerevisiae* TOP1 by circulating IgG of ATA⁺ SSc patients**
**(A-B)** Levels of anti-human TOP1-IgG (A) and anti-*S. cerevisiae* TOP1-IgG (B) in plasma of ATA⁺ SSc patients (n=22), ACA⁺ SSc patients (n=20) and healthy donors (HD)(n=16) included in cohort 1. **(C)** Correlation between the anti-human TOP1-IgG and anti-*S. cerevisiae* TOP1-IgG levels in the ATA⁺ SSc patients of cohort 1. **(D-E)** Anti-human TOP1-IgG (A) and anti-*S. cerevisiae* TOP1-IgG (B) levels in serum of ATA⁺ SSc patients (n=39), ACA⁺ SSc patients (n=10) and healthy donors (HD)(n=10) included in cohort 2. **(F)** Association between the levels of anti-human TOP1-IgG and anti-*S. cerevisiae* TOP1-IgG in ATA⁺ SSc patients of cohort 2. (A-F) Levels are expressed in arbitrary units per milliliter (aU/ml). (A-B, D-E) Kruskal-Wallis test combined with Dunn's multiple comparison test was used to test for statistical significant differences. ns = not significant, * = p<0.05, ** = p<0.005, *** = p<0.001. (C, F) Correlations were described using the Spearman's rank correlation coefficient ($r_s$).

**Supplementary Figure 4.**

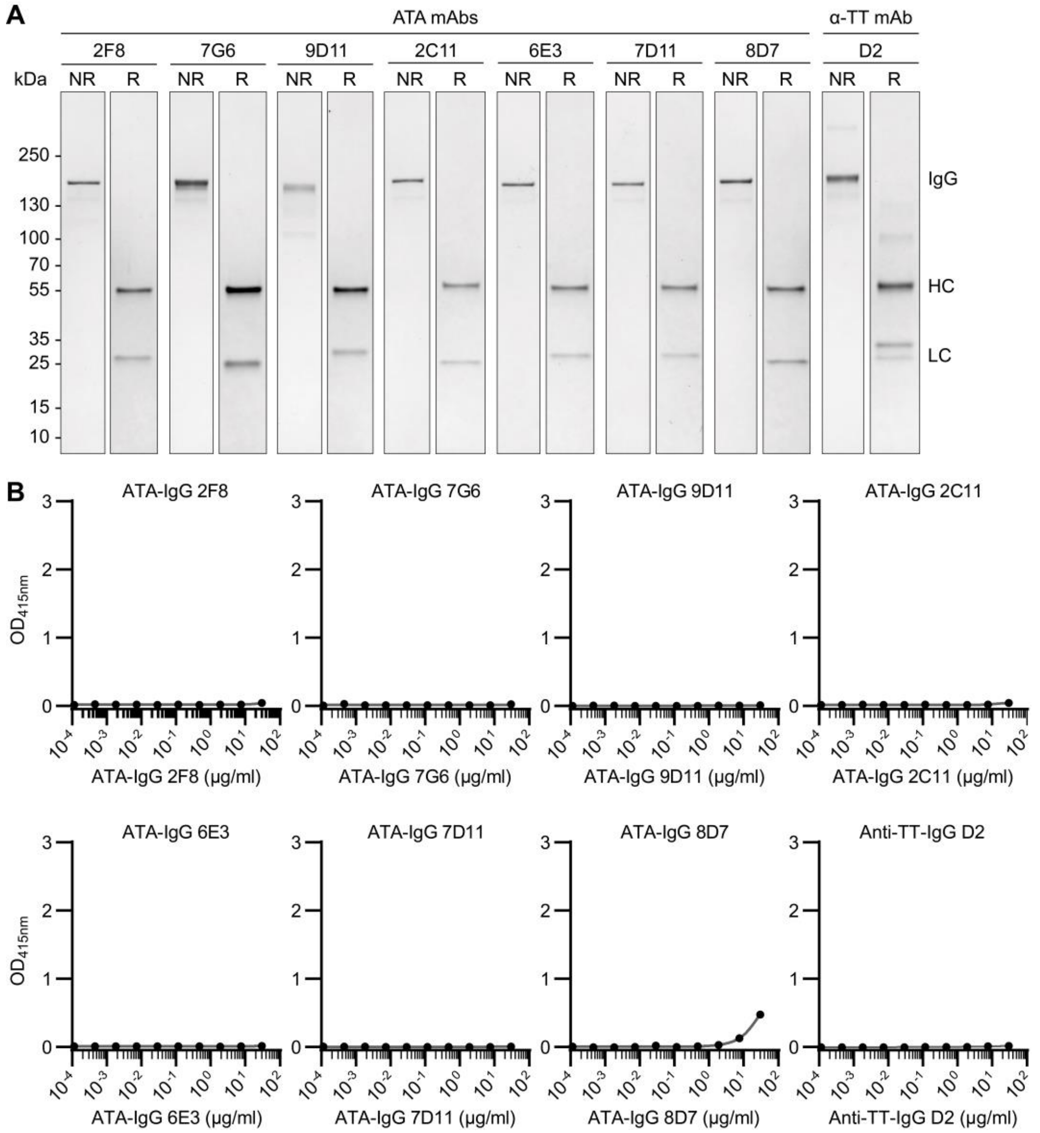


**Supplementary Figure 4. Generation of patient-derived ATA mAbs.**
**(A)** Gel electrophoresis of patient-derived ATA mAbs and an anti-TT IgG mAb upon Protein G purification under non-reducing (NR) and reducing (R) conditions on a polyacrylamide gel with a 4-15% gradient. Protein was visualized with a Coomassie protein stain and molecular weight in kilodaltons (kDa) was estimated using a protein ladder. **(B)** Reactivity of recombinantly produced ATA-IgG mAbs and anti-TT-IgG D2 towards centromeric protein B (a control protein) in an anti-centromeric protein B antibody-IgG (ACA-IgG) ELISA. mAbs were applied in a 4-fold serial dilution starting at 30µg/ml. Optical density was measured at 415 nm ($OD_{415nm}$) and blank was subtracted. Error bars represent standard deviation of technical duplicates. Data is representative of two independent experiments.

**Supplementary Figure 5. Generation of ATA-expressing RAMOS B cell lines.**

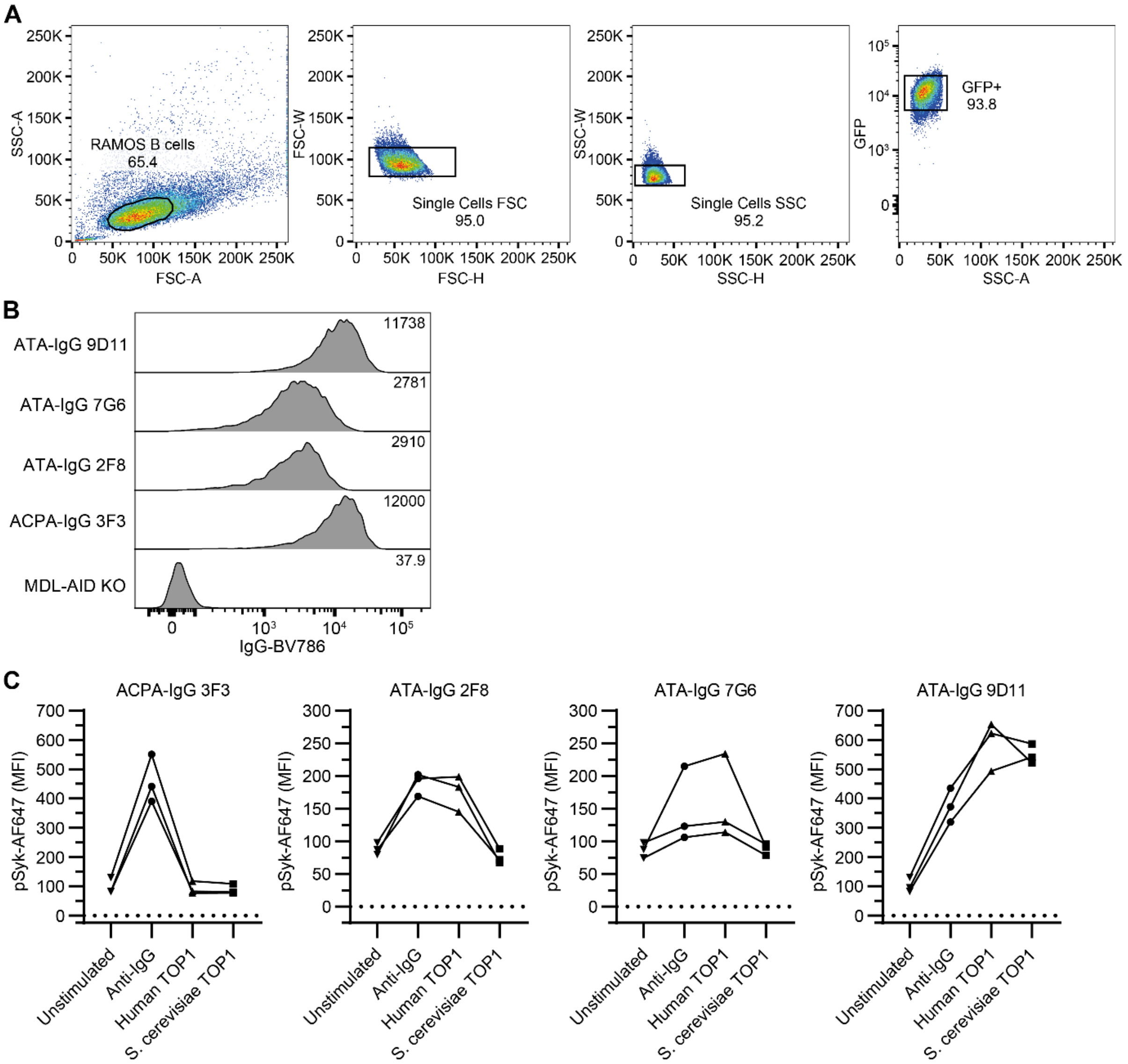


**Supplementary Figure 5. Generation of ATA-expressing Ramos B cell lines.**
**(A)** Gating strategy used to identify Ramos B cells by flow cytometry based on forward and side scatter area (FSC-A and SSC-A, respectively), on the exclusion of doublets based on FSC-width (FSC-W) versus FSC-height (FSC-H) and SSC-width (SSC-W) versus SSC-height (FSC-H), and based on the expression of green fluorescent protein (GFP). **(B)** Expression of IgG on the surface of the ATA-expressing Ramos B cell lines (ATA-IgG 2F8, 7G6 and 9D11) and an ACPA-IgG 3F3-expressing Ramos B cell line. Numbers indicate the median fluorescence intensity (MFI). MDL-AID KO cells were used as negative control. **(C)** Expression of pSyk by ATA-expressing Ramos B cell lines upon stimulation for 5 minutes with 5µg/ml anti-IgG, human TOP1 and *S. cerevisiae* TOP1. ACPA-IgG 3F3-expressing Ramos B cell line was used as negative control for the stimulation with human and *S. cerevisiae* TOP1. Expression of pSyk is expressed as MFI. Data is derived from three independent experiments, independent experiment are connected by lines.

### Supplementary Figure 6.

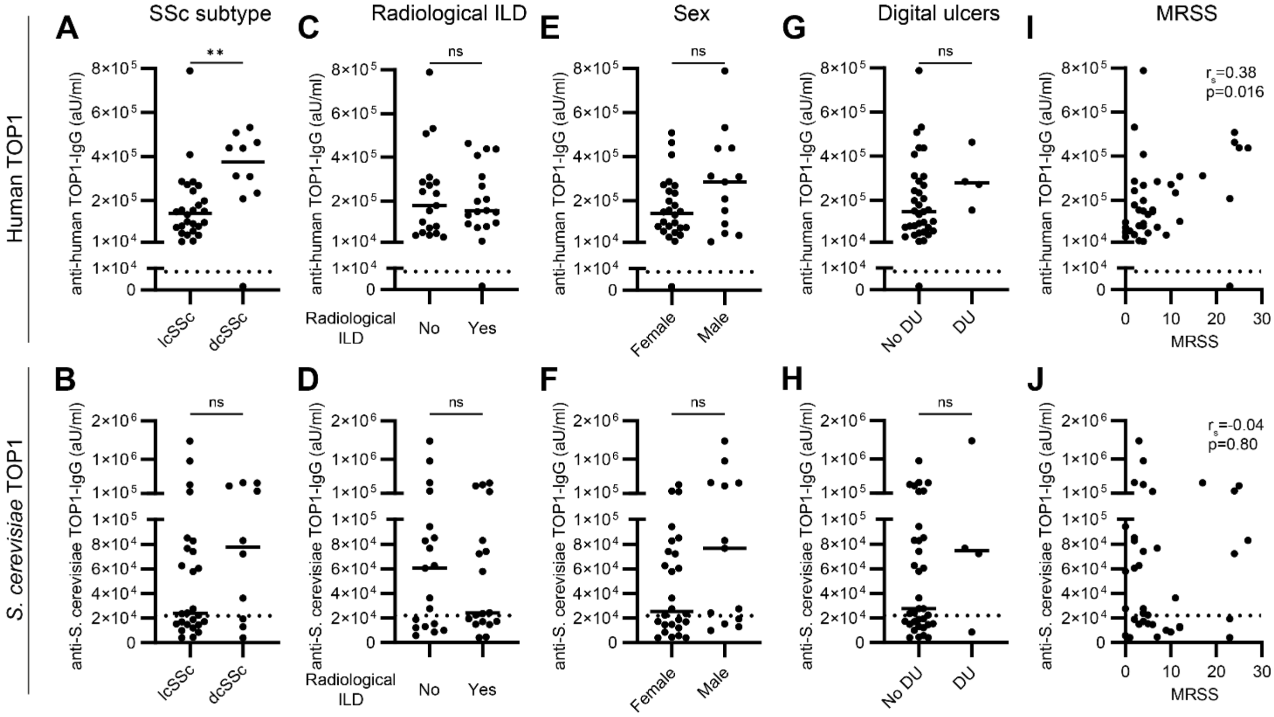


**Supplementary Figure 6. Association between the recognition of *S. cerevisiae* TOP1 and clinical disease parameters in ATA⁺ SSc patients included in cohort 2.**

**(A-H)** Comparison of the levels of anti-human TOP1-IgG and anti-*S. cerevisiae* TOP1-IgG between ATA⁺ SSc patients included in cohort 2 (n=39) with limited cutaneous SSc (lcSSc) versus diffuse cutaneous SSc (dcSSc) (A-B), with versus without interstitial lung disease (ILD) on high resolution computed tomography (HRCT) (C-D), between male and female subjects (E-F), and patients with or without digital ulcers (G-H). Mann-Whitney U test was used to test for significant differences. ns = not significant, * = p<0.05, ** = p<0.005, *** = p<0.001. **(I-J)** Association between the recognition of human and *S. cerevisiae* TOP1 by circulating IgG and MRSS of ATA⁺ SSc patients included in cohort 2 (n=39). Correlations were described using the Spearman's rank correlation coefficient ($r_s$). (A-J) Levels are expressed in arbitrary units per milliliter (aU/ml).

**Supplementary Figure 7.**

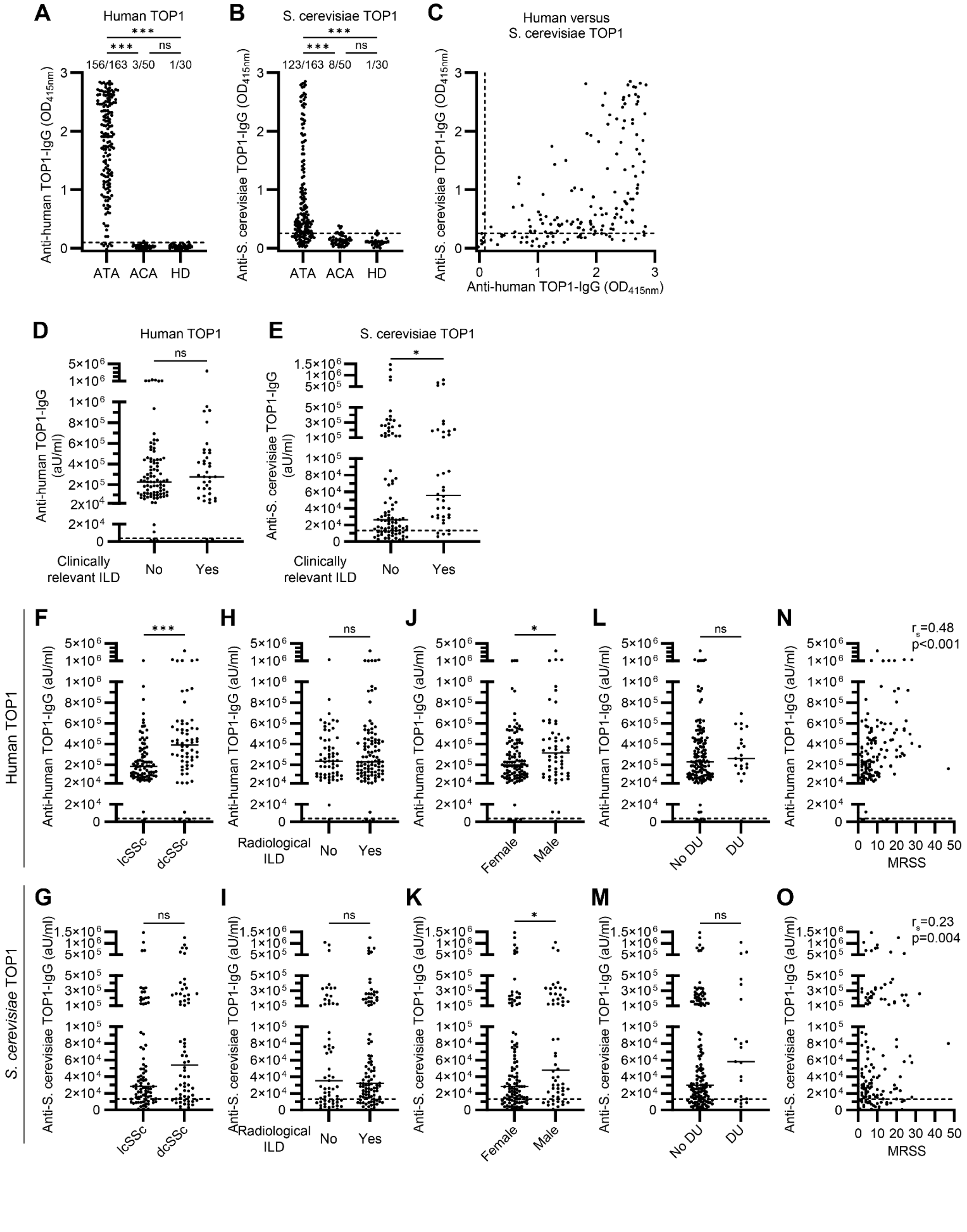

**Supplementary Figure 7. Association between the recognition of *S. cerevisiae* TOP1 and clinical disease parameters in ATA$^+$ SSc patients included in cohort 3.**

**(A-B)** Binding of human (A) and *S. cerevisiae* (B) TOP1 by IgG in serum of ATA$^+$ SSc patients (n=163), ACA$^+$ SSc patients (n=50) and healthy donors (HD)(n=30) included in cohort 3. Optical density was measured at 415 nanometers ($OD_{415nm}$) and blank was subtracted. Saturated samples (OD>2.5) were diluted further to determine antibody levels in arbitrary units per milliliter (aU/ml), see Figure 5. Kruskal-Wallis test combined with Dunn's multiple comparison test was used to test for statistical significant differences. **(C)** Relation between the reactivity towards human and *S. cerevisiae* TOP1 by ATA$^+$ SSc patients in cohort 3. **(D-E)** Comparison of anti-human TOP1-IgG (D) and anti-S. cerevisiae TOP1-IgG (E) levels in serum between ATA$^+$ SSc patients included in cohort 3, with exclusion of patients which are also included in cohort 2, with and without clinically relevant ILD. **(F-M)** Comparison of the levels of anti-human TOP1-IgG and anti-*S. cerevisiae* TOP1-IgG between ATA$^+$ SSc patients included in cohort 3 with limited cutaneous SSc (lcSSc) versus diffuse cutaneous SSc (dcSSc) (F-G), with versus without interstitial lung disease (ILD) on high resolution computed tomography (HRCT) (H-I), between male and female subjects (J-K), and patients with or without digital ulcers (L-M). **(N-O)** Correlation between the recognition of human and *S. cerevisiae* TOP1 and MRSS by circulating IgG of ATA$^+$ SSc patients included in cohort 3. (A-O) Striped lines represent cut-off: mean + 2x standard devation of HDs. (D-O) Levels are expressed in arbitrary units per milliliter (aU/ml). (C, N-O) Correlations were described using the Spearman's rank correlation coefficient ($r_s$). (D-M) Mann-Whitney U test was used to test for significant differences. (A-B, D-M) ns = not significant, * = p<0.05, ** = p<0.005, *** = p<0.001.
