## Supplementary Materials for "Clonal autoantibodies identify microbial antigen as trigger of autoreactive B cells in systemic sclerosis"

*Analysis of structural homology between human and microbial TOP1 using FoldSeek*

Structural homology between human TOP1 and microbial proteins was analyzed using Foldseek (*1*). A crystal structure of human TOP1 (1EJ9, Protein Data Bank) was used as input (*2*). Foldseek was operated in default mode (3Di/AA) with an E-value cut-off of 0.001 to search databases AlphaFold/Uniprot50 (v4), AlphaFold/Swiss-Prot (v4), AlphaFold/Proteome (v4), CATH50 (v3.4.0) and PDB100 (version 20240101). Taxonomic filters were applied to specifically identify proteins from fungi, bacteria and viruses.

*Recombinant production and purification of human TOP1*

Human TOP1 was recombinantly expressed in insect cells (Protein Facility, Leiden University Medical Center) and subsequently purified as described in detail before (*3*). In short, a recombinant bacmid encoding full-length human TOP1 with a N-terminal polyhistidine tag (6x histidine) was generated and transfected into sedentary Sf9 insect cells (ThermoFisher). Culture supernatant containing baculovirus was used to transduce suspension Sf9 cells to recombinantly express human TOP1. Upon culturing, TOP1 was purified from cell lysates using a HisTrap HP column (Cytiva) followed by cation exchange chromatography (HiTrap SP FF, Cytiva). Purity and integrity of purified human TOP1 was validated by gel electrophoresis.

*Recombinant production and purification of S. cerevisiae TOP1*

*S. cerevisiae* TOP1 expression was induced using galactose in a *S. cerevisiae* cell line with a galactose-inducible *S. cerevisiae* TOP1 gene containing a N-terminal calmodulin purification tag derived from the Calmodulin Binding Protein (CPB) followed by a TEV protease recognition sequence (yAE42 cells, provided by Dr. J.F.X. Diffley (*4*)). *S. cerevisiae* TOP1 expression was induced by addition of 2% galactose to yAE42 cells cultured in YP medium (1% yeast extract, 2% peptone) supplemented with 2% raffinose. Upon culturing for three hours, cells were harvested and resuspended in TOP1 lysis buffer (25mM Tris-

HCl (pH=7.5), 0.02% NP-40 substitute, 10% glycerol, 300mM NaCl, and 1mM DTT) supplemented with protease inhibitors (cOmplete$^{TM}$, EDTA-free Protease Inhibitor Cocktail, Sigma) and 0.3mM phenylmethylsulphonyl fluoride. Cell suspension was dropped into liquid nitrogen and frozen droplets were ground in a freezer mill (6875, SPEX SamplePrep) for six cycles (2 minute run time, 1 minute cool time, rate of 15 cycles per second). Resulting powder was resuspended in TOP1 lysis buffer supplemented with protease inhibitors and lysate was cleared by centrifugation. Cleared lysate was supplemented with $CaCl_2$ to a final concentration of 2mM and incubated with Sepharose 4B Calmodulin Beads (GE Healthcare) for 1 hour at 4°C. Beads were washed 20 times with TOP1 binding buffer (25mM Tris-HCl (pH=7.5), 0.02% NP-40 substitute, 10% glycerol, 300mM NaCl, 2 mM $CaCl_2$ and 1mM DTT) and protein was eluted using TOP1 elution buffer (25mM Tris-HCl (pH=7.5), 0.02% NP-40 substitute, 10% glycerol, 300mM NaCl, 2 mM EDTA, 2mM EGTA, and 1 mM DTT). *S. cerevisiae* TOP1 was further purified by size exclusion chromatography using a Superose 6 increase 10/300 GL column (GE Healthcare) equilibrated in TOP1 GF buffer (25mM Tris-HCl pH 7.5, 0.02% NP-40 substitute, 10% glycerol, 150mM NaCl and 1mM DTT). Purity and integrity of purified *S. cerevisiae* TOP1 was validated by gel electrophoresis.

*Generation of patient-derived ATA mAbs*

Patient-derived ATA mAbs were generated as described in detail previously (*3*). In short, BCR sequences obtained from single-cell sorted human TOP1-reactive B cells of an ATA$^+$ SSc patient were cloned into plasmids encoding antibody constant domains to recombinantly produce IgG1 antibodies in FreestyleTM 293-F cells (Gibco) (Table S4). Produced IgG was purified from culture supernatants using a Protein G column (GE Healthcare) and purity and integrity of the generated mAbs was validated by gel electrophoresis. An anti-TT-IgG mAb, generated in a similar procedure as described above, was used as negative control.

*Gel electrophoresis*

The purity and integrity of human and *S. cerevisiae* TOP1 and the mAbs were assessed by sodium dodecyl sulfate polyacrylamide gel electrophoresis (SDS-PAGE). Human and *S. cerevisiae* TOP1 was loaded on a polyacrylamide gel with a 4-15% gradient upon addition of Laemmli buffer (Bio-Rad). The mAbs were loaded on a polyacrylamide gel with a 4-15% gradient upon incubation for five minutes at 95°C in Laemmli buffer in presence (reduced) or absence (non-reduced) of 2% β-mercaptoethanol (Merck). PageRuler™ Plus Prestained Protein ladder (ThermoFisher Scientific) was used as reference. Proteins were separated for 60-90 minutes at 100-120 volt. Upon running the gels, gels were washed in Milli-Q water for 5 minutes and protein was subsequently stained using InstantBlue® Coomassie Protein Stain (Abcam) for one hour. Upon overnight destaining in Milli-Q water, gels were visualized using a ChemiDoc™ Touch Imaging system (Bio-Rad).

*Anti-human TOP1-IgG and anti-S. cerevisiae TOP1-IgG ELISA*

High binding 384-well microplates (Corning) were coated with 2.5μg/ml human or *S. cerevisiae* TOP1 in PBS (pH=8). Plates were blocked with 1% BSA in PBS and samples were subsequently applied. Serum samples were diluted 3200x or 800x for the ELISA coated with human and *S. cerevisiae* TOP1, respectively. Saturated serum samples were diluted further. The mAbs were applied in a four-fold serial dilution with a starting concentration of 30μg/ml. Pooled serum samples of ATA$^+$ SSc patients were used as standards. IgG was detected using 1:5000 diluted goat anti-human IgG-HRP (DAKO, P0214) with ABTS as substrate. Plates were measured with a Spectramax I3x Multi-Mode Microplate Reader (Molecular Devices) using SoftMax® Pro 7 (version 7.0.2, Molecular Devices). Samples of HDs were used to determine the cut-off (mean plus two times the standard deviation) for anti-human TOP1-IgG and anti-*S. cerevisiae* TOP1-IgG seropositivity.

*Generation of Ramos B cell lines expressing TOP1-reactive IgG BCRs*

Ramos B cell lines expressing TOP1-reactive IgG BCRs were generated based on the BCR sequence of B cell clones 2F8, 7G6 and 9D11 (Table S4). To this end, vectors encoding the variable domain of these

BCRs as membrane-bound IgG were retrovirally transduced into a Ramos B cell line in which IgD, IgM and AID was knocked out (MDL-AID KO Ramos cells), as described in detail before (*5, 6*). To determine GFP and surface IgG expression on the transduced cells, Ramos cells were stained with anti-human IgG-Fc BV786 (564230, BD Biosciences) for 30 minutes on ice and subsequently measured using a BD LSRFortessa 4L (BD Biosciences). Data was analyzed using FlowJo (version 10.9.0). Ramos cells were sorted repeatedly using a CytoFLEX SRT sorter (Beckman Coulter) based on GFP and surface IgG expression.

*Stimulation of ATA-expressing Ramos B cell lines with human and S. cerevisiae TOP1*

ATA-expressing Ramos B cell lines were stimulated with 5µg/ml human TOP1, *S. cerevisiae* TOP1 or polyclonal AffiniPure™ F(ab')$_2$ fragment goat anti-human IgG (109-006-097, Jackson ImmunoResearch) for 5 minutes at 37°C in RPMI 1640 supplemented with 100 units/ml penicillin/streptomycin, glutamax, 10mM HEPES (all Gibco) and 1% FCS (Bodinco). Upon stimulation, cells were fixed in 4% paraformaldehyde (BioLegend) for 20 minutes at 37°C and subsequently permeabilized in True-Phos™ Perm Buffer (BioLegend) for 75 minutes at -20°C. Afterwards, cells were stained with anti-Syk(pY352)-Alexa Fluor® 647 (557817, BD Biosciences) diluted tenfold in PBS, 0.5% BSA, 0.02% $NaN_3$. Cells were measured on a BD LSRFortessa 4L flow cytometer (BD Biosciences). Data were analyzed using FlowJo (version 10.9.0).

*Study approval*

The study was approved by the ethical committee of the Leiden University Medical Center (protocol number P17.151) and the Leiden University Biobank Toetsing Commissie (CME no. B16.037, REU 043/SH/sh, R09.003/SH/s, LuVDS23.039 and LuVDS25.011). All patients and HDs gave written informed consent for study participation.

*Statistical analysis*

The statistical analysis plan with defined primary and secondary research questions (indicated for separate analyses below) was formulated before performance of the statistical analysis. The researcher performing the laboratory studies (SN) was blinded for clinical parameters of SSc patients, and statistical analyses were performed by other researchers (SIEL/EMH). Use of statistical tests are indicated in the figure legends. A p-value < 0.05 was considered statistically significant. Statistical analyses were performed using Graphpad Prism (version 9.3.1) and IBM SPSS Statistics (version 29.0.0.0).

Cohort 1-3 - Kruskal Wallis test combined with Dunn's multiple comparisons test was used to statistically compare anti-human TOP1-IgG and anti-*S. cerevisiae* TOP1-IgG levels between ATA$^+$ SSc patients, ACA$^+$ SSc patients and HDs. The non-parametric Spearman's rank correlation coefficient was used to describe correlations between anti-human TOP1-IgG and anti-*S. cerevisiae* TOP1-IgG levels.

Cohort 2 - Primary research question: Does the recognition of *S. cerevisiae* TOP1 differ between patients with mild versus severe SSc? Mann-Whitney U test was used to test for statistically significant differences in anti-human TOP1-IgG and anti-*S. cerevisiae* TOP1-IgG levels between patients with mild versus severe SSc. Relation between other patient characteristics (clinically relevant ILD, radiographic ILD, disease subset, sex, MRSS and digital ulcers) and anti-human TOP1-IgG or anti-*S. cerevisiae* TOP1-IgG levels were explored using the Mann-Whitney U test or described using the non-parametric Spearman's rank correlation coefficient for dichotomous or continuous variables, respectively.

Cohort 3 - Primary research question: Does the recognition of *S. cerevisiae* TOP1 associate with clinically relevant ILD? Clinically relevant ILD was defined as primary research outcome based on the relation between anti-*S. cerevisiae* TOP1 and clinically relevant ILD observed in cohort 2, the strong association between ATA and ILD, and clinically relevant ILD being one of the main and most prevalent cause of morbidity and mortality in SSc (*7, 8*). To evaluate the independent association between the presence of anti-*S. cerevisiae* TOP1-IgG and clinically relevant ILD, a multivariable logistic regression was performed, adjusted for sex, disease subset and elevated C-reactive protein (CRP > 5mg/ml). These variables are known risk factors for interstitial lung disease (*7, 9-11*). Mann-Whitney U test was used to test for statistically significant differences in anti-human TOP1-IgG and anti-*S. cerevisiae* TOP1-IgG levels between patients with versus without clinically relevant ILD. Relation between other patient characteristics (radiographic ILD, disease subset, sex, MRSS and digital ulcers) and anti-human TOP1-IgG or anti-*S. cerevisiae* TOP1-IgG levels were analyzed using the Mann-Whitney U test or the non-parametric Spearman's rank correlation coefficient for dichotomous or continuous variables, respectively.
