## Supplementary Tables for "Clonal autoantibodies identify microbial antigen as trigger of autoreactive B cells in systemic sclerosis"

**Supplementary table 1. Fungal proteins having structural similarities with human TOP1 as identified by Foldseek.**

| No. | Target | Description | Scientific name | Prob. | Seq. id. | E-value | Score | Query pos. | Target pos. | Database |
|---|---|---|---|---|---|---|---|---|---|---|
| 1 | AF-A0A0W4ZQX5-F1-model_v4 | DNA topoisomerase I | Pneumocystis carinii B80 | 1,00 | 42,7 | 3,47E-55 | 2270 | 1-483 (483) | 203-838 (838) | AlphaFold/UniProt50 (v4) |
| 2 | AF-A0A397IWA0-F1-model_v4 | DNA topoisomerase I | Diversispora epigaea | 1,00 | 42,5 | 4,54E-55 | 2250 | 1-483 (483) | 253-885 (885) | AlphaFold/UniProt50 (v4) |
| 3 | AF-R4XH22-F1-model_v4 | DNA topoisomerase I | Taphrina deformans PYCC 5710 | 1,00 | 40,7 | 6,84E-54 | 2224 | 1-483 (483) | 297-931 (931) | AlphaFold/UniProt50 (v4) |
| 4 | AF-P04786-F1-model_v4 | DNA topoisomerase 1 | Saccharomyces cerevisiae S288C | 1,00 | 41,2 | 9,29E-54 | 2176 | 1-483 (483) | 131-769 (769) | AlphaFold/Swiss-Prot (v4) |
| 5 | AF-P04786-F1-model_v4 | DNA topoisomerase 1 | Saccharomyces cerevisiae S288C | 1,00 | 41,2 | 9,99E-54 | 2176 | 1-483 (483) | 131-769 (769) | AlphaFold/Proteome (v4) |
| 6 | AF-A0A5C3QDJ1-F1-model_v4 | DNA topoisomerase I | Pterula gracilis | 1,00 | 48,3 | 2,96E-53 | 2149 | 12-483 (483) | 186-664 (664) | AlphaFold/UniProt50 (v4) |
| 7 | AF-A0A1G4IVE0-F1-model_v4 | DNA topoisomerase I | Lachancea dasiensis | 1,00 | 40,2 | 6,32E-53 | 2155 | 1-483 (483) | 136-772 (772) | AlphaFold/UniProt50 (v4) |
| 8 | AF-A0A1X2HAU0-F1-model_v4 | DNA topoisomerase I | Syncephalastrum racemosum | 1,00 | 40,4 | 7,85E-53 | 2156 | 1-483 (483) | 53-683 (683) | AlphaFold/UniProt50 (v4) |
| 9 | AF-I2H4R6-F1-model_v4 | DNA topoisomerase I | Tetrapisispora blattae CBS 6284 | 1,00 | 41,5 | 1,35E-52 | 2149 | 1-483 (483) | 240-879 (879) | AlphaFold/UniProt50 (v4) |
| 10 | AF-A0A075AMX5-F1-model_v4 | DNA topoisomerase I | Rozella allomycis CSF55 | 1,00 | 38,7 | 1,42E-52 | 2159 | 1-483 (483) | 111-737 (737) | AlphaFold/UniProt50 (v4) |
| 11 | AF-A0A376BB63-F1-model_v4 | DNA topoisomerase I | Saccharomycodes ludwigii | 1,00 | 41,3 | 1,50E-52 | 2144 | 1-483 (483) | 62-700 (700) | AlphaFold/UniProt50 (v4) |
| 12 | AF-A0A5B0Q3Y8-F1-model_v4 | DNA topoisomerase I | Puccinia graminis f. sp. tritici | 1,00 | 35,3 | 1,59E-52 | 2178 | 1-483 (483) | 2-697 (697) | AlphaFold/UniProt50 (v4) |
| 13 | AF-G8JQY1-F1-model_v4 | DNA topoisomerase I | Eremothecium cymbalariae DBVPG#7215 | 1,00 | 40,8 | 3,21E-52 | 2140 | 1-483 (483) | 22-660 (660) | AlphaFold/UniProt50 (v4) |
| 14 | AF-W4K4L9-F1-model_v4 | DNA topoisomerase I | Heterobasidion irregulare TC 32-1 | 1,00 | 39,7 | 3,39E-52 | 2129 | 1-483 (483) | 149-788 (788) | AlphaFold/UniProt50 (v4) |
| 15 | AF-A0A165EC69-F1-model_v4 | DNA topoisomerase I | Exidia glandulosa HHB12029 | 1,00 | 41,5 | 3,78E-52 | 2145 | 1-483 (483) | 77-719 (719) | AlphaFold/UniProt50 (v4) |
| 16 | AF-A0A1D8PPI2-F1-model_v4 | DNA topoisomerase I | Candida albicans SC5314 | 1,00 | 39,6 | 4,45E-52 | 2105 | 1-483 (483) | 142-780 (780) | AlphaFold/Proteome (v4) |
| 17 | AF-A0A098VWA3-F1-model_v4 | DNA topoisomerase I | Mitosporidium daphniae | 1,00 | 39,1 | 4,45E-52 | 2083 | 2-483 (483) | 73-689 (689) | AlphaFold/UniProt50 (v4) |
| 18 | AF-A0A2N5SWR3-F1-model_v4 | DNA topoisomerase I | Puccinia coronata f. sp. avenae | 1,00 | 37,8 | 4,96E-52 | 2143 | 1-483 (483) | 266-930 (930) | AlphaFold/UniProt50 (v4) |
| 19 | AF-A0A109FDE5-F1-model_v4 | DNA topoisomerase I | Rhodotorula sp. JG-1b | 1,00 | 40 | 5,24E-52 | 2105 | 2-483 (483) | 234-874 (874) | AlphaFold/UniProt50 (v4) |
| 20 | AF-A0A507D1T0-F1-model_v4 | DNA topoisomerase I | Synchytrium endobioticum | 1,00 | 41,4 | 6,50E-52 | 2089 | 1-483 (483) | 286-897 (897) | AlphaFold/UniProt50 (v4) |
| 21 | AF-A0A1U7LQ83-F1-model_v4 | DNA topoisomerase I | Neolecta irregularis DAH-3 | 1,00 | 42 | 7,25E-52 | 2123 | 1-483 (483) | 292-928 (928) | AlphaFold/UniProt50 (v4) |
| 22 | AF-P07799-F1-model_v4 | DNA topoisomerase 1 | Schizosaccharomyces pombe 972h- | 1,00 | 40,2 | 7,51E-52 | 2096 | 1-483 (483) | 181-814 (814) | AlphaFold/Swiss-Prot (v4) |
| 23 | AF-P07799-F1-model_v4 | DNA topoisomerase 1 | Schizosaccharomyces pombe 972h- | 1,00 | 40,2 | 8,08E-52 | 2096 | 1-483 (483) | 181-814 (814) | AlphaFold/Proteome (v4) |
| 24 | AF-A0A166XD30-F1-model_v4 | DNA topoisomerase I | Fibularhizoctonia sp. CBS 109695 | 1,00 | 39,2 | 9,00E-52 | 2117 | 1-483 (483) | 135-772 (772) | AlphaFold/UniProt50 (v4) |
| 25 | AF-A0A0J9XHU2-F1-model_v4 | DNA topoisomerase I | Geotrichum candidum | 1,00 | 40,2 | 1,25E-51 | 2126 | 1-483 (483) | 134-779 (779) | AlphaFold/UniProt50 (v4) |
| 26 | AF-A0A1S7HN26-F1-model_v4 | DNA topoisomerase I | Zygosaccharomyces parabailii | 1,00 | 40,3 | 1,39E-51 | 2086 | 1-483 (483) | 123-761 (761) | AlphaFold/UniProt50 (v4) |
| 27 | AF-A0A238F9S3-F1-model_v4 | DNA topoisomerase I | Microbotryum intermedium | 1,00 | 39,1 | 1,82E-51 | 2117 | 1-483 (483) | 420-1071 (1071) | AlphaFold/UniProt50 (v4) |
| 28 | AF-Q00313-F1-model_v4 | DNA topoisomerase 1 | Candida albicans | 1,00 | 39 | 2,22E-51 | 2051 | 1-483 (483) | 140-778 (778) | AlphaFold/Swiss-Prot (v4) |
| 29 | AF-Q6FVE9-F1-model_v4 | DNA topoisomerase I | [Candida] glabrata CBS 138 | 1,00 | 40,4 | 2,26E-51 | 2118 | 1-483 (483) | 79-717 (717) | AlphaFold/UniProt50 (v4) |
| 30 | AF-P07799-F1-model_v4 | DNA topoisomerase 1 | Schizosaccharomyces pombe 972h- | 1,00 | 40,2 | 2,97E-51 | 2096 | 1-483 (483) | 181-814 (814) | AlphaFold/UniProt50 (v4) |
| 31 | AF-U7Q7S5-F1-model_v4 | DNA topoisomerase I | Sporothrix schenckii ATCC 58251 | 1,00 | 39,7 | 3,13E-51 | 2073 | 1-483 (483) | 245-894 (894) | AlphaFold/Proteome (v4) |
| 32 | AF-W6MJD9-F1-model_v4 | DNA topoisomerase I | Kuraishia capsulata CBS 1993 | 1,00 | 40,6 | 5,10E-51 | 2082 | 3-483 (483) | 150-784 (784) | AlphaFold/UniProt50 (v4) |
| 33 | AF-A0A068S3R7-F1-model_v4 | DNA topoisomerase I | Lichtheimia corymbifera JMRC:FSU:9682 | 1,00 | 39,3 | 5,10E-51 | 2017 | 1-483 (483) | 152-804 (804) | AlphaFold/UniProt50 (v4) |
| 34 | AF-A0A1X2J2J0-F1-model_v4 | DNA topoisomerase I | Absidia repens | 1,00 | 39 | 6,34E-51 | 2060 | 1-483 (483) | 169-809 (809) | AlphaFold/UniProt50 (v4) |
| 35 | AF-F4RJG2-F1-model_v4 | DNA topoisomerase I | Melampsora larici-populina 98AG31 | 1,00 | 38,5 | 8,32E-51 | 2086 | 1-483 (483) | 299-937 (937) | AlphaFold/UniProt50 (v4) |
| 36 | AF-A0A4P9ZKD5-F1-model_v4 | DNA topoisomerase I | Dimargaris cristalligena | 1,00 | 39,2 | 8,32E-51 | 2050 | 1-478 (483) | 235-874 (875) | AlphaFold/UniProt50 (v4) |
| 37 | AF-A0A1S8W1J7-F1-model_v4 | DNA topoisomerase I | Batrachochytrium salamandrivorans | 1,00 | 40,3 | 8,32E-51 | 2049 | 1-482 (483) | 290-899 (899) | AlphaFold/UniProt50 (v4) |
| 38 | AF-Q00313-F1-model_v4 | DNA topoisomerase 1 | Candida albicans | 1,00 | 39 | 8,78E-51 | 2051 | 1-483 (483) | 140-778 (778) | AlphaFold/UniProt50 (v4) |
| 39 | AF-A7TJW1-F1-model_v4 | DNA topoisomerase I | Vanderwaltozyma polyspora DSM 70294 | 1,00 | 38,7 | 9,27E-51 | 2054 | 1-483 (483) | 161-799 (799) | AlphaFold/UniProt50 (v4) |
| 40 | AF-A0A1A0H6Y2-F1-model_v4 | DNA topoisomerase I | Metschnikowia bicuspidata var. bicuspidata NRRL YB-4993 | 1,00 | 40,1 | 1,03E-50 | 2082 | 1-483 (483) | 160-795 (795) | AlphaFold/UniProt50 (v4) |
| 41 | AF-A0A137P234-F1-model_v4 | DNA topoisomerase I | Conidiobolus coronatus NRRL 28638 | 1,00 | 37,6 | 1,03E-50 | 2055 | 1-483 (483) | 138-768 (768) | AlphaFold/UniProt50 (v4) |
| 42 | AF-A0A163JAM1-F1-model_v4 | DNA topoisomerase I | Absidia glauca | 1,00 | 39,9 | 1,09E-50 | 2038 | 13-483 (483) | 191-809 (809) | AlphaFold/UniProt50 (v4) |
| 43 | AF-A0A0L0P4F8-F1-model_v4 | DNA topoisomerase 1 | [Candida] auris | 1,00 | 39,6 | 1,13E-50 | 2050 | 1-483 (483) | 113-749 (749) | AlphaFold/Swiss-Prot (v4) |
| 44 | AF-A0A1Y2B9B9-F1-model_v4 | DNA topoisomerase I | Rhizoclosmatium globosum | 1,00 | 40,8 | 1,22E-50 | 2047 | 1-475 (483) | 190-796 (823) | AlphaFold/UniProt50 (v4) |
| 45 | AF-A0A642UVX3-F1-model_v4 | DNA topoisomerase I | Trichomonascus ciferrii | 1,00 | 38,7 | 1,28E-50 | 2076 | 1-483 (483) | 158-798 (798) | AlphaFold/UniProt50 (v4) |
| 46 | AF-A0A194SAX2-F1-model_v4 | DNA topoisomerase I | Rhodotorula graminis WP1 | 1,00 | 40,1 | 1,36E-50 | 2053 | 1-483 (483) | 106-736 (736) | AlphaFold/UniProt50 (v4) |
| 47 | AF-A0A1E5RRP9-F1-model_v4 | DNA topoisomerase I | Hanseniaspora uvarum | 1,00 | 39,4 | 1,68E-50 | 2063 | 1-483 (483) | 98-743 (743) | AlphaFold/UniProt50 (v4) |
| 48 | AF-Q6C7R2-F1-model_v4 | DNA topoisomerase I | Yarrowia lipolytica CLIB122 | 1,00 | 40,2 | 1,68E-50 | 2052 | 1-483 (483) | 121-760 (760) | AlphaFold/UniProt50 (v4) |
| 49 | AF-A0A286UHL7-F1-model_v4 | DNA topoisomerase I | Pyrrhoderma noxium | 1,00 | 38,9 | 1,78E-50 | 2084 | 1-483 (483) | 275-916 (916) | AlphaFold/UniProt50 (v4) |
| 50 | AF-A0A7D9CVB4-F1-model_v4 | DNA topoisomerase I | Brettanomyces bruxellensis | 1,00 | 39 | 1,78E-50 | 2080 | 1-483 (483) | 128-776 (776) | AlphaFold/UniProt50 (v4) |

**Supplementary table 1. Fungal proteins having structural similarities with human TOP1 as identified by Foldseek.**

| No. | Target | Description | Scientific name | Prob. | Seq. id. | E-value | Score | Query pos. | Target pos. | Database |
|---|---|---|---|---|---|---|---|---|---|---|
| 51 | AF-A0A427Y9D4-F1-model_v4 | DNA topoisomerase I | Apiotrichum porosum | 1,00 | 40,4 | 1,78E-50 | 2063 | 1-483 (483) | 206-844 (844) | AlphaFold/UniProt50 (v4) |
| 52 | AF-A0A1B2J5X6-F1-model_v4 | DNA topoisomerase I | Komagataella pastoris | 1,00 | 39 | 2,33E-50 | 2067 | 1-483 (483) | 125-761 (761) | AlphaFold/UniProt50 (v4) |
| 53 | AF-A0A0L0P4F8-F1-model_v4 | DNA topoisomerase 1 | [Candida] auris | 1,00 | 39,6 | 4,47E-50 | 2050 | 1-483 (483) | 113-749 (749) | AlphaFold/UniProt50 (v4) |
| 54 | AF-A0A2T0FFC8-F1-model_v4 | DNA topoisomerase I | Wickerhamiella sorbophila | 1,00 | 37,3 | 4,72E-50 | 2030 | 2-483 (483) | 99-746 (746) | AlphaFold/UniProt50 (v4) |
| 55 | AF-A0A1E3PTY7-F1-model_v4 | DNA topoisomerase I | Nadsonia fulvescens var. elongata DSM 6958 | 1,00 | 38,2 | 7,28E-50 | 2045 | 1-483 (483) | 140-785 (785) | AlphaFold/UniProt50 (v4) |
| 56 | AF-A0A2U9N8A9-F1-model_v4 | DNA topoisomerase I | Pichia kudriavzevii | 1,00 | 39 | 7,28E-50 | 2037 | 1-483 (483) | 81-728 (728) | AlphaFold/UniProt50 (v4) |
| 57 | AF-I4Y6D9-F1-model_v4 | DNA topoisomerase I | Wallemia mellicola CBS 633.66 | 1,00 | 36,7 | 8,11E-50 | 2021 | 1-483 (483) | 128-767 (767) | AlphaFold/UniProt50 (v4) |
| 58 | AF-A0A7R7VTK1-F1-model_v4 | DNA topoisomerase I | Aspergillus chevalieri | 1,00 | 36,8 | 8,56E-50 | 2041 | 2-483 (483) | 146-777 (777) | AlphaFold/UniProt50 (v4) |
| 59 | AF-A0A1E3PW82-F1-model_v4 | DNA topoisomerase I | Lipomyces starkeyi NRRL Y-11557 | 1,00 | 37,8 | 1,47E-49 | 2036 | 1-483 (483) | 146-781 (781) | AlphaFold/UniProt50 (v4) |
| 60 | AF-E4ZXG5-F1-model_v4 | DNA topoisomerase I | Leptosphaeria maculans JN3 | 1,00 | 39,5 | 1,64E-49 | 2019 | 1-483 (483) | 380-1025 (1025) | AlphaFold/UniProt50 (v4) |
| 61 | AF-A0A1R1YIE8-F1-model_v4 | DNA topoisomerase I | Smittium culicis | 1,00 | 37 | 1,64E-49 | 2008 | 1-483 (483) | 102-752 (752) | AlphaFold/UniProt50 (v4) |
| 62 | AF-A0A1D8NGA4-F1-model_v4 | DNA topoisomerase I | Yarrowia lipolytica | 1,00 | 40,3 | 2,98E-49 | 1998 | 1-483 (483) | 287-926 (926) | AlphaFold/UniProt50 (v4) |
| 63 | AF-A0A0F7SHA4-F1-model_v4 | DNA topoisomerase I | Phaffia rhodozyma | 1,00 | 38,2 | 3,15E-49 | 2033 | 1-483 (483) | 321-956 (956) | AlphaFold/UniProt50 (v4) |
| 64 | AF-K2SIS5-F1-model_v4 | DNA topoisomerase I | Macrophomina phaseolina MS6 | 1,00 | 39,8 | 3,15E-49 | 1980 | 1-483 (483) | 209-854 (854) | AlphaFold/UniProt50 (v4) |
| 65 | AF-A0A553IBY9-F1-model_v4 | DNA topoisomerase I | Xylaria flabelliformis | 1,00 | 37,6 | 3,51E-49 | 1993 | 1-483 (483) | 259-907 (907) | AlphaFold/UniProt50 (v4) |
| 66 | AF-G7E2E9-F1-model_v4 | DNA topoisomerase I | Mixia osmundae IAM 14324 | 1,00 | 37,5 | 3,70E-49 | 1976 | 3-483 (483) | 224-863 (863) | AlphaFold/UniProt50 (v4) |
| 67 | AF-A0A2G5HJX0-F1-model_v4 | DNA topoisomerase I | Cercospora beticola | 1,00 | 38 | 4,13E-49 | 2001 | 1-483 (483) | 241-886 (886) | AlphaFold/UniProt50 (v4) |
| 68 | AF-A0A0G2GDK1-F1-model_v4 | DNA topoisomerase I | Phaeomoniella chlamydospora | 1,00 | 38,3 | 4,60E-49 | 2011 | 1-483 (483) | 249-895 (1240) | AlphaFold/UniProt50 (v4) |
| 69 | AF-A0A060T5Z3-F1-model_v4 | DNA topoisomerase I | Blastobotrys adeninivorans | 1,00 | 38,4 | 5,41E-49 | 1987 | 3-483 (483) | 2-640 (640) | AlphaFold/UniProt50 (v4) |
| 70 | AF-A0A6A7BJR1-F1-model_v4 | DNA topoisomerase I | Plenodomus tracheiphilus IPT5 | 1,00 | 39,5 | 6,03E-49 | 2001 | 1-483 (483) | 382-1027 (1027) | AlphaFold/UniProt50 (v4) |
| 71 | AF-A0A292PTU7-F1-model_v4 | DNA topoisomerase I | Tuber aestivum | 1,00 | 38,9 | 6,72E-49 | 2002 | 1-483 (483) | 310-952 (952) | AlphaFold/UniProt50 (v4) |
| 72 | AF-A0A319B5I4-F1-model_v4 | DNA topoisomerase I | Aspergillus vadensis CBS 113365 | 1,00 | 38 | 8,82E-49 | 1990 | 1-483 (483) | 219-864 (864) | AlphaFold/UniProt50 (v4) |
| 73 | AF-A0A4T0C001-F1-model_v4 | DNA topoisomerase I | Aureobasidium pullulans | 1,00 | 38,7 | 1,29E-48 | 1986 | 1-483 (483) | 459-1104 (1104) | AlphaFold/UniProt50 (v4) |
| 74 | AF-A0A1B7NYS1-F1-model_v4 | DNA topoisomerase I | Emergomyces africanus | 1,00 | 39,2 | 1,36E-48 | 1974 | 1-483 (483) | 180-825 (825) | AlphaFold/UniProt50 (v4) |
| 75 | AF-W7LYN8-F1-model_v4 | DNA topoisomerase I | Fusarium verticillioides 7600 | 1,00 | 37 | 2,47E-48 | 1941 | 1-483 (483) | 168-823 (823) | AlphaFold/UniProt50 (v4) |
| 76 | AF-A0A5N6UHD0-F1-model_v4 | DNA topoisomerase I | Aspergillus tamarii | 1,00 | 37,5 | 2,61E-48 | 1970 | 1-483 (483) | 212-857 (857) | AlphaFold/UniProt50 (v4) |
| 77 | AF-A0A3M6YAQ4-F1-model_v4 | DNA topoisomerase I | Hortaea werneckii | 1,00 | 38 | 2,91E-48 | 1956 | 1-483 (483) | 181-826 (826) | AlphaFold/UniProt50 (v4) |
| 78 | AF-A0A175WIZ4-F1-model_v4 | DNA topoisomerase I | Madurella mycetomatis | 1,00 | 39,3 | 3,07E-48 | 1919 | 1-483 (483) | 262-911 (911) | AlphaFold/Proteome (v4) |
| 79 | AF-C0NC53-F1-model_v4 | DNA topoisomerase I | Histoplasma capsulatum G186AR | 1,00 | 39,6 | 4,02E-48 | 1930 | 1-483 (483) | 264-909 (909) | AlphaFold/Proteome (v4) |
| 80 | AF-B6JVY3-F1-model_v4 | DNA topoisomerase I | Schizosaccharomyces japonicus yFS275 | 1,00 | 38,5 | 4,25E-48 | 1350 | 1-483 (483) | 227-842 (842) | AlphaFold/UniProt50 (v4) |
| 81 | AF-A0A5N6Z2G8-F1-model_v4 | DNA topoisomerase I | Aspergillus coremiiformis | 1,00 | 37,5 | 4,48E-48 | 1957 | 1-483 (483) | 214-859 (859) | AlphaFold/UniProt50 (v4) |
| 82 | AF-Q0UUB0-F1-model_v4 | DNA topoisomerase I | Parastagonospora nodorum SN15 | 1,00 | 39,6 | 6,21E-48 | 1941 | 1-483 (483) | 242-887 (887) | AlphaFold/UniProt50 (v4) |
| 83 | AF-A0A0L1IP81-F1-model_v4 | DNA topoisomerase I | Aspergillus nomiae NRRL 13137 | 1,00 | 38 | 1,01E-47 | 1946 | 1-483 (483) | 269-914 (914) | AlphaFold/UniProt50 (v4) |
| 84 | AF-C1H484-F1-model_v4 | DNA topoisomerase I | Paracoccidioides lutzii Pb01 | 1,00 | 38,5 | 1,26E-47 | 1885 | 1-483 (483) | 261-906 (906) | AlphaFold/Proteome (v4) |
| 85 | AF-A0A4P9X2X7-F1-model_v4 | DNA topoisomerase I | Caulochytrium protostelioides | 1,00 | 35,9 | 1,33E-47 | 1894 | 12-483 (483) | 10-620 (620) | AlphaFold/UniProt50 (v4) |
| 86 | AF-A0A0D2CGJ0-F1-model_v4 | DNA topoisomerase I | Cladophialophora immunda | 1,00 | 38,7 | 1,56E-47 | 1926 | 3-483 (483) | 183-826 (826) | AlphaFold/UniProt50 (v4) |
| 87 | AF-A0A1C1CLM2-F1-model_v4 | DNA topoisomerase I | Cladophialophora carrionii | 1,00 | 38 | 9,86E-47 | 1864 | 2-483 (483) | 272-916 (916) | AlphaFold/Proteome (v4) |
| 88 | AF-A0A1Y1WKA6-F1-model_v4 | DNA topoisomerase I | Linderina pennispora | 1,00 | 38,4 | 1,29E-46 | 1598 | 1-483 (483) | 150-689 (689) | AlphaFold/UniProt50 (v4) |
| 89 | AF-A0A0D2GYT0-F1-model_v4 | DNA topoisomerase I | Fonsecaea pedrosoi CBS 271.37 | 1,00 | 39,1 | 1,70E-46 | 1842 | 2-483 (483) | 267-911 (911) | AlphaFold/Proteome (v4) |
| 90 | AF-A0A3M7ALR7-F1-model_v4 | DNA topoisomerase I | Hortaea werneckii | 1,00 | 44,5 | 5,90E-46 | 1670 | 1-482 (483) | 267-756 (784) | AlphaFold/UniProt50 (v4) |
| 91 | AF-X0MEW1-F1-model_v4 | DNA topoisomerase I | Fusarium oxysporum f. sp. vasinfectum 25433 | 1,00 | 48,6 | 8,17E-46 | 1771 | 3-437 (483) | 256-702 (719) | AlphaFold/UniProt50 (v4) |
| 92 | AF-A0A0J9UDZ2-F1-model_v4 | DNA topoisomerase I | Fusarium oxysporum f. sp. lycopersici 4287 | 1,00 | 48,2 | 1,84E-45 | 1753 | 3-437 (483) | 170-616 (633) | AlphaFold/UniProt50 (v4) |
| 93 | AF-A0A5C3QSY9-F1-model_v4 | DNA topoisomerase I | Pterula gracilis | 1,00 | 38,3 | 2,23E-44 | 1718 | 40-483 (483) | 3-597 (597) | AlphaFold/UniProt50 (v4) |
| 94 | AF-A0A0C3BEK4-F1-model_v4 | DNA topoisomerase I | Serendipita vermifera MAFF 305830 | 1,00 | 36,1 | 3,36E-43 | 1709 | 1-483 (483) | 158-805 (805) | AlphaFold/UniProt50 (v4) |
| 95 | AF-A0A1R1YBS7-F1-model_v4 | DNA topoisomerase I | Smittium culicis | 1,00 | 34,8 | 3,36E-43 | 1561 | 1-483 (483) | 143-758 (758) | AlphaFold/UniProt50 (v4) |
| 96 | AF-A0A1E4TDV2-F1-model_v4 | DNA topoisomerase 1 | Tortispora caseinolytica NRRL Y-17796 | 1,00 | 30,1 | 7,58E-43 | 1709 | 13-483 (483) | 98-729 (729) | AlphaFold/UniProt50 (v4) |
| 97 | AF-A0A1Y2HQW3-F1-model_v4 | DNA topoisomerase I | Catenaria anguillulae PL171 | 1,00 | 38,5 | 5,34E-42 | 1525 | 1-483 (483) | 232-797 (797) | AlphaFold/UniProt50 (v4) |
| 98 | AF-A0A316YQF7-F1-model_v4 | DNA topoisomerase 1 | Acaromyces ingoldii | 1,00 | 37 | 5,80E-41 | 1457 | 5-482 (483) | 160-680 (1056) | AlphaFold/UniProt50 (v4) |
| 99 | AF-A0A177UHQ3-F1-model_v4 | DNA topoisomerase 1 | Tilletia walkeri | 1,00 | 37 | 7,21E-41 | 1443 | 13-477 (483) | 180-696 (1249) | AlphaFold/UniProt50 (v4) |
| 100 | AF-A0A1Y2HSC9-F1-model_v4 | DNA topoisomerase 1 | Catenaria anguillulae PL171 | 1,00 | 32,7 | 8,03E-41 | 1474 | 2-483 (483) | 37-515 (515) | AlphaFold/UniProt50 (v4) |

**Supplementary table 1. Fungal proteins having structural similarities with human TOP1 as identified by Foldseek.**

| No. | Target | Description | Scientific name | Prob. | Seq. id. | E-value | Score | Query pos. | Target pos. | Database |
|---|---|---|---|---|---|---|---|---|---|---|
| 101 | AF-A0A066WDL8-F1-model_v4 | DNA topoisomerase 1 | Tilletiaria anomala UBC 951 | 1,00 | 34,3 | 2,13E-40 | 1403 | 5-482 (483) | 40-568 (942) | AlphaFold/UniProt50 (v4) |
| 102 | AF-A0A3G2RZ58-F1-model_v4 | DNA topoisomerase 1 | Malassezia restricta CBS 7877 | 1,00 | 34,2 | 4,81E-40 | 1410 | 5-478 (483) | 72-599 (838) | AlphaFold/UniProt50 (v4) |
| 103 | AF-A0A316VNB8-F1-model_v4 | DNA topoisomerase 1 | Meira miltonrushii | 1,00 | 36,1 | 4,81E-40 | 1400 | 6-478 (483) | 178-694 (988) | AlphaFold/UniProt50 (v4) |
| 104 | AF-A0A022Y0R9-F1-model_v4 | DNA topoisomerase I | Trichophyton soudanense CBS 452.61 | 1,00 | 38 | 5,97E-40 | 1485 | 61-483 (483) | 1-585 (585) | AlphaFold/UniProt50 (v4) |
| 105 | AF-A0A0F7RWP5-F1-model_v4 | DNA topoisomerase I | Sporisorium scitamineum | 1,00 | 34,4 | 8,73E-40 | 1390 | 13-482 (483) | 9-529 (862) | AlphaFold/UniProt50 (v4) |
| 106 | AF-X0IBP8-F1-model_v4 | DNA topoisomerase 1 | Fusarium oxysporum f. sp. conglutinans race 2 54008 | 1,00 | 46,9 | 2,32E-39 | 1540 | 3-384 (483) | 256-645 (647) | AlphaFold/UniProt50 (v4) |
| 107 | AF-X0AM58-F1-model_v4 | DNA topoisomerase 1 | Fusarium oxysporum f. sp. melonis 26406 | 1,00 | 46,3 | 2,88E-39 | 1503 | 3-385 (483) | 170-557 (571) | AlphaFold/UniProt50 (v4) |
| 108 | AF-P41511-F1-model_v4 | DNA topoisomerase 1 | Ustilago maydis 521 | 1,00 | 34,2 | 6,72E-39 | 1361 | 13-478 (483) | 161-679 (1019) | AlphaFold/Swiss-Prot (v4) |
| 109 | AF-A6R590-F1-model_v4 | DNA topoisomerase 1 | Histoplasma mississippiense (nom. inval.) | 1,00 | 48,4 | 6,85E-39 | 1542 | 1-369 (483) | 264-637 (651) | AlphaFold/UniProt50 (v4) |
| 110 | AF-A0A146I8R7-F1-model_v4 | DNA topoisomerase 1 | Mycena chlorophos | 1,00 | 37 | 7,24E-39 | 1449 | 3-483 (483) | 8-488 (488) | AlphaFold/UniProt50 (v4) |
| 111 | AF-P41511-F1-model_v4 | DNA topoisomerase 1 | Ustilago maydis 521 | 1,00 | 34,2 | 2,66E-38 | 1361 | 13-478 (483) | 161-679 (1019) | AlphaFold/UniProt50 (v4) |
| 112 | AF-A0A5B0QSF2-F1-model_v4 | DNA topoisomerase I | Puccinia graminis f. sp. tritici | 1,00 | 47 | 1,51E-37 | 1306 | 105-482 (483) | 3-382 (564) | AlphaFold/UniProt50 (v4) |
| 113 | AF-A0A317B5V2-F1-model_v4 | DNA topoisomerase I | Pyrenophora tritici-repentis | 1,00 | 40,3 | 2,15E-36 | 1245 | 138-483 (483) | 2-500 (500) | AlphaFold/UniProt50 (v4) |
| 114 | AF-A0A022WA16-F1-model_v4 | DNA topoisomerase I | Trichophyton rubrum CBS 288.86 | 1,00 | 45,9 | 2,27E-36 | 1297 | 65-465 (483) | 6-406 (415) | AlphaFold/UniProt50 (v4) |
| 115 | AF-H0H0X3-F1-model_v4 | DNA topoisomerase I | Saccharomyces cerevisiae x Saccharomyces kudriavzevii VIN7 | 1,00 | 50,1 | 1,36E-35 | 1227 | 105-480 (483) | 3-380 (455) | AlphaFold/UniProt50 (v4) |
| 116 | AF-A0A5M9K1I7-F1-model_v4 | DNA topoisomerase 1 | Monilinia fructicola | 1,00 | 38,6 | 6,91E-35 | 1101 | 1-482 (483) | 247-687 (930) | AlphaFold/UniProt50 (v4) |
| 117 | AF-E6Z7M9-F1-model_v4 | DNA topoisomerase I | Fusarium solani | 1,00 | 50,7 | 1,56E-32 | 1091 | 148-482 (483) | 3-340 (480) | AlphaFold/UniProt50 (v4) |
| 118 | AF-A0A137PFK3-F1-model_v4 | TOPEUc domain-containing protein | Conidiobolus coronatus NRRL 28638 | 1,00 | 24,3 | 3,17E-32 | 1049 | 2-482 (483) | 251-724 (871) | AlphaFold/UniProt50 (v4) |
| 119 | AF-A0A059JF26-F1-model_v4 | DNA topoisomerase 1 | Trichophyton interdigitale MR816 | 1,00 | 47,6 | 6,62E-29 | 1091 | 61-367 (483) | 1-311 (331) | AlphaFold/UniProt50 (v4) |
| 120 | AF-A0A3M6VWQ8-F1-model_v4 | DNA topoisomerase I | Hortaea werneckii | 1,00 | 36,9 | 7,63E-26 | 845 | 189-483 (483) | 1-448 (448) | AlphaFold/UniProt50 (v4) |
| 121 | AF-A0A0C3LE35-F1-model_v4 | DNA topoisomerase I | Tulasnella calospora MUT 4182 | 1,00 | 35,9 | 1,54E-25 | 826 | 190-483 (483) | 1-448 (458) | AlphaFold/UniProt50 (v4) |
| 122 | AF-A0A5C3E6N2-F1-model_v4 | DNA topoisomerase 1 | Ustilago trichophora | 1,00 | 28,9 | 1,15E-24 | 680 | 6-483 (483) | 153-604 (835) | AlphaFold/UniProt50 (v4) |
| 123 | AF-E2LN97-F1-model_v4 | DNA topoisomerase 1 | Moniliophthora perniciosa FA553 | 1,00 | 48 | 3,21E-24 | 919 | 36-305 (483) | 2-273 (343) | AlphaFold/UniProt50 (v4) |
| 124 | 1ois-assembly1_A | yeast dna topoisomerase i, n-terminal fragment | Saccharomyces cerevisiae | 1,00 | 50,4 | 2,19E-23 | 901 | 13-228 (483) | 3-219 (220) | PDB100 (20240101) |
| 125 | AF-A0A3M6YBS4-F1-model_v4 | TOPEUc domain-containing protein | Hortaea werneckii | 1,00 | 44 | 2,97E-23 | 950 | 1-240 (483) | 267-516 (517) | AlphaFold/UniProt50 (v4) |
| 126 | AF-A0A2N5S3X5-F1-model_v4 | DNA topoisomerase I | Puccinia coronata f. sp. avenae | 1,00 | 32 | 6,34E-23 | 746 | 223-483 (483) | 3-442 (442) | AlphaFold/UniProt50 (v4) |
| 127 | AF-A0A4U0VHA9-F1-model_v4 | DNA topoisomerase I | Friedmanniomyces endolithicus | 1,00 | 34,2 | 1,93E-21 | 702 | 226-483 (483) | 1-419 (419) | AlphaFold/UniProt50 (v4) |
| 128 | AF-A0A163K8K5-F1-model_v4 | DNA topoisomerase I | Absidia glauca | 1,00 | 34,2 | 2,22E-20 | 631 | 223-483 (483) | 243-634 (634) | AlphaFold/UniProt50 (v4) |
| 129 | AF-A0A4U0WHM5-F1-model_v4 | Fe2OG dioxygenase domain-containing protein | Cryomyces minteri | 1,00 | 45 | 3,07E-20 | 816 | 1-221 (483) | 632-862 (863) | AlphaFold/UniProt50 (v4) |
| 130 | AF-A0A2I1GB72-F1-model_v4 | TOPEUc domain-containing protein | Rhizophagus irregularis | 1,00 | 36,3 | 3,72E-19 | 611 | 265-483 (483) | 3-372 (372) | AlphaFold/UniProt50 (v4) |
| 131 | AF-A0A1C7LRH2-F1-model_v4 | DNA topoisomerase 1 | Grifola frondosa | 1,00 | 54 | 3,35E-17 | 688 | 256-427 (483) | 11-176 (302) | AlphaFold/UniProt50 (v4) |
| 132 | AF-A0A1Y2EMU7-F1-model_v4 | DNA topoisomerase 1 | Neocallimastix californiae | 1,00 | 25 | 2,63E-16 | 366 | 3-483 (483) | 199-568 (568) | AlphaFold/UniProt50 (v4) |
| 133 | AF-A0A3M6XV51-F1-model_v4 | TOPEUc domain-containing protein | Hortaea werneckii | 1,00 | 32,1 | 1,08E-15 | 480 | 268-483 (483) | 1-385 (385) | AlphaFold/UniProt50 (v4) |
| 134 | AF-E2LGK1-F1-model_v4 | Topoisom_I_N domain-containing protein | Moniliophthora perniciosa FA553 | 1,00 | 43,1 | 3,46E-14 | 550 | 1-194 (483) | 70-262 (265) | AlphaFold/UniProt50 (v4) |
| 135 | 1oisA02 | Topoisomerase I | Saccharomyces cerevisiae S288C | 1,00 | 58,3 | 1,58E-13 | 501 | 121-228 (483) | 20-127 (128) | CATH50 (v4.3.0) |
| 136 | AF-A0A2I1GB49-F1-model_v4 | Topoisom_I_N domain-containing protein | Rhizophagus irregularis | 1,00 | 45,3 | 8,04E-13 | 490 | 1-176 (483) | 145-321 (324) | AlphaFold/UniProt50 (v4) |
| 137 | AF-M5BLW1-F1-model_v4 | DNA topoisomerase I | Rhizoctonia solani AG-1 IB | 1,00 | 57,9 | 1,11E-12 | 537 | 240-383 (483) | 2-140 (140) | AlphaFold/UniProt50 (v4) |
| 138 | AF-M5BKG0-F1-model_v4 | DNA topoisomerase I | Rhizoctonia solani AG-1 IB | 1,00 | 45,3 | 1,72E-12 | 502 | 1-175 (483) | 280-458 (461) | AlphaFold/UniProt50 (v4) |
| 139 | AF-A0A017SG24-F1-model_v4 | TOPEUc domain-containing protein | Aspergillus ruber CBS 135680 | 1,00 | 39,8 | 2,80E-12 | 341 | 268-483 (483) | 1-198 (241) | AlphaFold/UniProt50 (v4) |
| 140 | AF-A0A0L0TE33-F1-model_v4 | DNA topoisomerase 1 | Allomyces macrogynus ATCC 38327 | 1,00 | 24,2 | 7,84E-12 | 321 | 1-483 (483) | 179-593 (593) | AlphaFold/UniProt50 (v4) |
| 141 | AF-A0A017SF00-F1-model_v4 | Topoisom_I_N domain-containing protein | Aspergillus ruber CBS 135680 | 1,00 | 28,9 | 2,21E-07 | 270 | 2-199 (483) | 96-278 (278) | AlphaFold/UniProt50 (v4) |
| 142 | AF-A0A0D1YYX2-F1-model_v4 | TOPEUc domain-containing protein | Exophiala sideris | 1,00 | 32,1 | 4,48E-07 | 201 | 248-421 (483) | 180-379 (410) | AlphaFold/UniProt50 (v4) |
| 143 | AF-Q8WZF2-F1-model_v4 | Topoisomerase I | Cryptococcus neoformans | 1,00 | 39,5 | 1,10E-05 | 168 | 327-482 (483) | 1-159 (270) | AlphaFold/UniProt50 (v4) |
| 144 | AF-A0A4U0V000-F1-model_v4 | TOPEUc domain-containing protein | Cryomyces minteri | 1,00 | 27,4 | 1,29E-05 | 180 | 342-483 (483) | 1-301 (301) | AlphaFold/UniProt50 (v4) |
| 145 | 1oisA01 | Topoisomerase I | Saccharomyces cerevisiae S288C | 1,00 | 41,6 | 1,26E-04 | 193 | 30-117 (483) | 1-92 (92) | CATH50 (v4.3.0) |

No. = number, prob. = probability of target being homologous to human TOP1, seq. id. = sequence identity, E-value = expect value, score = structural bit score, pos. = position.

**Supplementary table 2. Bacterial proteins having structural similarities with human TOP1 as identified by Foldseek.**

| No. | Target | Description | Scientific name | Prob. | Seq. id. | E-value | Score | Query pos. | Target pos. | Database |
|---|---|---|---|---|---|---|---|---|---|---|
| 1 | AF-A0A399ZYW8-F1-model_v4 | DNA topoisomerase I | Chloroflexi bacterium | 1,00 | 25,3 | 3,71E-34 | 1261 | 13-474 (483) | 1-605 (618) | AlphaFold/UniProt50 (v4) |
| 2 | AF-A0A7C6XLZ8-F1-model_v4 | DNA topoisomerase I | Chloroflexi bacterium | 1,00 | 25 | 2,22E-33 | 1232 | 15-473 (483) | 2-572 (745) | AlphaFold/UniProt50 (v4) |
| 3 | AF-A0A7C3R1Z9-F1-model_v4 | DNA topoisomerase I | Chloroflexi bacterium | 1,00 | 30,3 | 6,57E-33 | 1196 | 15-430 (483) | 2-419 (543) | AlphaFold/UniProt50 (v4) |
| 4 | AF-A0A7V5YXE7-F1-model_v4 | DNA topoisomerase I | Anaerolineae bacterium | 1,00 | 29,3 | 1,85E-30 | 1134 | 15-409 (483) | 2-396 (399) | AlphaFold/UniProt50 (v4) |
| 5 | AF-A0A7C7C7X9-F1-model_v4 | DNA topoisomerase I | Candidatus Marinimicrobia bacterium | 1,00 | 32,9 | 9,40E-30 | 1061 | 56-437 (483) | 2-385 (403) | AlphaFold/UniProt50 (v4) |
| 6 | AF-A0A7C6TRA0-F1-model_v4 | TOPEUc domain-containing protein | Chloroflexi bacterium | 1,00 | 26,7 | 4,70E-24 | 770 | 149-473 (483) | 1-438 (607) | AlphaFold/UniProt50 (v4) |
| 7 | AF-A0A420Z5G6-F1-model_v4 | TOPEUc domain-containing protein | Candidatus Woesebacteria bacterium | 1,00 | 32,5 | 1,39E-23 | 792 | 2-329 (483) | 24-370 (370) | AlphaFold/UniProt50 (v4) |
| 8 | AF-A0A1W9UTK5-F1-model_v4 | TOPEUc domain-containing protein | Anaerolineaceae bacterium 4572_32.1 | 1,00 | 28,5 | 1,57E-17 | 567 | 210-475 (483) | 2-383 (385) | AlphaFold/UniProt50 (v4) |
| 9 | AF-G8RL29-F1-model_v4 | Topoisomerase IB | Mycolicibacterium rhodesiae NBB3 | 1,00 | 17,8 | 1,63E-12 | 366 | 147-475 (483) | 2-330 (338) | AlphaFold/UniProt50 (v4) |
| 10 | AF-A0A542SEE0-F1-model_v4 | DNA topoisomerase IB | Nocardioides sp. SLBN-35 | 1,00 | 18,9 | 1,81E-12 | 383 | 153-473 (483) | 10-314 (317) | AlphaFold/UniProt50 (v4) |
| 11 | AF-A0A0G1W433-F1-model_v4 | Topoisomerase IB | Candidatus Gottesmanbacteria bacterium GW2011_GWB1_49_7 | 1,00 | 21,2 | 2,65E-12 | 419 | 208-450 (483) | 78-300 (306) | AlphaFold/UniProt50 (v4) |
| 12 | AF-A0A7C1ZDP6-F1-model_v4 | Topoisom_I domain-containing protein | bacterium | 1,00 | 15,2 | 3,87E-12 | 316 | 6-470 (483) | 21-443 (503) | AlphaFold/UniProt50 (v4) |
| 13 | AF-A0A4P8KRI5-F1-model_v4 | DNA topoisomerase IB | Microbacterium sp. RG1 | 1,00 | 20,7 | 4,09E-12 | 402 | 194-473 (483) | 48-318 (320) | AlphaFold/UniProt50 (v4) |
| 14 | AF-U2XA09-F1-model_v4 | Topoisomerase IB | Microbacterium sp. TS-1 | 1,00 | 19,4 | 4,81E-12 | 409 | 207-474 (483) | 61-319 (320) | AlphaFold/UniProt50 (v4) |
| 15 | AF-A0A100ZN21-F1-model_v4 | DNA topoisomerase | Mycobacterium sp. GA-1999 | 1,00 | 17 | 5,08E-12 | 401 | 207-475 (483) | 60-330 (339) | AlphaFold/UniProt50 (v4) |
| 16 | AF-A0A7X0GDH2-F1-model_v4 | DNA topoisomerase-1 | Arthrobacter sp. AZCC_0090 | 1,00 | 18,7 | 5,36E-12 | 418 | 209-476 (483) | 30-284 (285) | AlphaFold/UniProt50 (v4) |
| 17 | AF-A0A1F7X2Q9-F1-model_v4 | Topoisom_I domain-containing protein | Candidatus Woesebacteria bacterium RBG_13_36_22 | 1,00 | 22,6 | 6,31E-12 | 443 | 209-452 (483) | 91-318 (322) | AlphaFold/UniProt50 (v4) |
| 18 | AF-A0A3C1CKK9-F1-model_v4 | DNA topoisomerase | Acidimicrobiaceae bacterium | 1,00 | 17,2 | 7,42E-12 | 356 | 147-475 (483) | 2-324 (354) | AlphaFold/UniProt50 (v4) |
| 19 | AF-A0A1Y5LFN8-F1-model_v4 | DNA topoisomerase | Rhodococcus sp. NCIMB 12038 | 1,00 | 18,9 | 8,28E-12 | 390 | 204-474 (483) | 58-329 (335) | AlphaFold/UniProt50 (v4) |
| 20 | AF-A0A7W3MHG4-F1-model_v4 | DNA topoisomerase-1 | Sphingobacterium soli | 1,00 | 20 | 9,22E-12 | 394 | 209-475 (483) | 75-336 (344) | AlphaFold/UniProt50 (v4) |
| 21 | AF-A0A7I7VPB8-F1-model_v4 | DNA topoisomerase | Mycobacterium doricum | 1,00 | 17 | 1,59E-11 | 433 | 207-448 (483) | 60-296 (302) | AlphaFold/UniProt50 (v4) |
| 22 | AF-D5UCE1-F1-model_v4 | Putative viral-like DNA topoisomerase | Cellulomonas flavigena DSM 20109 | 1,00 | 17 | 1,68E-11 | 404 | 209-475 (483) | 63-326 (330) | AlphaFold/UniProt50 (v4) |
| 23 | AF-A0A0Q8QF49-F1-model_v4 | DNA topoisomerase | Nocardioides sp. Root190 | 1,00 | 18,2 | 1,68E-11 | 392 | 209-474 (483) | 62-319 (325) | AlphaFold/UniProt50 (v4) |
| 24 | AF-U1LCK5-F1-model_v4 | Topoisom_I domain-containing protein | Agrococcus pavilionensis RW1 | 1,00 | 17,5 | 1,87E-11 | 398 | 207-473 (483) | 61-319 (320) | AlphaFold/UniProt50 (v4) |
| 25 | AF-A0A3A5M242-F1-model_v4 | DNA topoisomerase IB | Arthrobacter cheniae | 1,00 | 19,8 | 1,87E-11 | 349 | 144-473 (483) | 2-319 (324) | AlphaFold/UniProt50 (v4) |
| 26 | AF-A0A1Q4UH71-F1-model_v4 | DNA topoisomerase | Mycobacterium sp. ST-F2 | 1,00 | 17,7 | 2,08E-11 | 342 | 142-475 (483) | 20-327 (336) | AlphaFold/UniProt50 (v4) |
| 27 | AF-A0A7X5TVF5-F1-model_v4 | DNA topoisomerase-1 | Lysinibacter cavernae | 1,00 | 17,2 | 2,20E-11 | 403 | 207-474 (483) | 60-322 (326) | AlphaFold/UniProt50 (v4) |
| 28 | AF-A0A2E0KX27-F1-model_v4 | DNA topoisomerase I | Anaerolineaceae bacterium | 1,00 | 17,2 | 2,20E-11 | 390 | 205-475 (483) | 66-337 (339) | AlphaFold/UniProt50 (v4) |
| 29 | AF-A0A2S9ABI1-F1-model_v4 | Topoisomerase I | Microbacterium sp. MYb72 | 1,00 | 17,5 | 2,32E-11 | 416 | 205-452 (483) | 59-300 (319) | AlphaFold/UniProt50 (v4) |
| 30 | AF-A0A7Y9XAJ7-F1-model_v4 | DNA topoisomerase IB | Nocardiopsis sinuspersici | 1,00 | 18 | 2,32E-11 | 347 | 155-475 (483) | 12-336 (340) | AlphaFold/UniProt50 (v4) |
| 31 | AF-V7KWC6-F1-model_v4 | DNA topoisomerase | Mycobacterium avium subsp. paratuberculosis 08-8281 | 1,00 | 17 | 2,32E-11 | 333 | 146-475 (483) | 51-378 (399) | AlphaFold/UniProt50 (v4) |
| 32 | AF-A0A7V9AS46-F1-model_v4 | DNA topoisomerase IB | Thermoleophilaceae bacterium | 1,00 | 16,3 | 2,45E-11 | 382 | 208-479 (483) | 30-307 (313) | AlphaFold/UniProt50 (v4) |
| 33 | AF-A0A6N8KWR6-F1-model_v4 | DNA topoisomerase IB | Sphingobacterium humi | 1,00 | 19,5 | 2,45E-11 | 379 | 209-473 (483) | 101-365 (372) | AlphaFold/UniProt50 (v4) |
| 34 | AF-A0A2V7YSE1-F1-model_v4 | Topoisom_I domain-containing protein | Acidobacteria bacterium | 1,00 | 17,9 | 2,88E-11 | 394 | 208-475 (483) | 315-578 (590) | AlphaFold/UniProt50 (v4) |
| 35 | AF-A0A0N0GNT4-F1-model_v4 | Eukaryotic DNA topoisomerase I, catalytic core | Amantichitinum ursilacus | 1,00 | 16,4 | 3,04E-11 | 344 | 156-475 (483) | 54-365 (374) | AlphaFold/UniProt50 (v4) |
| 36 | AF-A0A367XY99-F1-model_v4 | DNA topoisomerase IB | Microbacterium sorbitolivorans | 1,00 | 17,1 | 3,04E-11 | 342 | 147-478 (483) | 20-341 (341) | AlphaFold/UniProt50 (v4) |
| 37 | AF-A0A2W5Z4V8-F1-model_v4 | DNA topoisomerase | Solirubrobacterales bacterium | 1,00 | 18,2 | 3,39E-11 | 322 | 146-483 (483) | 4-344 (358) | AlphaFold/UniProt50 (v4) |
| 38 | AF-A0A6I4NUZ9-F1-model_v4 | DNA topoisomerase IB | Agromyces sp. MMS17-SY077 | 1,00 | 16,4 | 3,58E-11 | 382 | 205-477 (483) | 59-323 (323) | AlphaFold/UniProt50 (v4) |
| 39 | AF-A0A0B2CHR5-F1-model_v4 | DNA topoisomerase | Xanthomonas cannabis pv. cannabis | 1,00 | 17,9 | 3,58E-11 | 331 | 156-475 (483) | 60-373 (393) | AlphaFold/UniProt50 (v4) |
| 40 | AF-A0A7Y9MC46-F1-model_v4 | DNA topoisomerase-1 | Leifsonia sp. AK011 | 1,00 | 18,9 | 3,99E-11 | 377 | 194-472 (483) | 47-315 (318) | AlphaFold/UniProt50 (v4) |
| 41 | AF-A0A3M9MQ19-F1-model_v4 | DNA topoisomerase IB | Rufibacter immobilis | 1,00 | 17,4 | 3,99E-11 | 370 | 207-474 (483) | 80-350 (355) | AlphaFold/UniProt50 (v4) |
| 42 | AF-K0EWH0-F1-model_v4 | DNA topoisomerase | Nocardia brasiliensis ATCC 700358 | 1,00 | 16,1 | 4,21E-11 | 327 | 154-475 (483) | 7-330 (336) | AlphaFold/Proteome (v4) |
| 43 | AF-A0A4Q6C944-F1-model_v4 | DNA topoisomerase IB | Proteobacteria bacterium | 1,00 | 18,8 | 4,21E-11 | 335 | 155-475 (483) | 45-364 (374) | AlphaFold/UniProt50 (v4) |
| 44 | AF-A0A1B4I0E4-F1-model_v4 | DNA topoisomerase | Burkholderia metallica | 1,00 | 17,6 | 4,44E-11 | 336 | 156-475 (483) | 9-323 (331) | AlphaFold/UniProt50 (v4) |
| 45 | AF-A0A4Q2JNI7-F1-model_v4 | DNA topoisomerase IB | Agromyces fucosus | 1,00 | 18,9 | 4,69E-11 | 382 | 208-475 (483) | 52-311 (316) | AlphaFold/UniProt50 (v4) |
| 46 | AF-A0A7Y0LXD4-F1-model_v4 | DNA topoisomerase IB | Cellulomonas fimi | 1,00 | 17,4 | 4,95E-11 | 330 | 147-473 (483) | 3-325 (364) | AlphaFold/UniProt50 (v4) |
| 47 | AF-A0A7Z2JGA1-F1-model_v4 | DNA topoisomerase IB | Paraburkholderia acidisoli | 1,00 | 19,1 | 5,23E-11 | 376 | 209-475 (483) | 104-367 (381) | AlphaFold/UniProt50 (v4) |
| 48 | AF-A0A2S6B4D6-F1-model_v4 | DNA topoisomerase | Pseudoxanthomonas sp. KAs_5_3 | 1,00 | 21,4 | 5,52E-11 | 359 | 194-475 (483) | 48-322 (331) | AlphaFold/UniProt50 (v4) |
| 49 | AF-A0A1D9FCR1-F1-model_v4 | Topoisomerase I | Arthrobacter sp. ZXY-2 | 1,00 | 18 | 5,52E-11 | 357 | 149-475 (483) | 7-315 (318) | AlphaFold/UniProt50 (v4) |
| 50 | AF-A0A7I7JVH4-F1-model_v4 | DNA topoisomerase | Mycobacterium novum | 1,00 | 15,8 | 5,83E-11 | 396 | 207-464 (483) | 59-313 (378) | AlphaFold/UniProt50 (v4) |

**Supplementary table 2. Bacterial proteins having structural similarities with human TOP1 as identified by Foldseek.**

| No. | Target | Description | Scientific name | Prob. | Seq. id. | E-value | Score | Query pos. | Target pos. | Database |
|---|---|---|---|---|---|---|---|---|---|---|
| 51 | AF-A0A1G7U768-F1-model_v4 | DNA topoisomerase IB | Lentzea fradiae | 1,00 | 17,7 | 5,83E-11 | 384 | 204-476 (483) | 57-307 (307) | AlphaFold/UniProt50 (v4) |
| 52 | AF-A0A7X0FL06-F1-model_v4 | DNA topoisomerase IB | Xanthomonas sacchari | 1,00 | 16,3 | 5,83E-11 | 364 | 207-475 (483) | 85-349 (409) | AlphaFold/UniProt50 (v4) |
| 53 | AF-A0A3G8V364-F1-model_v4 | Topoisomerase I | Microbacterium sp. Y-01 | 1,00 | 17,5 | 6,50E-11 | 332 | 155-471 (483) | 12-319 (337) | AlphaFold/UniProt50 (v4) |
| 54 | AF-A0A3A5AJC9-F1-model_v4 | DNA topoisomerase IB | Desulfobacteraceae bacterium | 1,00 | 16,4 | 6,86E-11 | 376 | 209-475 (483) | 72-339 (342) | AlphaFold/UniProt50 (v4) |
| 55 | AF-A0A6N4RBJ9-F1-model_v4 | DNA topoisomerase IB | Blastochloris viridis | 1,00 | 19,5 | 6,86E-11 | 376 | 207-475 (483) | 85-351 (353) | AlphaFold/UniProt50 (v4) |
| 56 | AF-A0A0Q5LC00-F1-model_v4 | Topoisomerase I | Microbacterium sp. Leaf161 | 1,00 | 17,9 | 6,86E-11 | 355 | 153-452 (483) | 9-299 (319) | AlphaFold/UniProt50 (v4) |
| 57 | AF-A0A4Q7RTZ8-F1-model_v4 | DNA topoisomerase IB | Stenotrophomonas sp. BK441 | 1,00 | 20,3 | 7,64E-11 | 372 | 209-475 (483) | 76-337 (351) | AlphaFold/UniProt50 (v4) |
| 58 | AF-A0A7W9FHS0-F1-model_v4 | DNA topoisomerase-1 | Micrococcus sp. TA1 | 1,00 | 15,2 | 7,64E-11 | 316 | 153-473 (483) | 7-347 (356) | AlphaFold/UniProt50 (v4) |
| 59 | AF-A0A1Q2CSG5-F1-model_v4 | Topoisom_I domain-containing protein | Tessaracoccus aquimaris | 1,00 | 18,1 | 8,07E-11 | 381 | 209-471 (483) | 64-320 (325) | AlphaFold/UniProt50 (v4) |
| 60 | AF-A0A2V8LSQ8-F1-model_v4 | DNA topoisomerase I | Acidobacteria bacterium | 1,00 | 17,5 | 8,07E-11 | 379 | 209-476 (483) | 53-310 (310) | AlphaFold/UniProt50 (v4) |
| 61 | AF-A0A0Q5PNC3-F1-model_v4 | Topoisomerase I | Microbacterium sp. Leaf179 | 1,00 | 17,2 | 8,07E-11 | 373 | 194-476 (483) | 48-321 (321) | AlphaFold/UniProt50 (v4) |
| 62 | AF-A0A1G8FFU9-F1-model_v4 | DNA topoisomerase-1 | Agrococcus jejuensis | 1,00 | 15,9 | 8,07E-11 | 344 | 153-472 (483) | 10-317 (319) | AlphaFold/UniProt50 (v4) |
| 63 | AF-A0A6N7JTW8-F1-model_v4 | DNA topoisomerase IB | Stenotrophomonas sp. MYb238 | 1,00 | 16,6 | 8,52E-11 | 368 | 207-475 (483) | 62-325 (332) | AlphaFold/UniProt50 (v4) |
| 64 | AF-A0A7Y8WXH4-F1-model_v4 | DNA topoisomerase IB | Streptomyces sp. BR123 | 1,00 | 18,2 | 8,52E-11 | 361 | 209-475 (483) | 94-366 (373) | AlphaFold/UniProt50 (v4) |
| 65 | AF-A0A2S8M188-F1-model_v4 | DNA topoisomerase | Mycobacterium sp. ITM-2016-00317 | 1,00 | 15,4 | 8,99E-11 | 364 | 208-475 (483) | 15-286 (294) | AlphaFold/UniProt50 (v4) |
| 66 | AF-A0A7W1URS5-F1-model_v4 | DNA topoisomerase IB | Acidimicrobiia bacterium | 1,00 | 17,7 | 9,49E-11 | 319 | 152-475 (483) | 22-342 (344) | AlphaFold/UniProt50 (v4) |
| 67 | AF-A0A7W5IHG4-F1-model_v4 | DNA topoisomerase-1 | Paraburkholderia sp. WP4_3_2 | 1,00 | 18,2 | 1,00E-10 | 369 | 207-474 (483) | 79-342 (401) | AlphaFold/UniProt50 (v4) |
| 68 | AF-A0A369QE86-F1-model_v4 | DNA topoisomerase | Adhaeribacter pallidiroseus | 1,00 | 16,7 | 1,00E-10 | 351 | 190-476 (483) | 119-397 (401) | AlphaFold/UniProt50 (v4) |
| 69 | AF-A0A5N0TEZ0-F1-model_v4 | DNA topoisomerase IB | Microbacterium caowuchunii | 1,00 | 19 | 1,06E-10 | 354 | 194-473 (483) | 48-318 (320) | AlphaFold/UniProt50 (v4) |
| 70 | AF-A0A0Q8F1E4-F1-model_v4 | DNA topoisomerase | Pseudoxanthomonas sp. Root65 | 1,00 | 19,4 | 1,12E-10 | 329 | 156-475 (483) | 12-324 (331) | AlphaFold/UniProt50 (v4) |
| 71 | AF-A0A7J9WHB1-F1-model_v4 | DNA topoisomerase IB | Nitriliruptorales bacterium | 1,00 | 17,4 | 1,12E-10 | 319 | 147-475 (483) | 3-336 (355) | AlphaFold/UniProt50 (v4) |
| 72 | AF-A0A562CWX4-F1-model_v4 | DNA topoisomerase IB | Nocardioides sp. J9 | 1,00 | 18 | 1,18E-10 | 361 | 208-474 (483) | 38-300 (306) | AlphaFold/UniProt50 (v4) |
| 73 | AF-A0A1W9LYD6-F1-model_v4 | Topoisom_I domain-containing protein | Desulfobulbaceae bacterium A2 | 1,00 | 16,7 | 1,18E-10 | 341 | 192-455 (483) | 530-761 (764) | AlphaFold/UniProt50 (v4) |
| 74 | AF-A0A538KHE6-F1-model_v4 | DNA topoisomerase IB | Actinomycetia bacterium | 1,00 | 19,2 | 1,25E-10 | 369 | 220-475 (483) | 11-271 (276) | AlphaFold/UniProt50 (v4) |
| 75 | AF-A0A1H5XHC1-F1-model_v4 | DNA topoisomerase-1 | Sphingobacterium lactis | 1,00 | 21,1 | 1,25E-10 | 347 | 209-475 (483) | 69-331 (340) | AlphaFold/UniProt50 (v4) |
| 76 | AF-A0A1F5C1X4-F1-model_v4 | Topoisom_I domain-containing protein | Candidatus Atribacteria bacterium RBG_16_35_8 | 1,00 | 18,5 | 1,32E-10 | 349 | 210-460 (483) | 105-351 (355) | AlphaFold/UniProt50 (v4) |
| 77 | AF-A0A1Q8BKJ6-F1-model_v4 | DNA topoisomerase I | Cyanobacteria bacterium 13_1_20CM_4_61_6 | 1,00 | 18,8 | 1,47E-10 | 383 | 208-446 (483) | 68-300 (303) | AlphaFold/UniProt50 (v4) |
| 78 | AF-A0A5C7PW81-F1-model_v4 | DNA topoisomerase IB | Desulfurellales bacterium | 1,00 | 17,7 | 1,47E-10 | 379 | 208-450 (483) | 93-321 (323) | AlphaFold/UniProt50 (v4) |
| 79 | AF-A0A4P2PVF0-F1-model_v4 | DNA topoisomerase I | Sorangium cellulosum | 1,00 | 15,1 | 1,55E-10 | 366 | 207-475 (483) | 129-401 (453) | AlphaFold/UniProt50 (v4) |
| 80 | AF-A0A6P2BUD7-F1-model_v4 | DNA topoisomerase IB | Trebonia kvetii | 1,00 | 17,7 | 1,72E-10 | 360 | 209-482 (483) | 74-347 (353) | AlphaFold/UniProt50 (v4) |
| 81 | AF-A0A7W0M938-F1-model_v4 | DNA topoisomerase IB | Planctomycetaceae bacterium | 1,00 | 17,5 | 1,72E-10 | 348 | 204-475 (483) | 83-371 (378) | AlphaFold/UniProt50 (v4) |
| 82 | AF-A0A6P2JH63-F1-model_v4 | DNA topoisomerase | Burkholderia territorii | 1,00 | 16,7 | 1,72E-10 | 315 | 148-475 (483) | 64-386 (398) | AlphaFold/UniProt50 (v4) |
| 83 | AF-A0A1R1L7B9-F1-model_v4 | Topoisom_I domain-containing protein | Tersicoccus phoenicis | 1,00 | 17,6 | 1,82E-10 | 369 | 207-472 (483) | 66-328 (332) | AlphaFold/UniProt50 (v4) |
| 84 | AF-A0A7X7QEI1-F1-model_v4 | DNA topoisomerase IB | Actinomycetales bacterium | 1,00 | 15 | 1,82E-10 | 349 | 209-475 (483) | 23-313 (317) | AlphaFold/UniProt50 (v4) |
| 85 | AF-A0A1H7WG08-F1-model_v4 | DNA topoisomerase IB | Pseudoxanthomonas sp. GM95 | 1,00 | 17,3 | 1,92E-10 | 351 | 205-475 (483) | 88-354 (375) | AlphaFold/UniProt50 (v4) |
| 86 | AF-A0A2N7VCZ3-F1-model_v4 | DNA topoisomerase | Trinickia dabaoshanensis | 1,00 | 16 | 1,92E-10 | 304 | 155-482 (483) | 45-345 (431) | AlphaFold/UniProt50 (v4) |
| 87 | AF-A0A0U4FM98-F1-model_v4 | Topoisom_I domain-containing protein | Microbacterium sp. XT11 | 1,00 | 17,7 | 2,03E-10 | 387 | 208-452 (483) | 62-300 (368) | AlphaFold/UniProt50 (v4) |
| 88 | AF-A0A5M8B5X1-F1-model_v4 | DNA topoisomerase IB | Cupriavidus cauae | 1,00 | 15,1 | 2,03E-10 | 363 | 209-475 (483) | 64-329 (338) | AlphaFold/UniProt50 (v4) |
| 89 | AF-A0A7Y9J8X4-F1-model_v4 | DNA topoisomerase IB | Actinomycetospora corticicola | 1,00 | 16,3 | 2,03E-10 | 308 | 154-475 (483) | 9-343 (349) | AlphaFold/UniProt50 (v4) |
| 90 | AF-A0A7Z6TT41-F1-model_v4 | DNA topoisomerase | Streptomyces sp. ZS0098 | 1,00 | 15,6 | 2,26E-10 | 354 | 207-483 (483) | 60-345 (345) | AlphaFold/UniProt50 (v4) |
| 91 | AF-A0A1X7FDS4-F1-model_v4 | DNA topoisomerase-1 | Trinickia caryophylli | 1,00 | 17,1 | 2,26E-10 | 311 | 156-475 (483) | 53-368 (381) | AlphaFold/UniProt50 (v4) |
| 92 | AF-A0A1Q7IXZ0-F1-model_v4 | DNA topoisomerase | Betaproteobacteria bacterium 13_1_40CM_4_64_4 | 1,00 | 16,7 | 2,26E-10 | 299 | 156-475 (483) | 37-352 (383) | AlphaFold/UniProt50 (v4) |
| 93 | AF-A0A0Q5IMH0-F1-model_v4 | DNA topoisomerase | Xanthomonas sp. Leaf148 | 1,00 | 18,5 | 2,39E-10 | 356 | 207-475 (483) | 109-373 (407) | AlphaFold/UniProt50 (v4) |
| 94 | AF-A0A2V8ND24-F1-model_v4 | DNA topoisomerase I | Acidobacteria bacterium | 1,00 | 19 | 2,52E-10 | 356 | 208-473 (483) | 26-295 (318) | AlphaFold/UniProt50 (v4) |
| 95 | AF-A0A4Q3AIB1-F1-model_v4 | DNA topoisomerase IB | Verrucomicrobiaceae bacterium | 1,00 | 17,3 | 2,52E-10 | 340 | 207-476 (483) | 125-400 (431) | AlphaFold/UniProt50 (v4) |
| 96 | AF-A0A6G7Z4C1-F1-model_v4 | DNA topoisomerase IB | Sanguibacter sp. HDW7 | 1,00 | 17,6 | 2,52E-10 | 329 | 149-475 (483) | 7-317 (322) | AlphaFold/UniProt50 (v4) |
| 97 | AF-A0A7W9JDQ3-F1-model_v4 | DNA topoisomerase IB | Kribbella italica | 1,00 | 18,1 | 2,52E-10 | 304 | 143-475 (483) | 32-361 (369) | AlphaFold/UniProt50 (v4) |
| 98 | AF-A0A653WC74-F1-model_v4 | DNA topoisomerase | Pseudoclavibacter sp. 8L | 1,00 | 19,3 | 2,66E-10 | 380 | 204-452 (483) | 59-301 (325) | AlphaFold/UniProt50 (v4) |
| 99 | AF-A0A1W6Z464-F1-model_v4 | DNA topoisomerase | Bordetella genomosp. 9 | 1,00 | 17,4 | 2,66E-10 | 353 | 209-473 (483) | 91-355 (386) | AlphaFold/UniProt50 (v4) |
| 100 | AF-A0A6L9G719-F1-model_v4 | DNA topoisomerase IB | Glutamicibacter soli | 1,00 | 17,7 | 2,66E-10 | 351 | 208-473 (483) | 56-324 (328) | AlphaFold/UniProt50 (v4) |

**Supplementary table 2. Bacterial proteins having structural similarities with human TOP1 as identified by Foldseek.**

| No. | Target | Description | Scientific name | Prob. | Seq. id. | E-value | Score | Query pos. | Target pos. | Database |
|---|---|---|---|---|---|---|---|---|---|---|
| 101 | AF-A0A356FHK5-F1-model_v4 | DNA topoisomerase IB | Janibacter terrae | 1,00 | 18,3 | 2,66E-10 | 330 | 153-476 (483) | 7-328 (328) | AlphaFold/UniProt50 (v4) |
| 102 | AF-A4AFI7-F1-model_v4 | Topoisom_I domain-containing protein | marine actinobacterium PHSC20C1 | 1,00 | 18,5 | 2,81E-10 | 359 | 204-475 (483) | 59-316 (324) | AlphaFold/UniProt50 (v4) |
| 103 | AF-A0A7W6QS02-F1-model_v4 | DNA topoisomerase-1 | Rhodoblastus acidophilus | 1,00 | 17,2 | 2,81E-10 | 354 | 207-473 (483) | 83-345 (352) | AlphaFold/UniProt50 (v4) |
| 104 | AF-A0A0Q8QWX8-F1-model_v4 | Topoisomerase I | Sphingomonas sp. Root720 | 1,00 | 17,6 | 2,97E-10 | 336 | 192-477 (483) | 51-319 (319) | AlphaFold/UniProt50 (v4) |
| 105 | AF-A0A7D7WIA9-F1-model_v4 | DNA topoisomerase IB | Devosia sp. D6-9 | 1,00 | 16,7 | 3,13E-10 | 335 | 191-475 (483) | 54-321 (323) | AlphaFold/UniProt50 (v4) |
| 106 | AF-A0A4Y8KHM6-F1-model_v4 | DNA topoisomerase IB | Cryobacterium sp. Hb1 | 1,00 | 16,5 | 3,13E-10 | 320 | 204-478 (483) | 59-369 (369) | AlphaFold/UniProt50 (v4) |
| 107 | AF-A0A2V3V3J2-F1-model_v4 | DNA topoisomerase-1 | Blastomonas natatoria | 1,00 | 17,9 | 3,30E-10 | 365 | 207-473 (483) | 64-320 (327) | AlphaFold/UniProt50 (v4) |
| 108 | AF-A0A372JXH7-F1-model_v4 | DNA topoisomerase IB | Paraburkholderia sp. DHOC27 | 1,00 | 18,4 | 3,30E-10 | 348 | 207-475 (483) | 89-354 (453) | AlphaFold/UniProt50 (v4) |
| 109 | AF-A0A2R7KD04-F1-model_v4 | DNA topoisomerase | Pseudomonas sp. HMWF006 | 1,00 | 19 | 3,30E-10 | 321 | 155-434 (483) | 21-293 (298) | AlphaFold/UniProt50 (v4) |
| 110 | AF-A0A358IPV8-F1-model_v4 | DNA topoisomerase | Micrococcaceae bacterium | 1,00 | 15,6 | 3,49E-10 | 380 | 208-446 (483) | 62-287 (329) | AlphaFold/UniProt50 (v4) |
| 111 | AF-A0A0U5B8E2-F1-model_v4 | Topoisom_I domain-containing protein | Microcella alkaliphila | 1,00 | 21,5 | 3,49E-10 | 342 | 205-469 (483) | 45-282 (304) | AlphaFold/UniProt50 (v4) |
| 112 | AF-A0A5A7NRC4-F1-model_v4 | Topoisom_I domain-containing protein | Zafaria cholistanensis | 1,00 | 17,3 | 3,49E-10 | 323 | 153-473 (483) | 6-315 (319) | AlphaFold/UniProt50 (v4) |
| 113 | AF-A0A1G5S96-F1-model_v4 | DNA topoisomerase IB | Nitrosospira sp. Nsp18 | 1,00 | 15 | 3,68E-10 | 346 | 207-475 (483) | 145-411 (428) | AlphaFold/UniProt50 (v4) |
| 114 | AF-A7HB17-F1-model_v4 | Type I topoisomerase | Anaeromyxobacter sp. Fw109-5 | 1,00 | 16,2 | 3,68E-10 | 336 | 208-473 (483) | 60-332 (376) | AlphaFold/UniProt50 (v4) |
| 115 | AF-A0A537WIL5-F1-model_v4 | DNA topoisomerase IB | Actinomycetia bacterium | 1,00 | 17,6 | 3,68E-10 | 293 | 154-475 (483) | 142-465 (479) | AlphaFold/UniProt50 (v4) |
| 116 | AF-A0A7T6XA37-F1-model_v4 | Topoisom_I domain-containing protein | Ralstonia pickettii | 1,00 | 15,8 | 4,10E-10 | 311 | 160-473 (483) | 63-374 (392) | AlphaFold/UniProt50 (v4) |
| 117 | AF-A0A2U1F8G7-F1-model_v4 | DNA topoisomerase IB | Actinomycetospora cinnamomea | 1,00 | 15,3 | 4,10E-10 | 296 | 153-475 (483) | 1-342 (360) | AlphaFold/UniProt50 (v4) |
| 118 | AF-A0A7V9JQS0-F1-model_v4 | DNA topoisomerase IB | Actinomycetia bacterium | 1,00 | 15,6 | 4,33E-10 | 359 | 207-473 (483) | 203-474 (492) | AlphaFold/UniProt50 (v4) |
| 119 | AF-A0A4P7DVQ4-F1-model_v4 | DNA topoisomerase IB | Sphingobacterium sp. CZ-2 | 1,00 | 17,5 | 5,10E-10 | 342 | 194-475 (483) | 60-337 (346) | AlphaFold/UniProt50 (v4) |
| 120 | AF-M3CDN7-F1-model_v4 | DNA topoisomerase | Streptomyces mobaraensis NBRC 13819 = DSM 40847 | 1,00 | 15,9 | 5,10E-10 | 339 | 190-475 (483) | 101-375 (390) | AlphaFold/UniProt50 (v4) |
| 121 | AF-A0A1U9NSK3-F1-model_v4 | DNA topoisomerase | Streptomyces sp. fd1-xmd | 1,00 | 19 | 5,10E-10 | 330 | 207-475 (483) | 59-333 (340) | AlphaFold/UniProt50 (v4) |
| 122 | AF-A0A437MJR6-F1-model_v4 | DNA topoisomerase IB | Rhodovarius crocodyli | 1,00 | 15,2 | 5,38E-10 | 366 | 203-446 (483) | 64-301 (343) | AlphaFold/UniProt50 (v4) |
| 123 | AF-A0A4R6ECH5-F1-model_v4 | DNA topoisomerase-1 | Azoarcus indigens | 1,00 | 15,1 | 5,38E-10 | 343 | 208-475 (483) | 63-327 (392) | AlphaFold/UniProt50 (v4) |
| 124 | AF-A0A1Q8BPN1-F1-model_v4 | DNA topoisomerase I | Cyanobacteria bacterium 13_1_20CM_4_61_6 | 1,00 | 18,6 | 5,68E-10 | 368 | 208-452 (483) | 42-279 (285) | AlphaFold/UniProt50 (v4) |
| 125 | AF-A0A7U3ZLP8-F1-model_v4 | DNA topoisomerase | Runella slithyformis DSM 19594 | 1,00 | 17,7 | 5,68E-10 | 341 | 205-475 (483) | 89-366 (368) | AlphaFold/UniProt50 (v4) |
| 126 | AF-A0A2S0N8F9-F1-model_v4 | DNA topoisomerase I | Phreatobacter cathodiphilus | 1,00 | 18 | 5,68E-10 | 331 | 194-472 (483) | 47-329 (366) | AlphaFold/UniProt50 (v4) |
| 127 | AF-A0A2V8R255-F1-model_v4 | DNA topoisomerase I | Acidobacteria bacterium | 1,00 | 18 | 6,00E-10 | 340 | 208-464 (483) | 107-362 (414) | AlphaFold/UniProt50 (v4) |
| 128 | AF-A0A172Y5Q3-F1-model_v4 | DNA topoisomerase | Brevundimonas naejangsanensis | 1,00 | 15,3 | 6,33E-10 | 349 | 209-474 (483) | 60-321 (322) | AlphaFold/UniProt50 (v4) |
| 129 | AF-A0A848WZB5-F1-model_v4 | DNA topoisomerase IB | Verrucomicrobiales bacterium | 1,00 | 19,2 | 6,33E-10 | 345 | 207-473 (483) | 63-329 (340) | AlphaFold/UniProt50 (v4) |
| 130 | AF-A0A1H6X2J1-F1-model_v4 | DNA topoisomerase IB | Frateuria terrea | 1,00 | 16 | 6,69E-10 | 344 | 207-473 (483) | 114-378 (392) | AlphaFold/UniProt50 (v4) |
| 131 | AF-A0A259U0M0-F1-model_v4 | Topoisom_I domain-containing protein | Rubricoccus marinus | 1,00 | 16,6 | 7,06E-10 | 342 | 209-475 (483) | 74-337 (341) | AlphaFold/UniProt50 (v4) |
| 132 | AF-A0A840KG51-F1-model_v4 | DNA topoisomerase-1 | Chryseobacterium defluvii | 1,00 | 15,4 | 7,06E-10 | 334 | 209-483 (483) | 32-307 (307) | AlphaFold/UniProt50 (v4) |
| 133 | AF-A0A2V8RAS2-F1-model_v4 | DNA topoisomerase I | Acidobacteria bacterium | 1,00 | 17,1 | 7,06E-10 | 327 | 208-473 (483) | 107-379 (383) | AlphaFold/UniProt50 (v4) |
| 134 | AF-A0A7X6PNL9-F1-model_v4 | DNA topoisomerase IB | Corynebacterium humireducens | 1,00 | 17,7 | 7,06E-10 | 324 | 209-475 (483) | 76-346 (350) | AlphaFold/UniProt50 (v4) |
| 135 | AF-A0A7W5YE09-F1-model_v4 | DNA topoisomerase IB | Nonomuraea dietziae | 1,00 | 17,3 | 7,06E-10 | 318 | 153-476 (483) | 7-303 (303) | AlphaFold/UniProt50 (v4) |
| 136 | AF-A0A1X0ZZD0-F1-model_v4 | DNA topoisomerase | Pseudomonas putida | 1,00 | 15,8 | 7,45E-10 | 347 | 209-475 (483) | 69-332 (336) | AlphaFold/UniProt50 (v4) |
| 137 | AF-A0A1G1TH49-F1-model_v4 | DNA topoisomerase I | Hymenobacter coccineus | 1,00 | 16,7 | 7,45E-10 | 334 | 202-477 (483) | 73-352 (352) | AlphaFold/UniProt50 (v4) |
| 138 | AF-A0A1H7QZ13-F1-model_v4 | DNA topoisomerase IB | Nitrosovibrio tenuis | 1,00 | 16,3 | 7,45E-10 | 317 | 194-476 (483) | 68-345 (345) | AlphaFold/UniProt50 (v4) |
| 139 | AF-A0A520PPQ7-F1-model_v4 | Topoisom_I domain-containing protein | Proteobacteria bacterium | 1,00 | 14,3 | 7,45E-10 | 295 | 136-446 (483) | 62-335 (345) | AlphaFold/UniProt50 (v4) |
| 140 | AF-A0A0Q7D325-F1-model_v4 | DNA topoisomerase | Phenylobacterium sp. Root1277 | 1,00 | 17,9 | 7,87E-10 | 338 | 207-473 (483) | 70-328 (339) | AlphaFold/UniProt50 (v4) |
| 141 | AF-A0A150R7M3-F1-model_v4 | DNA topoisomerase | Sorangium cellulosum | 1,00 | 16,3 | 8,77E-10 | 344 | 207-472 (483) | 131-401 (423) | AlphaFold/UniProt50 (v4) |
| 142 | AF-A0A349B1L3-F1-model_v4 | DNA topoisomerase | Acidimicrobiaceae bacterium | 1,00 | 15 | 8,77E-10 | 285 | 146-476 (483) | 18-327 (338) | AlphaFold/UniProt50 (v4) |
| 143 | AF-A0A239LT13-F1-model_v4 | DNA topoisomerase-1 | Granulicella rosea | 1,00 | 19,3 | 9,26E-10 | 337 | 207-473 (483) | 91-360 (375) | AlphaFold/UniProt50 (v4) |
| 144 | AF-A0A4Q2XAB2-F1-model_v4 | DNA topoisomerase IB | Verrucomicrobiaceae bacterium | 1,00 | 18,4 | 9,26E-10 | 337 | 207-473 (483) | 117-390 (392) | AlphaFold/UniProt50 (v4) |
| 145 | AF-A0A365YG39-F1-model_v4 | DNA topoisomerase | Glutamicibacter soli | 1,00 | 17,7 | 1,03E-09 | 337 | 208-473 (483) | 104-372 (376) | AlphaFold/UniProt50 (v4) |
| 146 | AF-Q0RIW9-F1-model_v4 | Putative type I DNA topoisomerase | Frankia alni ACN14a | 1,00 | 16,7 | 1,03E-09 | 327 | 208-475 (483) | 24-297 (318) | AlphaFold/UniProt50 (v4) |
| 147 | AF-A0A519YYY8-F1-model_v4 | DNA topoisomerase IB | Rubrivivax sp. | 1,00 | 17 | 1,03E-09 | 323 | 208-475 (483) | 13-280 (290) | AlphaFold/UniProt50 (v4) |
| 148 | AF-A0A2T0VBI7-F1-model_v4 | DNA topoisomerase-1 | Glaciihabitans tibetensis | 1,00 | 15,4 | 1,09E-09 | 346 | 208-475 (483) | 49-301 (309) | AlphaFold/UniProt50 (v4) |
| 149 | AF-A0A1G5N651-F1-model_v4 | DNA topoisomerase-1 | Afifella marina DSM 2698 | 1,00 | 15,9 | 1,09E-09 | 330 | 207-475 (483) | 70-335 (354) | AlphaFold/UniProt50 (v4) |
| 150 | AF-A0A2W5TKQ9-F1-model_v4 | DNA topoisomerase I | Archangium gephyra | 1,00 | 15,5 | 1,15E-09 | 339 | 209-474 (483) | 78-342 (353) | AlphaFold/UniProt50 (v4) |

**Supplementary table 2. Bacterial proteins having structural similarities with human TOP1 as identified by Foldseek.**

| No. | Target | Description | Scientific name | Prob. | Seq. id. | E-value | Score | Query pos. | Target pos. | Database |
|---|---|---|---|---|---|---|---|---|---|---|
| 151 | AF-A0A0L8C270-F1-model_v4 | DNA topoisomerase | Ensifer adhaerens | 1,00 | 17,3 | 1,15E-09 | 314 | 209-475 (483) | 64-328 (338) | AlphaFold/UniProt50 (v4) |
| 152 | AF-M7N2V3-F1-model_v4 | Eukaryotic DNA topoisomerase I, catalytic core | Cesiribacter andamanensis AMV16 | 1,00 | 16,9 | 1,15E-09 | 296 | 190-474 (483) | 74-349 (373) | AlphaFold/UniProt50 (v4) |
| 153 | AF-A0A1H9K5D3-F1-model_v4 | DNA topoisomerase IB | Actinokineospora terrae | 1,00 | 16,6 | 1,21E-09 | 341 | 208-475 (483) | 43-299 (303) | AlphaFold/UniProt50 (v4) |
| 154 | AF-A0A2D8Q593-F1-model_v4 | Topoisomerase I | Tistrella sp. | 1,00 | 15,6 | 1,21E-09 | 329 | 207-474 (483) | 63-332 (338) | AlphaFold/UniProt50 (v4) |
| 155 | AF-A0A3A1U1I9-F1-model_v4 | DNA topoisomerase IB | Amnibacterium setariae | 1,00 | 18,6 | 1,21E-09 | 329 | 204-483 (483) | 78-349 (369) | AlphaFold/UniProt50 (v4) |
| 156 | AF-A0A433JY05-F1-model_v4 | DNA topoisomerase IB | Legionella sp. km772 | 1,00 | 17,5 | 1,28E-09 | 331 | 209-473 (483) | 41-301 (312) | AlphaFold/UniProt50 (v4) |
| 157 | AF-A0A1N7B2T4-F1-model_v4 | DNA topoisomerase-1 | Rhizobium sp. RU20A | 1,00 | 17,6 | 1,28E-09 | 323 | 194-469 (483) | 53-313 (336) | AlphaFold/UniProt50 (v4) |
| 158 | AF-A0A6N6JJ61-F1-model_v4 | DNA topoisomerase | Litoreibacter roseus | 1,00 | 17,4 | 1,28E-09 | 310 | 208-474 (483) | 16-276 (282) | AlphaFold/UniProt50 (v4) |
| 159 | AF-A0A1C6SL32-F1-model_v4 | DNA topoisomerase IB | Micromonospora inyonensis | 1,00 | 17,3 | 1,35E-09 | 339 | 207-476 (483) | 60-327 (327) | AlphaFold/UniProt50 (v4) |
| 160 | AF-A0A2D7CNN2-F1-model_v4 | Topoisom_I domain-containing protein | Rhodospirillaceae bacterium | 1,00 | 20 | 1,35E-09 | 319 | 205-482 (483) | 71-330 (330) | AlphaFold/UniProt50 (v4) |
| 161 | AF-A0A7G9RVJ6-F1-model_v4 | DNA topoisomerase IB | Diaphorobacter ruginosibacter | 1,00 | 17,8 | 1,35E-09 | 315 | 208-474 (483) | 52-315 (326) | AlphaFold/UniProt50 (v4) |
| 162 | AF-A0A536SS95-F1-model_v4 | DNA topoisomerase IB | Betaproteobacteria bacterium | 1,00 | 17,5 | 1,43E-09 | 344 | 205-434 (483) | 73-298 (334) | AlphaFold/UniProt50 (v4) |
| 163 | AF-A0A2U9PM35-F1-model_v4 | Type I topoisomerase | Mycolicibacterium smegmatis MKD8 | 1,00 | 16,8 | 1,43E-09 | 340 | 209-473 (483) | 62-329 (338) | AlphaFold/UniProt50 (v4) |
| 164 | AF-A0A1G9EQA0-F1-model_v4 | DNA topoisomerase-1 | Methylophilus rhizosphaerae | 1,00 | 18,8 | 1,43E-09 | 337 | 207-474 (483) | 70-337 (351) | AlphaFold/UniProt50 (v4) |
| 165 | AF-A0A7Y9I3Q2-F1-model_v4 | DNA topoisomerase IB | Microlunatus parietis | 1,00 | 15,3 | 1,43E-09 | 331 | 209-475 (483) | 62-330 (333) | AlphaFold/UniProt50 (v4) |
| 166 | AF-A0A7V2IN07-F1-model_v4 | Topoisom_I domain-containing protein | Phycisphaerales bacterium | 1,00 | 15,7 | 1,43E-09 | 259 | 158-483 (483) | 255-623 (633) | AlphaFold/UniProt50 (v4) |
| 167 | AF-A0A1H9FDD7-F1-model_v4 | DNA topoisomerase-1 | Chelativorans sp. A52C2 | 1,00 | 17,6 | 1,51E-09 | 332 | 207-475 (483) | 71-336 (382) | AlphaFold/UniProt50 (v4) |
| 168 | AF-A0A1K1RIF1-F1-model_v4 | DNA topoisomerase-1 | Chitinophaga sancti | 1,00 | 17,8 | 1,51E-09 | 329 | 208-475 (483) | 86-354 (361) | AlphaFold/UniProt50 (v4) |
| 169 | AF-A0A4V2MPQ1-F1-model_v4 | DNA topoisomerase IB | Oricola cellulosilytica | 1,00 | 18,5 | 1,59E-09 | 321 | 209-475 (483) | 78-336 (359) | AlphaFold/UniProt50 (v4) |
| 170 | AF-A0A5P9QC81-F1-model_v4 | DNA topoisomerase | Luteimicrobium xylanilyticum | 1,00 | 15,9 | 1,59E-09 | 319 | 207-475 (483) | 59-337 (344) | AlphaFold/UniProt50 (v4) |
| 171 | AF-A0A7V9EZD2-F1-model_v4 | DNA topoisomerase IB | Thermoleophilaceae bacterium | 1,00 | 19,7 | 1,59E-09 | 314 | 213-475 (483) | 4-271 (286) | AlphaFold/UniProt50 (v4) |
| 172 | AF-A0A354TWB3-F1-model_v4 | DNA topoisomerase | Hyphomonadaceae bacterium | 1,00 | 16,9 | 1,59E-09 | 303 | 194-476 (483) | 61-326 (326) | AlphaFold/UniProt50 (v4) |
| 173 | AF-A0A4Q3V729-F1-model_v4 | DNA topoisomerase IB | bacterium | 1,00 | 17,5 | 1,68E-09 | 340 | 207-474 (483) | 70-330 (339) | AlphaFold/UniProt50 (v4) |
| 174 | AF-A0A2V3X046-F1-model_v4 | DNA topoisomerase-1 | Paraburkholderia tropica | 1,00 | 17,7 | 1,68E-09 | 336 | 207-474 (483) | 86-348 (362) | AlphaFold/UniProt50 (v4) |
| 175 | AF-A0A7V8YWS3-F1-model_v4 | DNA topoisomerase IB | Blastocatellia bacterium | 1,00 | 15,3 | 1,77E-09 | 315 | 191-476 (483) | 68-350 (351) | AlphaFold/UniProt50 (v4) |
| 176 | AF-A0A6L9MUZ8-F1-model_v4 | DNA topoisomerase IB | Alteromonas hispanica | 1,00 | 20,1 | 1,77E-09 | 267 | 194-475 (483) | 47-321 (325) | AlphaFold/UniProt50 (v4) |
| 177 | AF-Q9I1M5-F1-model_v4 | Topoisom_I domain-containing protein | Pseudomonas aeruginosa PAO1 | 1,00 | 17,3 | 1,87E-09 | 317 | 209-473 (483) | 74-329 (333) | AlphaFold/Proteome (v4) |
| 178 | AF-A0A2V6DKN9-F1-model_v4 | DNA topoisomerase I | Verrucomicrobia bacterium | 1,00 | 16,5 | 1,87E-09 | 339 | 208-473 (483) | 53-319 (336) | AlphaFold/UniProt50 (v4) |
| 179 | AF-A0A7J9ZWN6-F1-model_v4 | MFS transporter | Streptosporangiales bacterium | 1,00 | 18,1 | 1,87E-09 | 330 | 209-475 (483) | 323-595 (604) | AlphaFold/UniProt50 (v4) |
| 180 | AF-B5H816-F1-model_v4 | DNA topoisomerase I | Streptomyces pristinaespiralis ATCC 25486 | 1,00 | 16,6 | 1,87E-09 | 309 | 208-483 (483) | 9-294 (476) | AlphaFold/UniProt50 (v4) |
| 181 | AF-A0A521R5I1-F1-model_v4 | DNA topoisomerase IB | Reyranella sp. | 1,00 | 17,5 | 1,98E-09 | 329 | 209-473 (483) | 130-386 (393) | AlphaFold/UniProt50 (v4) |
| 182 | AF-A0A6N7CXX8-F1-model_v4 | Topoisom_I domain-containing protein | Rhodococcus sp. T7 | 1,00 | 18,1 | 1,98E-09 | 328 | 252-474 (483) | 8-236 (242) | AlphaFold/UniProt50 (v4) |
| 183 | AF-A0A346NLP3-F1-model_v4 | DNA topoisomerase IB | Salinimonas sediminis | 1,00 | 14,8 | 1,98E-09 | 296 | 208-476 (483) | 56-332 (332) | AlphaFold/UniProt50 (v4) |
| 184 | AF-A0A2D4SFN3-F1-model_v4 | DNA topoisomerase I | Salinisphaeraceae bacterium | 1,00 | 15,8 | 1,98E-09 | 288 | 156-475 (483) | 27-343 (349) | AlphaFold/UniProt50 (v4) |
| 185 | AF-A0A368EMV1-F1-model_v4 | DNA topoisomerase IB | Microbacterium sp. | 1,00 | 18,8 | 2,21E-09 | 335 | 192-443 (483) | 47-293 (293) | AlphaFold/UniProt50 (v4) |
| 186 | AF-A0A172X6F0-F1-model_v4 | Topoisom_I domain-containing protein | Leifsonia xyli | 1,00 | 15,7 | 2,21E-09 | 304 | 208-475 (483) | 65-343 (346) | AlphaFold/UniProt50 (v4) |
| 187 | AF-A0A5S9P0W2-F1-model_v4 | Topoisom_I domain-containing protein | Starkeya sp. HF14-78462 | 1,00 | 16,7 | 2,33E-09 | 327 | 207-483 (483) | 73-342 (343) | AlphaFold/UniProt50 (v4) |
| 188 | AF-A0A2S9CA75-F1-model_v4 | DNA topoisomerase | Arthrobacter sp. MYb214 | 1,00 | 18,4 | 2,46E-09 | 327 | 208-473 (483) | 118-388 (392) | AlphaFold/UniProt50 (v4) |
| 189 | AF-A0A3A0DVQ5-F1-model_v4 | DNA topoisomerase | Planctomycetota bacterium | 1,00 | 15,8 | 2,46E-09 | 313 | 205-473 (483) | 72-338 (353) | AlphaFold/UniProt50 (v4) |
| 190 | AF-A0A0S9E0B4-F1-model_v4 | Topoisom_I domain-containing protein | Agreia sp. Leaf244 | 1,00 | 13,7 | 2,59E-09 | 307 | 208-475 (483) | 64-342 (359) | AlphaFold/UniProt50 (v4) |
| 191 | AF-A0A7X6HDF1-F1-model_v4 | DNA topoisomerase IB | Arthrobacter mobilis | 1,00 | 19,7 | 2,59E-09 | 299 | 207-471 (483) | 61-354 (355) | AlphaFold/UniProt50 (v4) |
| 192 | AF-A0A1I4IL58-F1-model_v4 | DNA topoisomerase-1 | Leifsonia sp. CL147 | 1,00 | 18 | 2,74E-09 | 347 | 194-452 (483) | 48-300 (323) | AlphaFold/UniProt50 (v4) |
| 193 | AF-A0A6G7BVF3-F1-model_v4 | DNA topoisomerase IB | Sphingobacterium sp. DR205 | 1,00 | 15,1 | 2,89E-09 | 339 | 208-474 (483) | 57-316 (321) | AlphaFold/UniProt50 (v4) |
| 194 | AF-A0A7T7XJ13-F1-model_v4 | DNA topoisomerase IB | Luteimonas sp. MC1750 | 1,00 | 15,9 | 2,89E-09 | 317 | 207-473 (483) | 82-344 (366) | AlphaFold/UniProt50 (v4) |
| 195 | AF-A0A1G7YUD6-F1-model_v4 | DNA topoisomerase-1 | Bradyrhizobium sp. Rc2d | 1,00 | 14,1 | 2,89E-09 | 316 | 209-475 (483) | 175-440 (444) | AlphaFold/UniProt50 (v4) |
| 196 | AF-A0A847DST1-F1-model_v4 | DNA topoisomerase IB | Limnobacter sp. | 1,00 | 17,4 | 2,89E-09 | 300 | 207-478 (483) | 66-344 (373) | AlphaFold/UniProt50 (v4) |
| 197 | AF-A0A363RRI6-F1-model_v4 | DNA topoisomerase | Comamonas sp. JNW | 1,00 | 16,9 | 3,05E-09 | 301 | 207-474 (483) | 65-330 (340) | AlphaFold/UniProt50 (v4) |
| 198 | AF-A0A536VDF1-F1-model_v4 | DNA topoisomerase IB | Betaproteobacteria bacterium | 1,00 | 18,7 | 3,40E-09 | 336 | 205-427 (483) | 73-291 (293) | AlphaFold/UniProt50 (v4) |
| 199 | AF-A0A3M5V5S3-F1-model_v4 | Topoisom_I domain-containing protein | Pseudomonas syringae pv. avii | 1,00 | 16,2 | 3,40E-09 | 271 | 149-483 (483) | 222-558 (560) | AlphaFold/UniProt50 (v4) |
| 200 | AF-A0A2R8AAX6-F1-model_v4 | Topoisom_I domain-containing protein | Pontivivens insulae | 1,00 | 15,9 | 3,59E-09 | 311 | 208-473 (483) | 77-336 (337) | AlphaFold/UniProt50 (v4) |

**Supplementary table 2. Bacterial proteins having structural similarities with human TOP1 as identified by Foldseek.**

| No. | Target | Description | Scientific name | Prob. | Seq. id. | E-value | Score | Query pos. | Target pos. | Database |
|---|---|---|---|---|---|---|---|---|---|---|
| 201 | AF-A0A1P8JZB8-F1-model_v4 | DNA topoisomerase | Rhodoferax koreense | 1,00 | 15,5 | 3,59E-09 | 251 | 149-473 (483) | 10-347 (399) | AlphaFold/UniProt50 (v4) |
| 202 | AF-A0A7Y6URU3-F1-model_v4 | DNA topoisomerase IB | Ensifer sp. HO-A22 | 1,00 | 19 | 3,79E-09 | 330 | 209-446 (483) | 70-297 (344) | AlphaFold/UniProt50 (v4) |
| 203 | AF-A0A7K2KWZ2-F1-model_v4 | DNA topoisomerase IB | Streptomyces sp. SID6041 | 1,00 | 18,7 | 3,79E-09 | 320 | 236-472 (483) | 15-265 (271) | AlphaFold/UniProt50 (v4) |
| 204 | AF-A0A059DX67-F1-model_v4 | Topoisom_I domain-containing protein | Hyphomonas atlantica | 1,00 | 18 | 3,79E-09 | 309 | 207-474 (483) | 78-323 (329) | AlphaFold/UniProt50 (v4) |
| 205 | AF-Q5NXZ3-F1-model_v4 | Type I topoisomerase | Aromatoleum aromaticum EbN1 | 1,00 | 14,6 | 4,00E-09 | 311 | 208-474 (483) | 62-329 (381) | AlphaFold/UniProt50 (v4) |
| 206 | AF-A0A2S3YJ69-F1-model_v4 | DNA topoisomerase | Sinorhizobium americanum | 1,00 | 16,1 | 4,00E-09 | 285 | 194-475 (483) | 70-346 (374) | AlphaFold/UniProt50 (v4) |
| 207 | AF-A0A2N5KCB6-F1-model_v4 | DNA topoisomerase I | Candidatus Saccharibacteria bacterium | 1,00 | 17,4 | 4,46E-09 | 285 | 194-474 (483) | 67-346 (362) | AlphaFold/UniProt50 (v4) |
| 208 | AF-A0A059GB36-F1-model_v4 | DNA topoisomerase | Hyphomonas oceanitis SCH89 | 1,00 | 16,1 | 4,71E-09 | 312 | 206-472 (483) | 76-323 (329) | AlphaFold/UniProt50 (v4) |
| 209 | AF-A0A554RQ51-F1-model_v4 | DNA topoisomerase IB | Variovorax sp. KBS0712 | 1,00 | 15,6 | 4,71E-09 | 307 | 207-471 (483) | 79-342 (502) | AlphaFold/UniProt50 (v4) |
| 210 | AF-A0A2R3QH91-F1-model_v4 | DNA topoisomerase | Melaminivora sp. SC2-9 | 1,00 | 15,4 | 4,97E-09 | 298 | 206-473 (483) | 78-361 (428) | AlphaFold/UniProt50 (v4) |
| 211 | AF-A0A858X1Q7-F1-model_v4 | DNA topoisomerase IB | Starkeya sp. ORNL1 | 1,00 | 19 | 5,25E-09 | 308 | 209-473 (483) | 75-336 (344) | AlphaFold/UniProt50 (v4) |
| 212 | AF-A0A2S5A3Z3-F1-model_v4 | DNA topoisomerase I | Solitalea longa | 1,00 | 17,4 | 5,25E-09 | 297 | 207-473 (483) | 62-329 (341) | AlphaFold/UniProt50 (v4) |
| 213 | AF-A0A7W7AAC0-F1-model_v4 | DNA topoisomerase-1 | Novosphingobium taihuense | 1,00 | 17,9 | 5,54E-09 | 298 | 208-476 (483) | 16-278 (289) | AlphaFold/UniProt50 (v4) |
| 214 | AF-A0A2W4RQ91-F1-model_v4 | Topoisom_I domain-containing protein | Proteobacteria bacterium | 1,00 | 15,3 | 6,18E-09 | 301 | 209-473 (483) | 74-339 (356) | AlphaFold/UniProt50 (v4) |
| 215 | AF-A0A7V9PN58-F1-model_v4 | DNA topoisomerase IB | Solirubrobacterales bacterium | 1,00 | 16,8 | 6,52E-09 | 311 | 208-483 (483) | 30-309 (319) | AlphaFold/UniProt50 (v4) |
| 216 | AF-A0A2W6CLA9-F1-model_v4 | DNA topoisomerase I | Pseudonocardiales bacterium | 1,00 | 17,6 | 7,67E-09 | 311 | 209-474 (483) | 87-343 (355) | AlphaFold/UniProt50 (v4) |
| 217 | AF-A0A2A9EWP7-F1-model_v4 | DNA topoisomerase IB | Isoptericola jiangsuensis | 1,00 | 15,7 | 7,67E-09 | 307 | 209-475 (483) | 62-324 (327) | AlphaFold/UniProt50 (v4) |
| 218 | AF-A8I5I3-F1-model_v4 | Topoisomerase IB | Azorhizobium caulinodans ORS 571 | 1,00 | 15,9 | 7,67E-09 | 307 | 207-473 (483) | 95-358 (381) | AlphaFold/UniProt50 (v4) |
| 219 | AF-A0A653JE75-F1-model_v4 | DNA topoisomerase (Modular protein) | Burkholderiales bacterium 8X | 1,00 | 18,2 | 7,67E-09 | 297 | 209-475 (483) | 146-427 (481) | AlphaFold/UniProt50 (v4) |
| 220 | AF-A0A853D2W2-F1-model_v4 | DNA topoisomerase IB | Leifsonia shinshuensis | 1,00 | 13,7 | 7,67E-09 | 263 | 149-474 (483) | 256-579 (593) | AlphaFold/UniProt50 (v4) |
| 221 | AF-A0A5Q3HI46-F1-model_v4 | DNA topoisomerase IB | Polaromonas sp. Pch-P | 1,00 | 13,5 | 8,55E-09 | 241 | 156-475 (483) | 35-375 (403) | AlphaFold/UniProt50 (v4) |
| 222 | AF-A0A355ER45-F1-model_v4 | Topoisom_I domain-containing protein | Deltaproteobacteria bacterium | 1,00 | 18 | 9,03E-09 | 295 | 210-448 (483) | 124-347 (372) | AlphaFold/UniProt50 (v4) |
| 223 | AF-A0A2D8BQ10-F1-model_v4 | DNA topoisomerase | Maricaulis sp. | 1,00 | 16,1 | 9,53E-09 | 287 | 194-472 (483) | 62-325 (327) | AlphaFold/UniProt50 (v4) |
| 224 | AF-A0A4R7I138-F1-model_v4 | DNA topoisomerase-1 | Ilumatobacter fluminis | 1,00 | 13,7 | 1,12E-08 | 293 | 207-473 (483) | 71-335 (337) | AlphaFold/UniProt50 (v4) |
| 225 | AF-A0A239L040-F1-model_v4 | DNA topoisomerase-1 | Tropicimonas sediminicola | 1,00 | 15,9 | 1,18E-08 | 314 | 207-446 (483) | 60-281 (315) | AlphaFold/UniProt50 (v4) |
| 226 | AF-A0A1H6UDN6-F1-model_v4 | DNA topoisomerase-1 | Deinococcus reticulitermitis | 1,00 | 14,2 | 1,18E-08 | 261 | 208-475 (483) | 62-332 (342) | AlphaFold/UniProt50 (v4) |
| 227 | AF-A0A3D0CU13-F1-model_v4 | DNA topoisomerase | Chloroflexi bacterium | 1,00 | 16,3 | 1,32E-08 | 300 | 208-474 (483) | 58-329 (351) | AlphaFold/UniProt50 (v4) |
| 228 | AF-A0A519IQL0-F1-model_v4 | DNA topoisomerase IB | Haliea sp. | 1,00 | 14,4 | 1,32E-08 | 270 | 209-475 (483) | 151-453 (470) | AlphaFold/UniProt50 (v4) |
| 229 | AF-A0A3M5MMZ6-F1-model_v4 | Topoisom_I domain-containing protein | Pseudomonas azotoformans | 1,00 | 17,3 | 1,39E-08 | 310 | 213-475 (483) | 4-263 (273) | AlphaFold/UniProt50 (v4) |
| 230 | 3m4a-assembly1_A | Topoisomerase IB | Deinococcus radiodurans | 1,00 | 13,4 | 1,42E-08 | 242 | 209-475 (483) | 34-299 (304) | PDB100 (20240101) |
| 231 | AF-A0A3D5MJ89-F1-model_v4 | DNA topoisomerase | Brevundimonas sp. | 1,00 | 15,4 | 1,47E-08 | 314 | 250-474 (483) | 10-234 (235) | AlphaFold/UniProt50 (v4) |
| 232 | AF-D2PR95-F1-model_v4 | Topoisom_I domain-containing protein | Kribbella flavida DSM 17836 | 1,00 | 13,5 | 1,47E-08 | 234 | 153-474 (483) | 4-330 (349) | AlphaFold/UniProt50 (v4) |
| 233 | AF-A0A323UEJ6-F1-model_v4 | DNA topoisomerase IB | Rhodopseudomonas palustris | 1,00 | 15,6 | 1,64E-08 | 299 | 209-473 (483) | 96-357 (365) | AlphaFold/UniProt50 (v4) |
| 234 | AF-K8P2N2-F1-model_v4 | Topoisom_I domain-containing protein | Afipia clevelandensis ATCC 49720 | 1,00 | 14,7 | 1,73E-08 | 296 | 209-474 (483) | 108-371 (377) | AlphaFold/UniProt50 (v4) |
| 235 | AF-A0A1A9I2L3-F1-model_v4 | Topoisom_I domain-containing protein | Niabella ginsenosidivorans | 1,00 | 15,7 | 1,93E-08 | 300 | 207-475 (483) | 79-341 (347) | AlphaFold/UniProt50 (v4) |
| 236 | AF-A0A2S8V1T6-F1-model_v4 | DNA topoisomerase | Microbacterium sp. MYb43 | 1,00 | 16,2 | 2,15E-08 | 251 | 209-475 (483) | 89-369 (378) | AlphaFold/UniProt50 (v4) |
| 237 | AF-A0A2D6AJ78-F1-model_v4 | Topoisom_I domain-containing protein | Flammeovirgaceae bacterium | 1,00 | 19,2 | 2,27E-08 | 283 | 241-471 (483) | 30-260 (293) | AlphaFold/UniProt50 (v4) |
| 238 | AF-A0A1G9TFZ0-F1-model_v4 | DNA topoisomerase-1 | Oryzisolibacter propanilivorax | 1,00 | 15,7 | 2,53E-08 | 275 | 209-473 (483) | 78-358 (430) | AlphaFold/UniProt50 (v4) |
| 239 | AF-A0A2C6CRF3-F1-model_v4 | DNA topoisomerase I | Lewinellaceae bacterium SD302 | 1,00 | 14,7 | 2,53E-08 | 247 | 207-474 (483) | 74-346 (349) | AlphaFold/UniProt50 (v4) |
| 240 | 2f4q-assembly1_A | topoisomerase IB | Deinococcus radiodurans | 1,00 | 14,5 | 2,72E-08 | 222 | 209-475 (483) | 45-305 (309) | PDB100 (20240101) |
| 241 | AF-A0A366ILE6-F1-model_v4 | DNA topoisomerase-1 | Brevibacterium celere | 1,00 | 16,4 | 2,98E-08 | 251 | 209-476 (483) | 63-300 (300) | AlphaFold/UniProt50 (v4) |
| 242 | AF-A0A7W0P0D4-F1-model_v4 | DNA topoisomerase IB | Acidimicrobiia bacterium | 1,00 | 15,6 | 3,14E-08 | 280 | 207-475 (483) | 69-333 (335) | AlphaFold/UniProt50 (v4) |
| 243 | AF-A0A7I7KRT4-F1-model_v4 | Topoisom_I domain-containing protein | Mycobacterium cookii | 1,00 | 18,4 | 3,70E-08 | 285 | 269-475 (483) | 6-216 (225) | AlphaFold/UniProt50 (v4) |
| 244 | AF-A0A2D8M293-F1-model_v4 | Topoisom_I domain-containing protein | Micavibrio sp. | 1,00 | 15,5 | 4,12E-08 | 249 | 209-476 (483) | 77-349 (361) | AlphaFold/UniProt50 (v4) |
| 245 | AF-A0A2P7BK40-F1-model_v4 | DNA topoisomerase I | Phyllobacterium sophorae | 1,00 | 21,8 | 4,59E-08 | 282 | 217-475 (483) | 4-254 (261) | AlphaFold/UniProt50 (v4) |
| 246 | AF-A0A4Q5W5R9-F1-model_v4 | DNA topoisomerase IB | Comamonadaceae bacterium | 1,00 | 18,5 | 4,85E-08 | 294 | 209-429 (483) | 53-276 (280) | AlphaFold/UniProt50 (v4) |
| 247 | AF-A0A3D2NF76-F1-model_v4 | Topoisomerase I | Microbacterium sp. | 1,00 | 20,2 | 5,12E-08 | 297 | 209-407 (483) | 63-258 (258) | AlphaFold/UniProt50 (v4) |
| 248 | AF-A0A344WF02-F1-model_v4 | Topoisom_I domain-containing protein | Hyphomonas sp. CACIAM 19H1 | 1,00 | 22,4 | 5,40E-08 | 269 | 243-473 (483) | 11-225 (227) | AlphaFold/UniProt50 (v4) |
| 249 | AF-A0A0Q0AWQ7-F1-model_v4 | DNA topoisomerase | Pseudomonas syringae pv. syringae | 1,00 | 21,8 | 5,70E-08 | 308 | 209-394 (483) | 73-255 (257) | AlphaFold/UniProt50 (v4) |
| 250 | AF-A0A4R3YX53-F1-model_v4 | DNA topoisomerase I-like protein | Luteibacter rhizovicinus | 1,00 | 16,3 | 7,90E-08 | 290 | 248-475 (483) | 4-232 (239) | AlphaFold/UniProt50 (v4) |

**Supplementary table 2. Bacterial proteins having structural similarities with human TOP1 as identified by Foldseek.**

| No. | Target | Description | Scientific name | Prob. | Seq. id. | E-value | Score | Query pos. | Target pos. | Database |
|---|---|---|---|---|---|---|---|---|---|---|
| 251 | AF-A0A2D5MD62-F1-model_v4 | Topoisom_I domain-containing protein | Planctomycetaceae bacterium | 1,00 | 18,6 | 7,90E-08 | 281 | 207-446 (483) | 44-267 (272) | AlphaFold/UniProt50 (v4) |
| 252 | AF-A0A4V3I7Y8-F1-model_v4 | DNA topoisomerase IB | Cryobacterium sp. HLT2-23 | 1,00 | 19,7 | 9,81E-08 | 263 | 207-436 (483) | 61-279 (314) | AlphaFold/UniProt50 (v4) |
| 253 | AF-A0A5B8LUT8-F1-model_v4 | DNA topoisomerase IB | Devosia ginsengisoli | 1,00 | 18,7 | 1,09E-07 | 283 | 207-459 (483) | 60-306 (364) | AlphaFold/UniProt50 (v4) |
| 254 | AF-A0A538SHC1-F1-model_v4 | DNA topoisomerase IB | Candidatus Eisenbacteria bacterium | 1,00 | 15,4 | 1,15E-07 | 281 | 208-434 (483) | 158-394 (399) | AlphaFold/UniProt50 (v4) |
| 255 | AF-A0A356W4H9-F1-model_v4 | Topoisom_I domain-containing protein | Hyphomonas atlantica | 1,00 | 21,1 | 1,51E-07 | 260 | 272-475 (483) | 10-195 (199) | AlphaFold/UniProt50 (v4) |
| 256 | AF-A0A553KX14-F1-model_v4 | DNA topoisomerase IB | Aliiglaciecola sp. M165 | 1,00 | 15,3 | 1,60E-07 | 252 | 208-478 (483) | 61-334 (337) | AlphaFold/UniProt50 (v4) |
| 257 | AF-A0A538F763-F1-model_v4 | DNA topoisomerase IB | Actinomycetia bacterium | 1,00 | 19,3 | 2,10E-07 | 275 | 207-409 (483) | 61-262 (262) | AlphaFold/UniProt50 (v4) |
| 258 | AF-A0A346XZ00-F1-model_v4 | Topoisom_I domain-containing protein | Euzebya pacifica | 1,00 | 14,1 | 2,60E-07 | 240 | 207-477 (483) | 59-321 (338) | AlphaFold/UniProt50 (v4) |
| 259 | AF-A0A6P1AZT3-F1-model_v4 | TOPEUc domain-containing protein | bacterium LRH843 | 1,00 | 76,1 | 8,59E-07 | 260 | 335-422 (483) | 1-88 (89) | AlphaFold/UniProt50 (v4) |
| 260 | AF-A0A7C7QNS7-F1-model_v4 | DNA topoisomerase I | Anaerolineae bacterium | 1,00 | 28,6 | 1,33E-06 | 260 | 16-166 (483) | 2-151 (151) | AlphaFold/UniProt50 (v4) |
| 261 | AF-A0A7V9UEA0-F1-model_v4 | DNA topoisomerase IB | Actinomycetia bacterium | 1,00 | 17,9 | 1,65E-06 | 219 | 272-472 (483) | 3-211 (228) | AlphaFold/UniProt50 (v4) |
| 262 | AF-A0A0S8KK16-F1-model_v4 | TOPEUc domain-containing protein | Anaerolineae bacterium SM23_84 | 1,00 | 21,2 | 1,83E-06 | 212 | 299-475 (483) | 1-296 (310) | AlphaFold/UniProt50 (v4) |
| 263 | AF-A0A3C2E1M7-F1-model_v4 | DNA topoisomerase | Alphaproteobacteria bacterium | 1,00 | 18,5 | 4,61E-06 | 238 | 194-394 (483) | 62-255 (262) | AlphaFold/UniProt50 (v4) |
| 264 | AF-A0A7R7Y4N0-F1-model_v4 | Topoisom_I domain-containing protein | Luteitalea sp. TBR-22 | 1,00 | 17,3 | 6,05E-06 | 244 | 209-387 (483) | 93-268 (343) | AlphaFold/UniProt50 (v4) |
| 265 | AF-A0A175RIY7-F1-model_v4 | DNA topoisomerase | Curtobacterium luteum | 1,00 | 20,4 | 1,99E-05 | 206 | 204-362 (483) | 63-208 (209) | AlphaFold/UniProt50 (v4) |
| 266 | AF-Q5FR70-F1-model_v4 | DNA topoisomerase I | Gluconobacter oxydans 621H | 1,00 | 16,9 | 2,35E-05 | 213 | 207-363 (483) | 77-220 (242) | AlphaFold/UniProt50 (v4) |
| 267 | AF-A0A3D4ER35-F1-model_v4 | DNA topoisomerase I | Leeuwenhoekiella sp. | 1,00 | 17,5 | 4,75E-05 | 184 | 299-483 (483) | 11-199 (201) | AlphaFold/UniProt50 (v4) |
| 268 | AF-A0A3C1NIH3-F1-model_v4 | Topoisom_I domain-containing protein | Hyphomicrobiales bacterium | 1,00 | 14,5 | 9,61E-05 | 197 | 208-364 (483) | 72-215 (215) | AlphaFold/UniProt50 (v4) |
| 269 | AF-A0A4Q4D6Q1-F1-model_v4 | DNA topoisomerase IB | Actinomycetales bacterium | 1,00 | 17,5 | 1,33E-04 | 182 | 208-364 (483) | 45-189 (189) | AlphaFold/UniProt50 (v4) |
| 270 | AF-A0A354YES1-F1-model_v4 | DNA topoisomerase | Xanthomonadaceae bacterium | 1,00 | 20,4 | 1,57E-04 | 165 | 303-472 (483) | 2-172 (185) | AlphaFold/UniProt50 (v4) |
| 271 | 3m4aA03 | Topoisomerase 1 | Deinococcus radiodurans R1 = ATCC 13939 = DSM 20539 | 1,00 | 17,8 | 2,05E-04 | 171 | 269-382 (483) | 6-118 (118) | CATH50 (v4.3.0) |
| 272 | AF-A0A845QK30-F1-model_v4 | Uncharacterized protein | Anaerotruncus colihominis | 1,00 | 18,3 | 3,53E-04 | 162 | 259-437 (483) | 110-255 (264) | AlphaFold/UniProt50 (v4) |
| 273 | AF-A0A542SXY1-F1-model_v4 | DNA topoisomerase I-like protein | Streptomyces puniciscabiei | 1,00 | 16,3 | 4,89E-04 | 161 | 232-407 (483) | 30-196 (277) | AlphaFold/UniProt50 (v4) |

No. = number, prob. = probability of target being homologous to human TOP1, seq. id. = sequence identity, E-value = expect value, score = structural bit score, pos. = position.

**Supplementary table 3. Viral proteins having structural similarities with human TOP1 as identified by Foldseek.**

| No. | Target | Description | Scientific Name | Prob. | Seq. Id. | E-Value | Score | Query Pos. | Target Pos. | Database |
|---|---|---|---|---|---|---|---|---|---|---|
| 1 | 2h7g-assembly1_X | topoisomerase | Variola virus | 1,00 | 17,2 | 1,31E-09 | 309 | 209-475 (483) | 56-306 (312) | PDB100 (20240101) |
| 2 | 2h7gX02 | Topoisomerase 1 | Variola virus human/India/Ind3/1967 | 1,00 | 18,4 | 2,15E-08 | 284 | 236-475 (483) | 6-233 (239) | CATH50 (v4.3.0) |
| 3 | 1a41-assembly1_A | Type 1-Topoisomerase Catalytic Fragment | Vaccinia virus | 1,00 | 17,1 | 1,06E-05 | 169 | 236-477 (483) | 2-207 (221) | PDB100 (20240101) |

No. = number, prob. = probability of target being homologous to human TOP1, seq. id. = sequence identity, E-value = expect value, score = structural bit score, pos. = position.

**Supplementary Table 4. B cell receptor sequence details of mAbs.**

| Clone | Antigen | HC | IGHV | IGHV identity (%) | IGHD | IGHJ | IGH-CDR3 | LC | IGK/LV | IGK/LV identity (%) | IGK/LJ | IGK/L-CDR3 |
|---|---|---|---|---|---|---|---|---|---|---|---|---|
| 2F8 | H. TOP1 | IGHA1 | V1-18 | 95.14 | D6-6 | J4 | CARGRDFSSSPYDYW | λ | V3-21 | 97.49 | J2/J3 | CQVWDRSSDDVIF |
| 7G6 | H. TOP1 | IGHG1 | V3-48 | 90.97 | D6-6 | J5 | CATTPRTGRPW | κ | V4-1 | 95.96 | J4 | CQQYLTPLQIF |
| 9D11 | H. TOP1 | IGHG3 | V1-3 | 93.75 | D2-15 | J4 | CARDGGSSVAVISATLDSW | λ | V2-23 | 97.22 | J2/J3 | CCSYAGSRGVVF |
| 2C11 | H. TOP1 | IGHG1 | V3-30 | 96.88 | D5-18 | J4 | CAKDGSDVRRGYIYGSFDYW | κ | V1-33 | 97.77 | J4 | CQQYANLPLTF |
| 6E3 | H. TOP1 | IGHG1 | V3-23 | 89.58 | D2-2 | J4 | CAKQRCEATGCPSDFW | λ | V3-21 | 95.34 | J2/J3 | CQVWDRYSDTIFF |
| 7D11 | H. TOP1 | IGHG1 | V1-18 | 83.68 | D3-16 | J4 | CARGGDSSSAASDYW | λ | V3-21 | 94.27 | J2/J3 | CQVWDNSDDSVLF |
| 8D7 | H. TOP1 | IGHG1 | V3-48 | 89.24 | D1-14 | J5 | CATTPKAGRPW | κ | V4-1 | 96.63 | J4 | CQQYYTTFQTF |
| D2 | TT | IGHG1 | V4-39 | 92.44 | D3-16 | J4 | CARHVLITFGGLIEQTYYFDSW | κ | V2-28 | 96.60 | J5 | CMGGLQTPTF |
| 3F3 | CCP2 | IGHG2/4 | V1-2 | 80.56 | D2-8 | J3 | CARGTYLPVDESAAFDVW | κ | V4-1 | 87.88 | J2 | CQQYYEAPYTF |

Antigen = antigen used for isolation of the B cell, H. TOP1 = human TOP1, TT = tetanus toxoid, CCP2 = cyclic citrullinated peptide 2, HC = heavy chain, IGHV = heavy chain V gene, IGHV identity (%) = similarity of IGHV compared to closest germline IGHV expressed in a percentage, IGHD = heavy chain D gene, IGHJ = heavy chain J gene, IGH-CDR3 = amino acid sequence of heavy chain complementarity-determining region 3, LC = light chain, κ = kappa, λ = lambda, IGK/LV = light chain V gene, IGK/LV identity (%) = similarity of IGK/LV compared to germline IGK/LV expressed in a percentage, IGK/LJ = light chain J gene, IGK/L-CDR3 = amino acid sequence of light chain complementarity-determining region 3.
